# Total synthesis and structural characterization of a novel protein scaffold from the snail *Biomphalaria glabrata*

**DOI:** 10.64898/2026.04.03.712544

**Authors:** Oleg Melnyk, Stéphanie Caby, Armelle Vigouroux, Christine Demanche, Rémi Desmet, Magalie Sénéchal, Benoît Snella, Alexandra Mougel, Céline Boidin-Wichlacz, Aurélie Parmentier, Ugo Pasco, Sonia Cantel, Solange Moréra, Jérôme Vicogne

## Abstract

Disulfide-rich miniproteins constitute compact and highly stable scaffolds of growing interest for molecular and structural engineering. Schistosomins are ∼80-residue proteins conserved across gastropods that form a long-standing orphan family whose structure and biological roles have remained unknown. Here, we report the total chemical synthesis and structural characterization of a schistosomin isoform from *Biomphalaria glabrata*, a medically relevant intermediate host of the parasite *Schistosoma mansoni*. Using state-of-the-art solid-phase peptide synthesis, chemoselective peptide ligation, and controlled oxidative folding, we obtained homogeneous well folded schistosomin suitable for biophysical and structural studies. High-resolution X-ray crystallography reveals a previously undescribed disulfide-rich fold defining a new class of miniprotein scaffold. Nano differential scanning fluorimetry and circular dichroism experiments demonstrate the remarkable thermal stability of this scaffold, while molecular dynamics simulations confirm the intrinsic rigidity of its disulfide-stabilized core and show that the two naturally occurring isoforms differing by a single residue exhibit nearly indistinguishable structural and dynamic properties. Finally, transcript and protein analyses across snail tissues provide the first spatial expression map of schistosomin in a medically relevant mollusk. Together, this work establishes schistosomin as a novel and robust miniprotein scaffold and provides a structural and biological framework for exploring its function and potential applications.

## Introduction

Miniproteins are small polypeptides, typically below 10 kDa, that adopt well-defined tertiary structures. Owing to their compact size and constrained architecture, they have emerged as attractive molecular frameworks for both functional studies and therapeutic development (1). Their structural rigidity often confers exceptional stability and binding specificity, positioning them at the interface between small molecules and large biologics. Whereas small molecules diffuse efficiently but often struggle to engage extended protein-protein interfaces, antibodies provide exquisite selectivity at the cost of limited tissue penetration and complex manufacturing. Miniproteins can, in principle, combine some of the advantages of both classes (2–5).

This growing interest has been fueled by advances in computational design (6, 7), delivery strategies (4) and, importantly, modern solid-phase peptide synthesis (SPPS) (8), which in certain cases enables the synthesis of proteins comprising up to ∼100 residues (9). In parallel, the development of chemoselective peptide-ligation methods has profoundly expanded the scope of chemical protein synthesis by allowing the selective coupling of unprotected peptide segments (10–17). In synergy with SPPS, these approaches enable the modular assembly of proteins bearing complex modifications, non-canonical residues, or tailored disulfide patterns—features that often remain difficult or inaccessible through recombinant expression (18).

Natural ecosystems constitute a vast and still underexplored reservoir of compact, highly stabilized miniproteins. Venomous organisms such as snakes (19), spiders (20), and cone snails (21–23) have yielded numerous bioactive scaffolds of pharmaceutical relevance. In contrast, miniproteins from non-venomous mollusks remain comparatively understudied, despite the long-standing medicinal use of terrestrial and aquatic snails (24). Recent reports describing antimicrobial peptides from snail mucus further suggest that these organisms may harbor a broader and largely overlooked diversity of biologically active disulfide-rich miniproteins (25).

Among these, schistosomins form a small family of ∼80-residue, cysteine-rich proteins initially identified in the freshwater snail *Lymnaea stagnalis* during infection with the avian trematode *Trichobilharzia ocellata* (26, 27). Homologous sequences have since been reported in multiple gastropod species. Early studies described schistosomin as a neuroendocrine factor capable of antagonizing reproductive hormones in *L. stagnalis*, (28, 29) and highlighted its remarkable biochemical robustness, including preserved biological activity after heating at 100°C. Two schistosomin homologues were later identified in the freshwater snail *Biomphalaria glabrata*, the intermediate host of the human parasite *Schistosoma mansoni*. These two isoforms are encoded by two distinct genes but only differ by a single amino-acid substitution (A78P) within the mature form. In *B. glabrata*, however, schistosomins expression does not vary upon parasite infection, suggesting physiological roles beyond parasite-induced castration (30).

Despite more than three decades since their initial discovery, no experimental structural information has been available for any schistosomin, and their three-dimensional architecture has remained entirely speculative. The unusual combination of small size, multiple disulfide bonds, and the absence of close homologs in protein structure databases suggested that schistosomins might adopt an as-yet undescribed type of miniprotein fold. At the same time, the strict conservation of their cysteine framework across distantly related gastropod species raised the possibility that schistosomins could belong to a broader family of disulfide-rich molluskan scaffolds whose structural features are encoded primarily by cysteine topology rather than primary sequence conservation. However, progress toward addressing these questions has been hampered by the low abundance of native material and the technical challenges associated with producing correctly folded protein using recombinant expression systems.

Here, we combine chemical synthesis, crystallography, biophysics approaches, and molecular simulations to characterize the structure and biochemical properties of a schistosomin isoform from *B. glabrata*. To overcome the limitations of the recombinant expression of this cysteine-rich protein, we implemented a fully synthetic strategy based on solid-phase peptide synthesis, chemoselective ligation of unprotected segments followed by controlled oxidative folding. This approach provided homogeneous material allowing (i) its structure determination showing a new miniprotein fold and (ii) thermal stability studies using nano differential scanning calorimetry (nanoDSC) and circular dichroism (CD). Molecular dynamics (MD) simulations were used to evaluate the rigidity or flexibility of the two isoforms scaffold (alanine versus proline at position 78). We also examined schistosomin transcript and protein distribution across *B. glabrata* tissues, providing the first spatial expression map for this protein family in a medically relevant mollusk. These analyses reveal that schistosomin expression is not restricted to neuronal tissues and support a secreted, systemic distribution, challenging the view of schistosomin as a strictly neuropeptide-like factor. Finally, structural comparisons between AlphaFold models and our structures indicate that schistosomin belongs to a broader family of disulfide-rich molluskan miniproteins, including sequences tentatively classified among uncharacterized conotoxin-like peptides (31), whose three-dimensional architecture converges despite pronounced sequence divergence.

## Results and discussion

### 1. Identification and characterization of predicted native schistosomin isoforms from *Biomphalaria glabrata*

*Biomphalaria glabrata* is readily maintained in fresh water at a temperature of 28°C under standardized lab conditions. The obtention of adult snails from the egg stage requires 3-4 months and its high fecundity provides sufficient biological material for biochemical analyses (Figure 1A). Genomic and transcriptomic data predict the expression of two schistosomin isoforms in this species (30) (Figure 1B). These two isoforms differ only by a single amino-acid substitution (Ala/Pro) at position 78. The two mature isoforms, which were identified in *B. glabrata* protein extracts, start at Asp18 following signal peptide cleavage as predicted, consistent with a secreted protein as observed with the *L. stagnalis* schistosomin.

**Figure 1.**
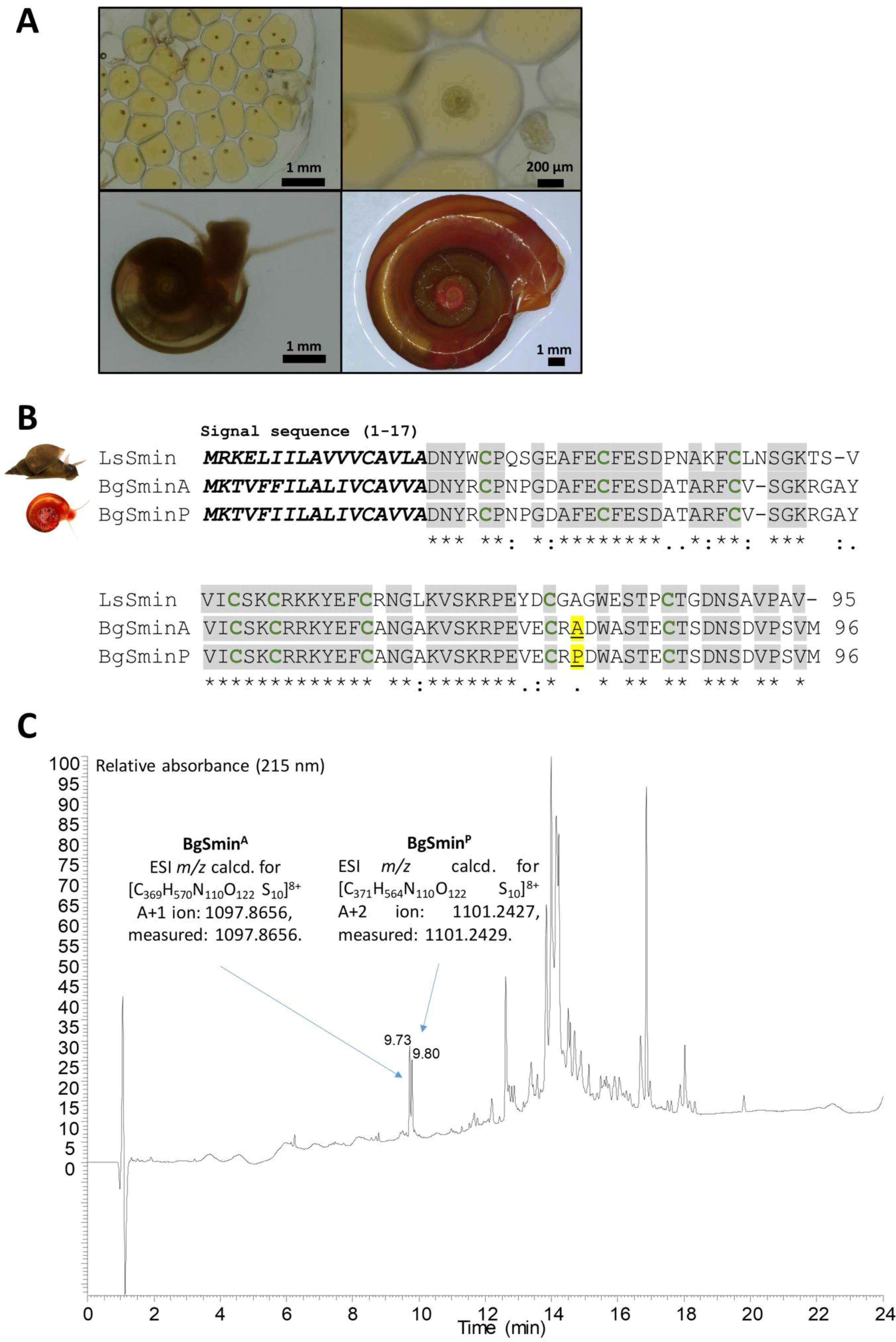
Identification of native schistosomin isoforms in *Biomphalaria glabrata*. A. Developmental stages of *B. glabrata* providing a reproducible source of biological material for protein extraction. Freshly laid egg mass (top left), ∼6-day embryonated egg (top right), 15-day juvenile snail (bottom left), and 3-month adult snail (bottom right). B. Sequence alignment of schistosomin from *B. glabrata* (BgSmin^A^ and BgSmin^P^; GenBank accession numbers EU126802 and ES488206) with schistosomin from *Lymnaea stagnalis* (LsSmin; AAB20290), the first schistosomin identified. Conserved residues are indicated by an asterisk (*), strongly similar residues by a colon (:), and weakly similar residues by a period (.). The two *B. glabrata* isoforms differ by a single amino-acid substitution at position 78 (Ala/Pro) within the mature sequence (underlined and highlighted in yellow) C. UPLC–MS analysis of protein extracts from adult *B. glabrata* showing two chromatographic peaks corresponding to BgSmin^A^ (9.73 min) and BgSmin^P^ (9.80 min). Column: ACQUITY UPLC Peptide BEH C18, 300 Å, 1.7 µm, 2.1 × 150 mm. Gradient: 0-70% B in 15 min. Eluents: water with 0.1% (v/v) TFA; acetonitrile with 0.1% (v/v) TFA. Flow rate: 0.4 mL min⁻¹. Column temperature: 70°C.

Native schistosomins were isolated as two individual chromatographic peaks by reversed-phase HPLC (RP-HPLC) from protein extracts prepared from dissected foot tissues of adult snails following ammonium sulfate precipitation (see Suppl. Info.). High-resolution mass spectrometry (HRMS) analyses matched precisely the theoretical masses for the expected mature sequences minus 8 mass units, consistent with two distinct forms stabilized by four intramolecular disulfide bonds (Figure 1C, Suppl. Info.). The two proteins were detected in comparable amounts in the extracts and are hereafter referred to as BgSmin^A^ and BgSmin^P^.

Although these analyses establish the identity of the two native schistosomin isoforms, the limited amount of material obtainable from snail tissues precludes extensive biophysical and structural characterization, therefore motivating the implementation of a total chemical synthesis strategy to obtain sufficient quantities of homogeneous material for detailed analysis.

### 2. Total chemical synthesis of schistosomin enables access to homogeneous and correctly folded protein

Previous attempts to produce BgSmin^A^ isoform using classical recombinant expression strategies resulted in insoluble material (29, 30)_,_ most likely due to incorrect disulfide bond formation. In our hands, we also obtained a heterogeneous product difficult to purify. These limitations led us to implement a total chemical synthesis strategy to obtain sufficient quantities of homogeneous material with full control over folding.

The first attempts to chemically produce BgSmin^A^ isoform revealed that the Asp88-Asn89 diad was prone to aspartimide formation. To avoid this side reaction, Asp88 was substituted by Glu. In addition, a C-terminal lysine (Lys97) bearing a biotin moiety was introduced to facilitate future functional studies without interfering with the structural core. The synthetic approach relied on the assembly in solution of three peptide segments produced by SPPS (See details in Suppl. Info.). The full-length linear precursor was obtained in only two steps through a redox-controlled sequential bis(2-sulfanylethyl)amido (SEA)-mediated peptide ligation/native chemical ligation strategy (residues 18-96; Figure 2A). The purified linear precursor was used to generate polyclonal antibodies that proved highly specific and enabled subsequent detection of native schistosomin in tissue extracts.

**Figure 2.**
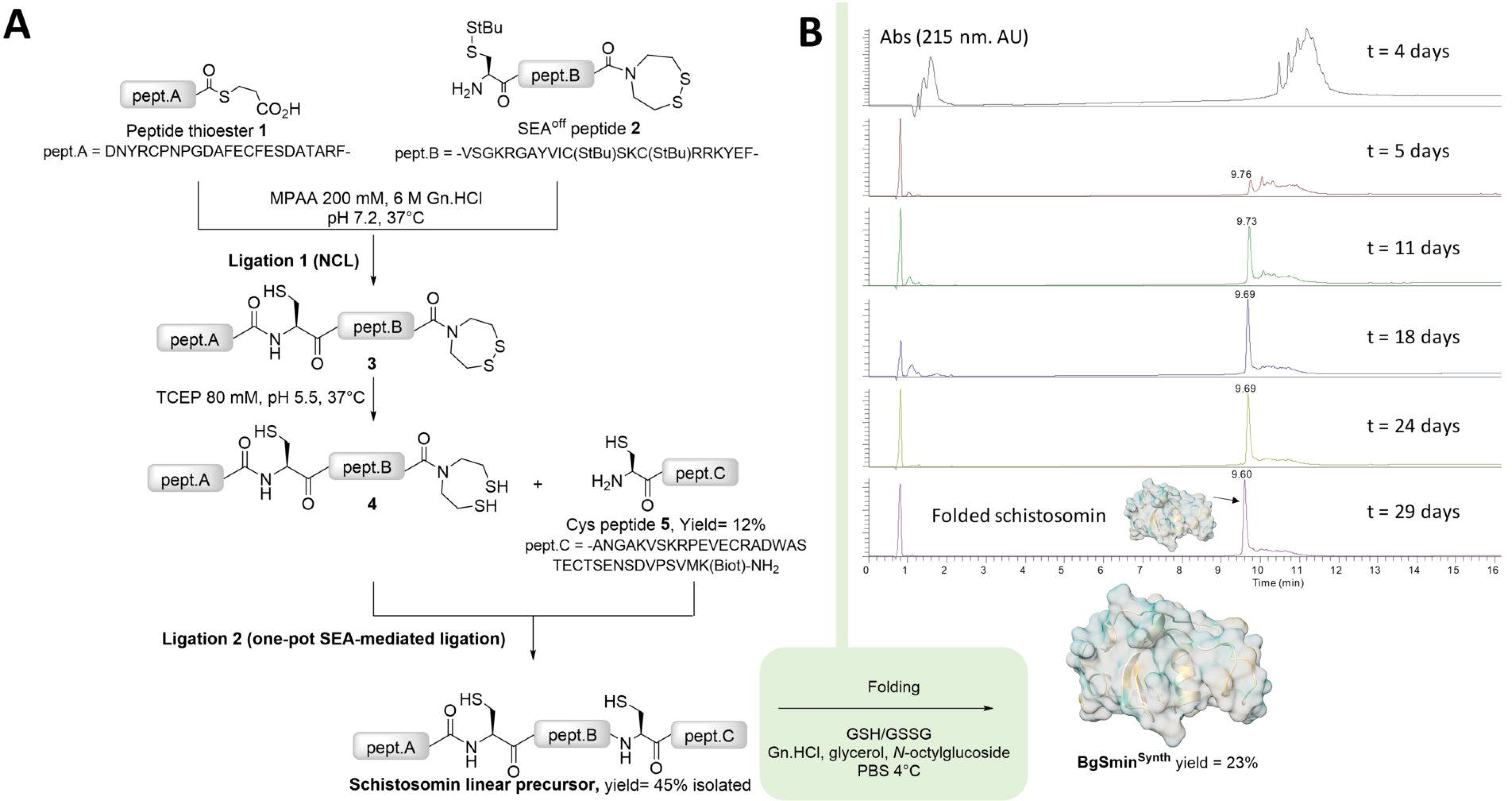
Total chemical synthesis and oxidative folding of BgSmin^Synth^. A: Design and workflow of the chemical synthesis of BgSmin^Synth^. The mature schistosomin sequence (residues 18-96) was assembled from three peptide segments (B, C and D) produced by solid-phase peptide synthesis (SPPS). The first ligation was achieved by native chemical ligation (NCL) between segments B and C. The resulting intermediate was then subjected to reductive activation of the SEA group to generate an *in situ* thioester, enabling a second native chemical ligation with segment D to afford the full-length linear precursor. To prevent aspartimide formation during synthesis, Asp88 was substituted by Glu. In addition, a C-terminal lysine bearing a biotin moiety (Lys97) was introduced to facilitate downstream functional studies (See details and analysis in Suppl. Info.). B: Monitoring of oxidative folding of the synthetic BgSmin precursor by analytical UPLC. The unfolded linear precursor was first solubilized in 6 M guanidinium hydrochloride and diluted into a glutathione-based redox buffer supplemented with glycerol and the non-ionic detergent *n*-octylglucoside. Folding was carried out at 4°C and progressively converged over 29 days toward a single dominant species corresponding to the correctly folded BgSminSynth protein.

Folding of the synthetic schistosomin required carefully optimized oxidative conditions due to limited solubility and aggregation tendencies. A robust protocol was established by first solubilizing the linear precursor in 6 M guanidinium hydrochloride (Gn.HCl) and adding this solution to a glutathione-based redox buffer supplemented with glycerol and non-ionic detergent *n*-octylglucoside. The folding was performed at low temperature (4°C), which proved to be also critical for minimizing protein aggregation. Although the oxidative folding process proceeded slowly (up to 4 weeks), it reproducibly converged toward a single dominant species, consistent with a well-defined and thermodynamically favored disulfide connectivity (Figure 2B). HRMS analysis confirmed the formation of the fully oxidized protein with the expected molecular mass (See Suppl. Info.).

The availability of milligram quantities of homogeneous synthetic protein (BgSmin^Synth^), together with a robust folding protocol, provided the basis for detailed structural and biophysical investigations

### 3. Crystal structures of BgSmin protein reveal a novel disulfide-rich fold

The same crystallization condition (Table 1) of BgSmin^Synth^ protein (residues 18-96) led to two different space groups (P2₁ and C2), reflecting distinct packing arrangements that involve the additional C-terminal biotinylated lysine residue at position 97. In both crystal forms (PDB entries 9RT6 and 9FDO; Table 1), two molecules are present in the asymmetric unit, yielding four independent and identical molecules, as indicated by the average root mean square deviation (RMSD) of 0.55 Å for all Cα atoms. In line with the very low sequence identity of BgSmin (below 10%) against the Protein Data Bank members, no homologous model could be used as a search model for molecular replacement. A reliable predicted model generated using AlphaFold2 allowed structure determination.

**Table 1.** Crystallographic data and refinement parameters for BgSmin^Synth^

| PDB code | 9RT6 (9FDM) | 9FDO |
| --- | --- | --- |
| Crystallization conditions | 2 M AS/ 0.1 M Tris-HCl pH 8/ 0.2 M NaBr |  |
| Space group | P2<sub>1</sub> | C2 |
| Cell parameters (Å, °) | a= 21.8<br>b= 38.4<br>c= 87.8<br>β= 91.2 | a= 87.7<br>b= 38.0<br>c= 43.2<br>β= 95.3 |
| Resolution (Å) | 19.7-1.9<br>(2-1.9) | 43.6-2.07<br>(2.17-2.07) |
| Estimated resolution limit (Å)<br>STARANISO | 1.80; 1.98 ; 1.94 | 2.12; 2.06; 2.05 |
| No. of observed reflections | 77793 (3325) | 43741 (856) |
| No. of unique reflections | 10239 (512) | 7585 (379) |
| Completeness (%) | 87.8 (30.3) | 84.5 (29.4) |
| R<sub>merge</sub> (%) | 26 (133.6) | 13.3 (115.8) |
| R<sub>pim</sub> (%) | 10 (56.6) | 5.8 (94.6) |
| I/σ(I) | 5.6 (1.4) | 7.7 (0.8) |
| CC<sub>1/2</sub> | 0.985 (0.483) | 0.994 (0.313) |
| R<sub>work</sub> (%) | 20.7 | 22.9 |
| R<sub>free</sub> (%) | 25 | 27.5 |
| rms bond deviation (Å) | 0.008 | 0.008 |
| rms angle deviation (°) | 0.96 | 0.93 |
| Average B (Å<sup>2</sup>) |  |  |
| Protein A/B | 18.7/27 | 33.5/35.5 |
| solvent | 25.7 | 35.2 |
| <sup>a</sup>Clashscore | 1.68 | 0.82 |
| MolProbity score | 1.46 | 1.51 |
| <sup>a</sup>Ramachandran plot (%) |  |  |
| Favoured | 94.00 | 94.16 |
| Outliers | 1.33 | 0.65 |

Values for the highest resolution shell are in parentheses. CC<sub>1/2</sub>= percentage of correlation between intensities from random half-dataset. <sup>a</sup>Calculated with MolProbity
STARANISO applies ellipsoidal mask: estimated resolution limits along the three crystallographic directions a\*, b\*, c\*.

BgSmin^Synth^ adopts a compact monomeric fold stabilized by four intramolecular disulfide bridges (Cys22-Cys62, Cys31-Cys55, Cys41-Cys76, and Cys52-Cys85; Figure 4), corresponding to a 1-6, 2-5, 3-7, and 4-8 connectivity pattern with no equivalent in the Protein Data Bank (PDB). The first two disulfide bonds tightly anchor the N-terminal region (residues 18-38) to the central α-helix spanning residues 54-64, while the latter two constrain the β-strand elements preceding the helix (residues 39-44 and 47-52) against the C-terminal region. Such a topology is expected to contribute to the intrinsic stability of the scaffold, in agreement with previous reports describing the resistance of native schistosomin to thermal treatment in *L. stagnalis* (29). As classically observed in protein structures, the two-three-first N-terminal residues (Asp18, Asn19, and Tyr20), are poorly defined or not visible in the electron density maps in the four molecules suggesting conformational flexibility in this region. Consistently, molecular dynamics simulations of BgSmin isoforms in explicit water at 300 K show elevated root-mean-square fluctuations (RMSF) for these residues (see Suppl. Info.).

**Figure 3:**
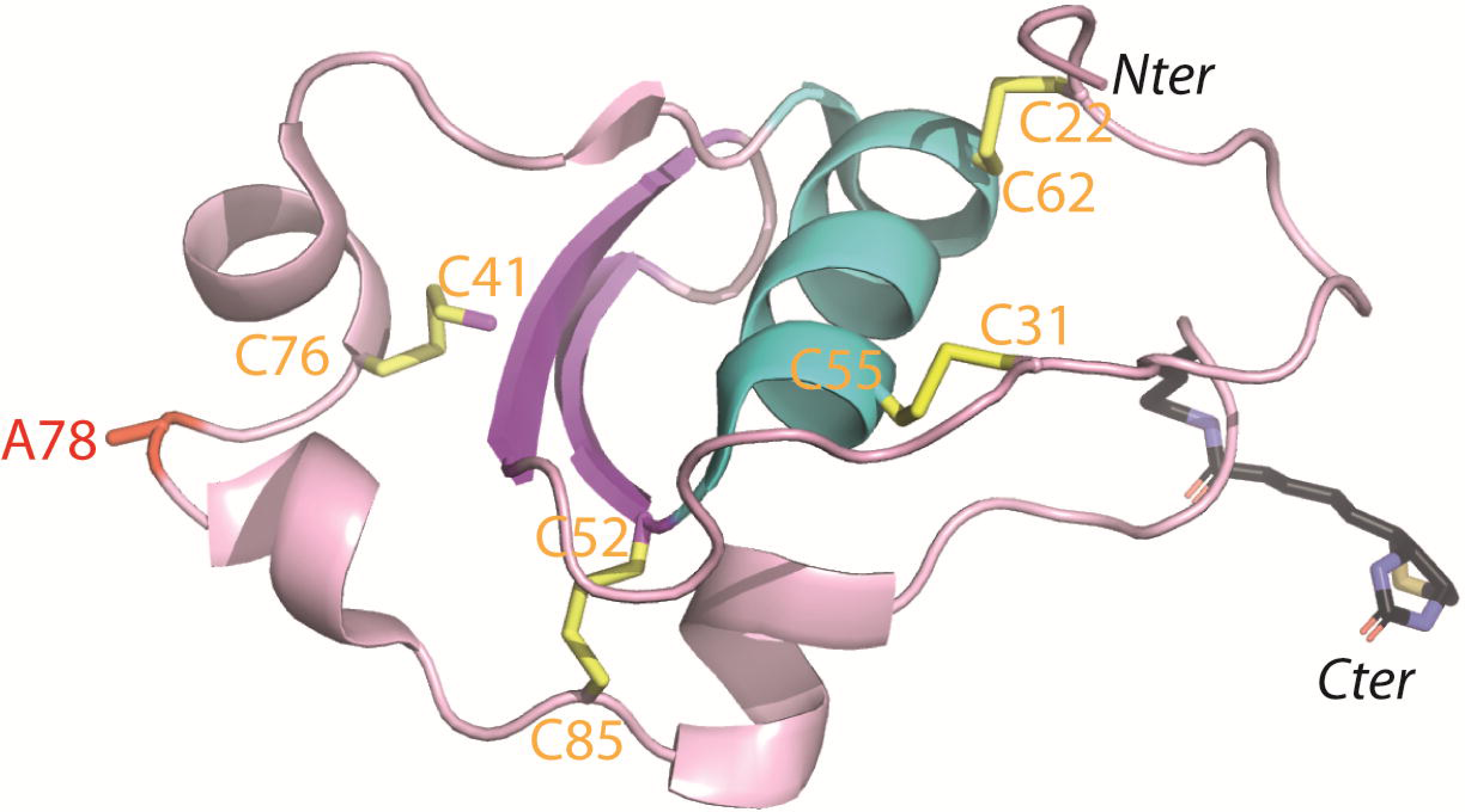
Structure of BgSmin^Synth^ Ribbon representation of BgSmin^Synth^ (PDB 9FDO) in pink with the central α-helix in cyan and two β-strands in purple. The four intramolecular disulfide bridges are shown in yellow with the cysteine residues labelled. Both the Nter and Cter are labeled. The C-terminal biotinylated lysine is indicated with its side chain and shown in grey. Ala78 side chain is shown in red.

**Figure 4.**
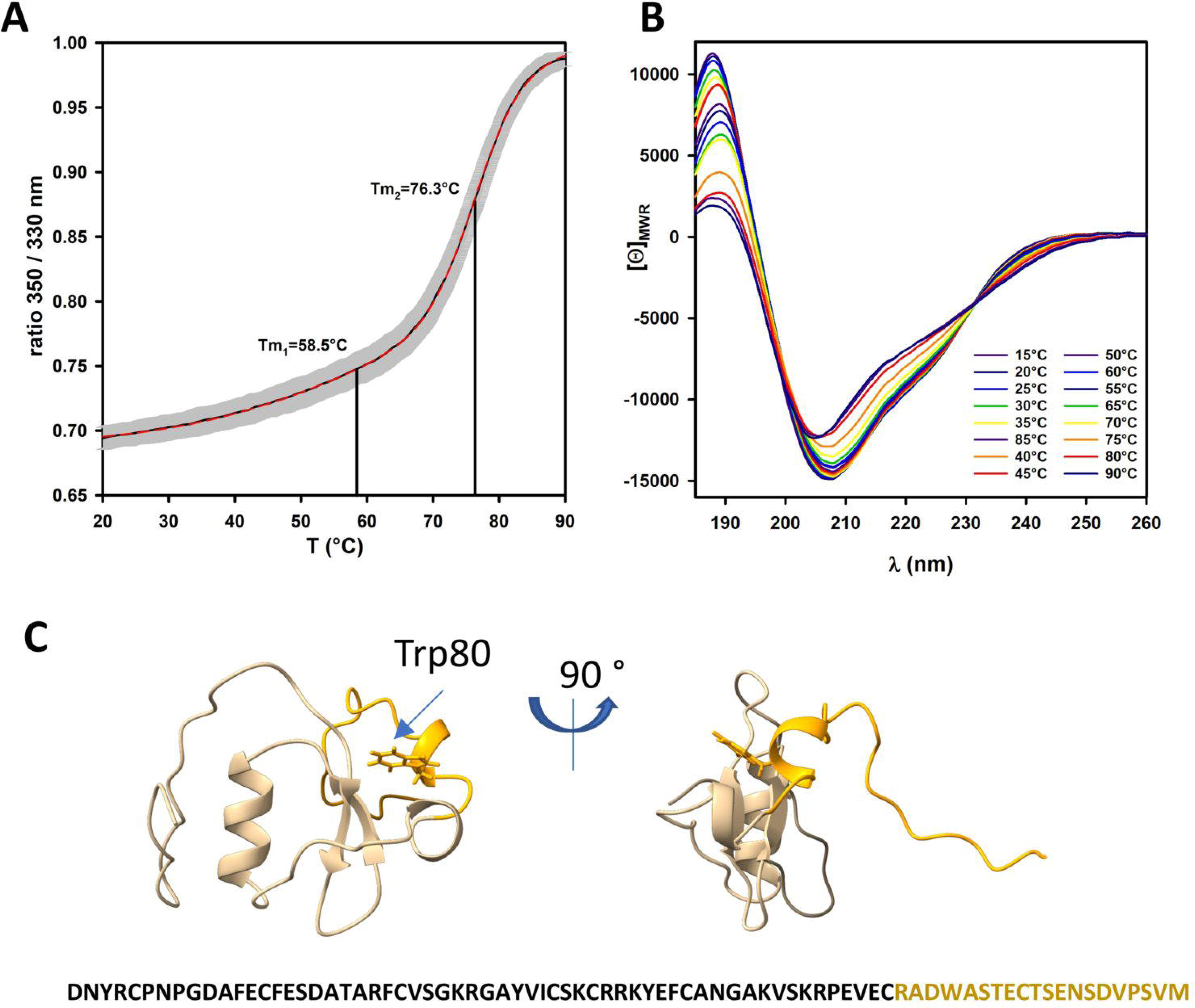
Biophysical and computational analysis of the thermal stability of BgSminSynth. A: Thermal denaturation profile of BgSmin^Synth^ monitored by nano differential scanning fluorimetry (nanoDSF). DSF spectra were acquired on a Prometheus NT.48 instrument (NanoTemper Technologies) using 5 µL of BgSmin^Synth^ at 30 µM. The intrinsic fluorescence ratio (350/330 nm) was plotted as a function of temperature. Thermal transitions were determined by fitting the data using a double-sigmoidal regression model. B: Thermal denaturation monitored by circular dichroism (CD). Full CD spectra were recorded between 185 and 260 nm over a temperature range of 15-90°C with increments of 5°C, revealing progressive changes in secondary structure upon heating. C: Molecular dynamics (MD) simulations performed from the X-ray crystal structure of BgSmin^Synth^. Three independent trajectories were generated for the BgSmin^A^ D88E analog using the GROMACS simulation package (version 2023.x) with the CHARMM36 force field and explicit TIP3P water (I = 70 mM NaCl). The panel shows a structural snapshot from trajectory 1 at 345 K (96.8 ns). The C-terminal region (residues 77-96) is highlighted in orange (cartoon representation), and the side chain of Trp80 is displayed.

Unlike the full-length sequence, which includes the signal sequence, the AlphaFold3 predicted models of the mature sequence of both native BgSmin sequences (residues 18-96) resemble the crystal structures of BgSmin^Synth^, with an average RMSD value of 0.6 Å comparable to those observed between the independent crystal molecules. This predictive accuracy supports the notion that schistosomin adopts a highly constrained and energetically well-defined fold, strongly encoded by its disulfide framework, and suggests that reliable structural models can be obtained for related schistosomin-like sequences. Retrospectively, these observations validate the folding strategy developed for the synthetic protein.

Given the compact architecture and dense disulfide connectivity revealed by crystallography, we next investigated the thermal stability of the schistosomin scaffold in solution using complementary biophysical approaches.

### 4. NanoDSF, CD and MD analysis supports the exceptional stability of the schistosomin fold

We first monitored temperature-induced conformational changes using nano differential scanning fluorimetry (nanoDSF), by recording the intrinsic tryptophan fluorescence ratio (350/330 nm) as a function of temperature (20-90°C) from the single tryptophan (Trp80) present in the BgSmin^Synth^. The fluorescence ratio displayed a biphasic temperature dependence, indicative of two apparent unfolding transitions (Figure 4A). Fitting the data using a double-sigmoidal model significantly improved the quality of the fit compared to a single-transition model, as evidenced by the analysis of residuals (See Suppl. Info.). The two apparent midpoint temperatures were determined to be Tm_1_= 58.5 ± 0.1°C and Tm_2_= 76.3 ± 0.02°C. Notably, the second transition was sharp and highly reproducible across replicates, in contrast to the first transition.

In parallel, we performed complementary circular dichroism (CD) experiments by monitoring full spectral CD scans recorded between 185 and 260 nm over a temperature range of 15-90°C. These scans confirmed a progressive loss of secondary structural features upon heating (Figure 4B). Analysis of the mean residue ellipticity ([Θ]_MRW_) at 222 nm and 208 nm revealed a similar biphasic behavior, with a major transition centered at 75.1 ± 0.5°C (See Suppl. Info.), whereas the lower-temperature transition (49.0 ± 4.2°C) appeared broader and less well-defined.

The same high-temperature transition values, measured by NanoDSF and CD measurements are consistent with the compact and stable architecture of schistosomin and reflect the full denaturing protein. In contrast, the lower-temperature transition, which differs between NanoDSF and CD experiments up to 13°C may reflect partial protein destabilization. To explain this observation, we performed molecular dynamics simulations from the BgSmin^Synth^ structure devoid of the Lys(Biot) residue (BgSmin^A^ D88E)at three temperatures below the second transition (25°C, 65°C, and 72°C; see details in Suppl. Info.). At 65°C (338 K), the simulations revealed elevated RMSF values in the C-terminal region (residues 77 to 96 including Trp80 and Cys85). At 72°C (345 K), one replicate exhibited transient unpacking of this region from the protein core (Figure 4C), suggesting that the first unfolding transition can correspond to the break of the Cys52-Cys85 disulfide bond.

### 5. Molecular dynamics simulations of A78P analog

To explore the potential impact of the amino acid residue present at position 78 (Ala versus Pro) on the conformation of BgSmin, we performed MD simulations of BgSmin^A^ D88E and BgSmin^P^ D88E at 300 K at an ionic strength corresponding to those of the hemolympth of the snail (I= 70 mM NaCl). MD trajectories between the A and P isoforms revealed no significant differences in overall conformational behavior (See details in Suppl. Info). Backbone RMSD, radius of gyration, and RMSF profiles were nearly indistinguishable between BgSmin^A^ and BgSmin^P^ across independent simulations. Local conformational analyses indicated that A78 can adopt a Φ and Ψ angles distribution that are well tolerated by proline residue, in line with (i) the structure-based proline design tool (Proscan web server: https://proscan.ibbr.umd.edu) (32), which identified position 78 as highly permissive to proline substitution when applied to the schistosomin crystal structure (PDB 9FDO, chain A) and (ii) the identical AF3 models between BgSmin^A^ and BgSmin^P^. Therefore, A78P variation represents a conservative substitution that preserves both the structural integrity and the dynamic properties of the schistosomin scaffold.

### 6. Structural convergence among schistosomin-like sequences and relationship to conotoxin disulfide-rich peptides

To verify our hypothesis that schistosomin fold is strongly encoded by its disulfide framework, and that reliable structural models can be obtained for related schistosomin-like sequences, we compiled, analyzed and run AlphaFold3 on all currently available sequences identified across molluscan species, which are so far restricted to freshwater and marine gastropods. A substantial fraction of these sequences originates from ConoServer (https://www.conoserver.org/), a curated database dedicated to cone snail peptides, reflecting the long-standing focus on venom-derived miniproteins (33). As a consequence, cone snail sequences are overrepresented in current datasets, a bias that likely reflects uneven sampling rather than a true biological restriction of this scaffold to venomous lineages (31).

On the basis of sequence features and cysteine topology, several of these cone snail peptides are currently classified in ConoServer as class XXII conotoxins. However, this assignment remains purely descriptive, as members of this class are largely uncharacterized and lack functional annotation. Importantly, the presence of closely related sequences in non-venomous gastropods strongly suggests that these peptides should not be viewed as canonical conotoxins, but rather as representatives of a broader and previously unrecognized family of disulfide-rich molluskan miniproteins.

Sequence analyses reveal that the 17 confident schistosomin-like protein sequences identified so far—corresponding to full-length mature peptides after removal of predicted signal peptides—share a strictly conserved cysteine framework despite overall sequence identities ranging from 18.6 to 98.7% (Figure 5A). Among these, only the schistosomins from *B. glabrata* and *L. stagnalis* have been experimentally characterized, while the remaining sequences are inferred from transcriptomic data. This conserved cysteine pattern therefore constitutes a robust molecular signature for identifying novel Smin-like peptides in omics datasets. AlphaFold3 models were generated and structural superposition of the 17 predicted models using FoldSeek (34) showed a common compact three-dimensional architecture similar to our BgSmin^Synth^ structure (Figures 5B and 5C) and the presence of distinct but closely related clusters (Figure 5C). In particular, peptides from *Conus* species (*Caenogastropoda*) tend to group separately from schistosomins of *Heterobranchia* gastropods such as *Biomphalaria, Bulinus* and *Lymnaea*, while remaining within the same overall structural family.

**Figure 5.**
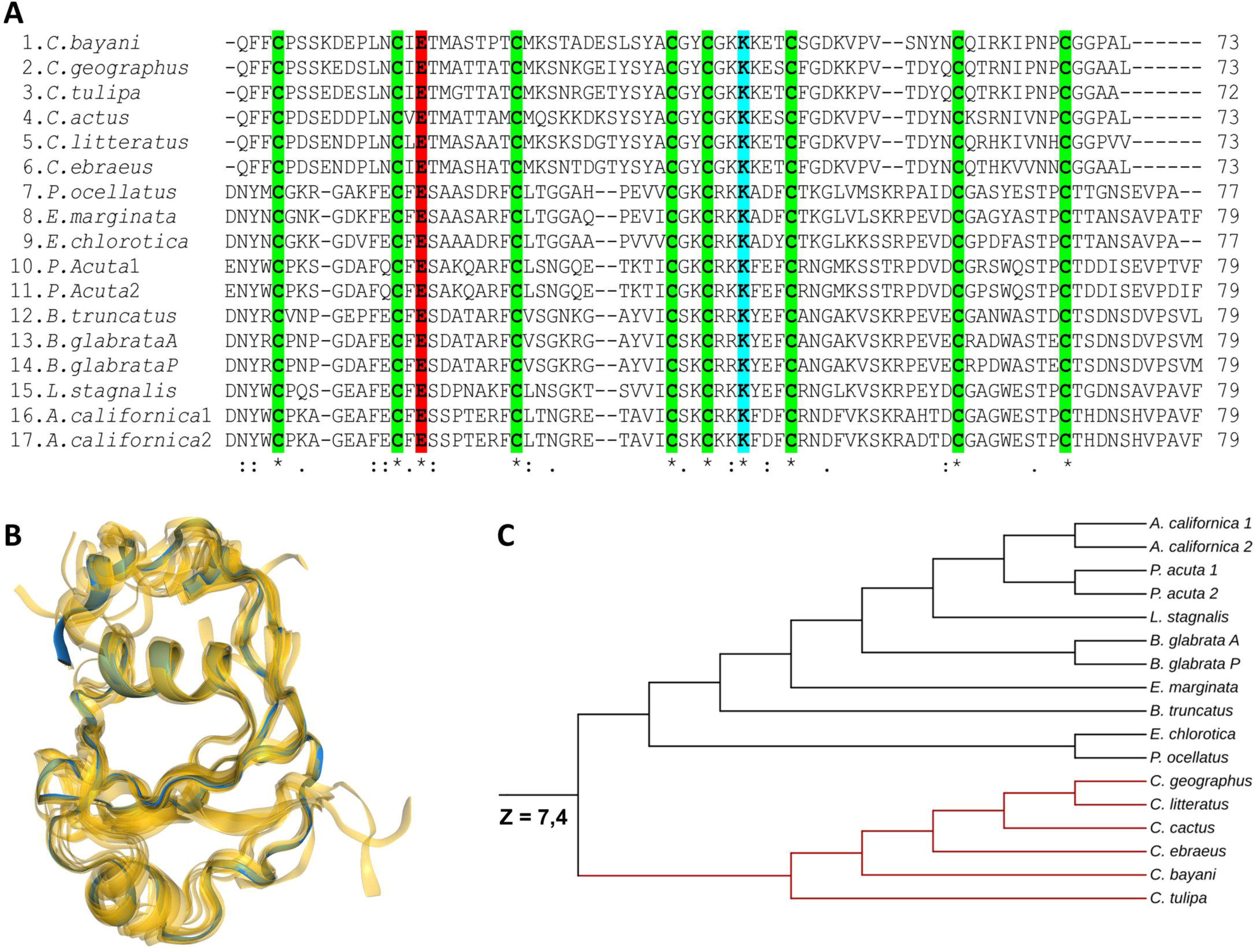
Sequences and structural comparison of schistosomin-like peptides across molluscs A. Sequence alignment of schistosomin-like peptides. Multiple sequence alignment of 17 schistosomin-like proteins identified across gastropods species, including freshwater snails (*Lymnaea stagnalis*, *Biomphalaria glabrata*, *Bulinus truncatus*), sea slugs (*Aplysia californica*, *Elysia marginata*, *Elysia chlorotica*, *Plakobranchus ocellatus*, *Physella acuta*), and marine cone snails (*Conus* spp.). Sequences correspond to predicted mature peptides after removal of signal peptides. Conserved cysteine residues are highlighted and define a characteristic disulfide framework shared across all sequences. Conserved residues are indicated by an asterisk (*), strongly similar residues by a colon (:), and weakly similar residues by a period (.). B. Structural comparison of schistosomin-like peptides. Superposition of AlphaFold3-predicted structures of the 17 schistosomin-like peptides using FoldSeek. Despite low sequence identity, all structures display a highly similar compact fold stabilized by four conserved disulfide bonds, illustrating strong structural conservation across gastropods lineages, including peptides currently classified as class XXII conotoxins. C. Structural similarity dendrogram from FoldSeek. Hierarchical clustering of schistosomin-like peptides based on pairwise structural similarity (Z-scores) computed using the DALI algorithm. The heatmap and dendrogram reveal both a high overall structural conservation (Z-scores ≥ 7.4) and the presence of distinct structural clusters, separating primarily *Conus*-derived peptides from heterobranch gastropod schistosomins. Higher Z-scores indicate stronger structural similarity.

These results highlight a remarkable combination of structural conservation and diversification: schistosomin-like proteins have diverged extensively at the sequence level while preserving a highly conserved disulfide-stabilized fold. The presence of distinct structural clusters within this conserved framework suggests that evolutionary diversification has occurred under strong structural constraints, likely imposed by the stability and functional requirements of the scaffold.

These observations establish schistosomin as the prototype of a broadly distributed family of disulfide-rich miniproteins whose architecture is encoded by the cysteine disulfides. They further provide a strong rationale for expanding the repertoire of Smin-like peptides and integrating structural, phylogenetic, and functional approaches to investigate their evolutionary trajectories and biological roles across molluscs.

### 7. Schistosomin expression in the *B. glabrata* organs

Although schistosomin was originally described as a neuroendocrine factor potentially involved in host-parasite interactions and parasitic castration in *L. stagnalis* (27, 29), its precise biological function in *B. glabrata* remains unresolved (30). In this context, we examined the tissue distribution of schistosomin at both the transcript and protein levels, taking advantage of the isoform-specific qPCR strategy and the antibodies generated using the synthetic protein.

Quantitative isoform-specific RT-qPCR analyses allowed discrimination between the two schistosomin isoforms, BgSmin^A^ and BgSmin^P^, and revealed a marked tissue-specific expression pattern. The highest transcript levels were detected in the foot, nervous ganglia, mantle, and tentacles, whereas hepatopancreas, and hemocytes exhibited low or undetectable expression levels (Figure 6A). Importantly, in all tissues analyzed, the BgSmin^A^ isoform was consistently and significantly more highly expressed than isoform P. However, no tissue displayed exclusive expression of either isoform, indicating that both variants are co-expressed across tissues, albeit at different relative levels. Overall, this transcriptional profile indicates that schistosomin expression is not restricted to neuronal tissues but extends to several peripheral and epithelial compartments.

**Figure 6:**
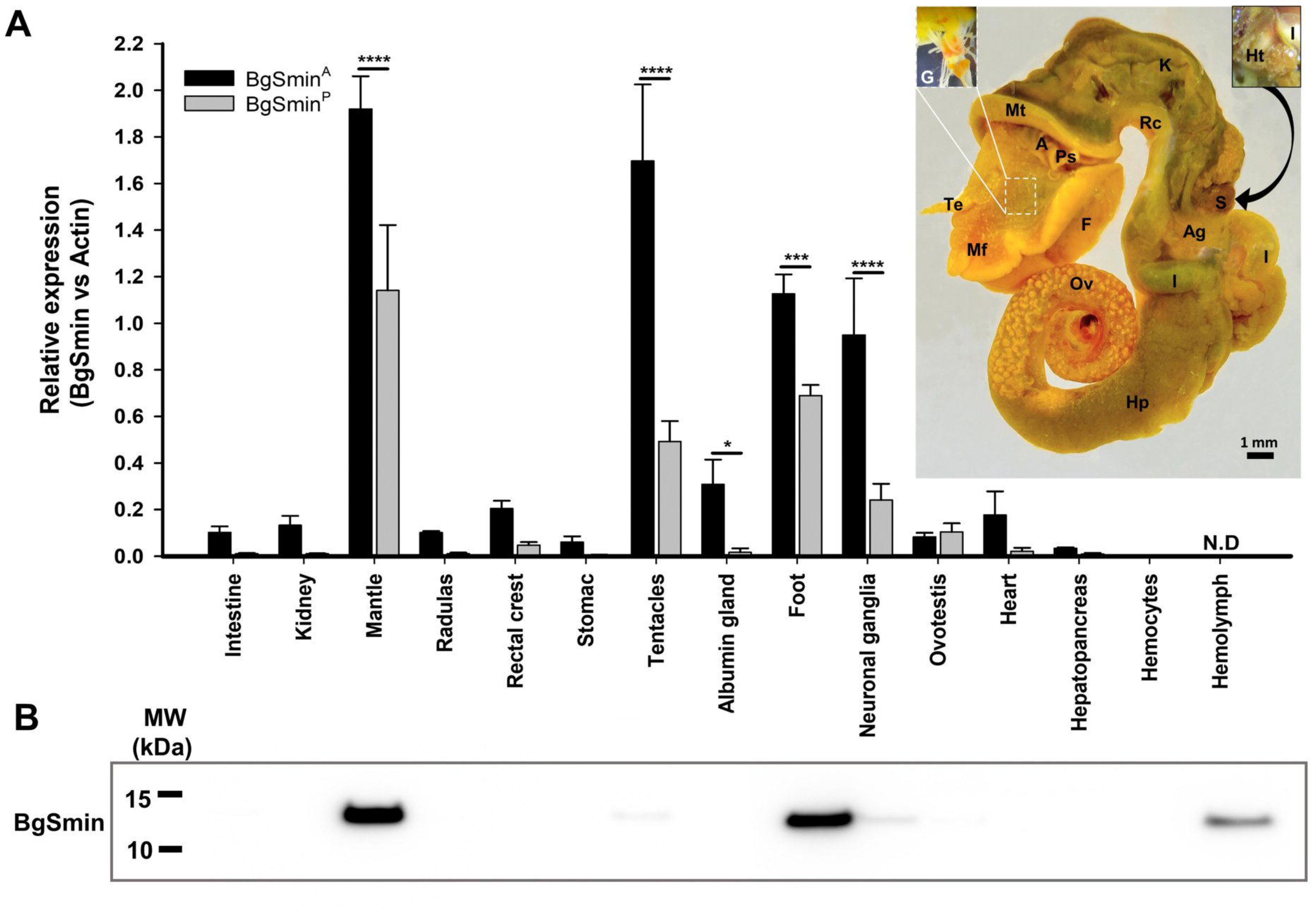
Tissue distribution of schistosomin transcripts and protein in *B. glabrata*. B. Isoform-specific RT-qPCR analysis of schistosomin expression across *B. glabrata* tissues (n = 3 biological replicates). Relative transcript levels of BgSmin^A^ (black bars) and BgSmin^P^ (grey bars) were quantified and normalized to actin expression. The anatomical origin of each sampled tissue is indicated on the dissected *B. glabrata* specimen. Abbreviations: Albumen gland (Ag), Anus (A), Foot (F), Ganglia (G), Heart (H), Hepatopancreas (Hp), Intestine (I), Kidney (K), Mantle (Mt), Mantle fold (Mf), Pseudobranch (Ps), Ovotestis (Ov), Rectal crest (Rc), Stomach (S), and Tentacle (Te). Hemocytes were collected from the hemolymph of approximately one hundred snails. No RNA was extracted from hemolymph and RT-qPCR was therefore not performed (N.D.). C. Western blot analysis of schistosomin in selected tissues. Protein extracts were obtained from the same biological samples used for the RT-qPCR analysis. A single immunoreactive band with an apparent molecular mass between ∼10 and 15 kDa is detected, consistent with the expected size of the mature schistosomin protein.

At the protein level, Western blot analyses detected schistosomin in the foot and nervous ganglia, but also in the mantle and tentacles (Figure 6B). While no signal was observed in hepatopancreas or hemocytes, schistosomin was readily detected in the hemolymph. This apparent discrepancy between low transcript levels and detectable protein abundance in the hemolymph is consistent with the secreted nature of schistosomin. Indeed, the protein contains a signal peptide and has previously been reported as a circulating factor in snail hemolymph (26, 29). These observations suggest that schistosomin is synthesized in specific tissues and subsequently released into the circulatory system, where it can distribute independently of local transcriptional activity, thereby partially decoupling mRNA abundance from protein levels measured by immunoblotting. Notably, the protein-level data reflect the overall tissue distribution of schistosomin rather than isoform-specific expression patterns because Western blot analysis cannot distinguish the two schistosomin isoforms.

These results demonstrate that schistosomin is expressed in multiple tissues and circulates within *B. glabrata*. Remarkably, strong expression in the foot, mantle, and tentacles, tissues that are directly exposed to the external environment, raises the possibility that schistosomin may participate in interactions with surrounding biological agents such as microorganisms or parasites. The absence of a strictly neuron-specific expression pattern, combined with its secreted and systemic distribution, argues against a classical neuropeptide function (35–37) and indicates that the physiological role of schistosomin in this gastropod species remains to be elucidated.

In this respect, *B. glabrata* provides access to schistosomin-like peptides expressed in non-venomous tissues and physiological contexts, in contrast to cone snails, where related disulfide-rich peptides are predominantly studied in the framework of venom specialization. Moreover, the availability of well-established laboratory breeding conditions enables reproducible access to biological material, making this organism a tractable experimental model to investigate the structural, evolutionary, and functional properties of this family independently of venom-associated constraints. The tools developed in this study therefore provide a robust foundation to address the long-standing question of schistosomin-like protein function in gastropods and in the host-parasite relationship.

## Aknowlegments

We gratefully acknowledge CNRS and IDRIS for providing access to the Jean Zay supercomputer (AD010816084R1) for conducting the molecular dynamics simulations. We acknowledge SOLEIL for provision of synchrotron radiation facilities (proposals ID 20191181) in using PROXIMA beamlines. We deeply thank Pierre Legrand on PROXIMA 1 beamline for running the first AF2 model.

