## Supplementary information figures and tables for "Total synthesis and structural characterization of a novel protein scaffold from the snail *Biomphalaria glabrata*"

#### 1. General information for protein chemical synthesis

##### Reagents and solvents

1-[*bis*(dimethylamino)methylene]-1*H*-1,2,3-triazolo[4,5-*b*]pyridinium 3-oxid hexafluorophosphate (HATU), 2-(1*H*-Benzotriazol-1-yl)-1,1,3,3-tetramethyluronium fluorophosphates (HBTU) and *N*-Fmoc protected amino acids were obtained from Iris Biotech GmbH.

3-mercaptopropionic acid (97%, MPA), 4-mercaptophenylacetic acid (97%, MPAA), *tris*(2-carboxyethyl)phosphine hydrochloride (98%, TCEP·HCl), guanidine hydrochloride (Gn·HCl, 99%), triisopropylsilane (TIS), thiophenol, thioanisole, ethane-1,2-dithiol (EDT), hydrochloric acid (reagent grade, 37% w/v), glacial acetic acid (AcOH), acetic anhydride ($Ac_2O$), trifluoroacetic acid (TFA), sodium hydroxide (pellets, 97%), piperidine, *N,N,N*-diisopropylethylamine (DIEA), triethylamine (TEA, as well as other reagents were purchased from suppliers of chemicals (TCI, Acros Organics, Sigma-Aldrich, Merck…) and were of the purest grade available.

Peptide synthesis grade *N*,*N*-dimethylformamide (DMF), dichloromethane (DCM), 1,2-dichloroethane (DCE), acetone, methanol (MeOH), ethanol (EtOH), diethyl ether ($Et_2O$), acetonitrile, heptane, LC–MS-grade acetonitrile (0.1% TFA), LC–MS grade water (0.1% TFA) and trifluoroacetic acid (TFA) were purchased from Biosolve and Fisher-Chemical. Water was purified with a Milli-Q Ultra Pure Water Purification System.

Bis(2-sulfenylethyl)amino trityl 1% divinylbenzene crosslinked polystyrene solid support (SEA solid support) was produced on demand by Roowin.

Solvents and reagents were used as received.

**Caution:**
TFA is an extremely corrosive liquid; great care must be taken when using this reagent. Proper eye protection, lab coat, and gloves are mandatory. TFA must be manipulated under a fume hood.

DMF is a highly toxic solvent. Proper eye protection, lab coat, and gloves are mandatory. DMF must be manipulated under a fume hood.

##### Instrumentation

The following instrumentation was used to synthesize, purify and characterize the peptides prepared in this work.

Unless otherwise stated, peptides were synthesized using an automated peptide synthesizer (Intavis) and standard Fmoc solid phase peptide synthesis methods.

Reaction mixtures or isolated compounds were characterized by analytical UPLC–MS using a System Ultimate 3000 UPLC (Thermofisher) equipped with a C18 column, a diode array detector and a mass spectrometer (Ion trap LCQfleet, heat temperature 350 °C, spray voltage 2.8 kV, capillary temperature

350 °C, capillary voltage 10 V, tube lens voltage 75 V). Unless otherwise stated, analyses were performed using an appropriate linear gradient of eluent B (0.1% TFA in $CH_3CN$) in eluent A (0.1% TFA in $H_2O$) at a flow rate of 0.4 mL/min.

Peptides were purified by preparative RP-HPLC using a Buchi system equipped with C18 column and an UV detector. Purifications were performed using an appropriate linear gradient of eluent B (0.1% TFA in $CH_3CN$) in eluent A (0.1% TFA in $H_2O$) at a flow rate of 6 mL/min. Selected fractions were combined, frozen and lyophilized.

#### 2. Chemical synthesis of schistosomin

##### 2.1. Sequence of synthetic schistosomin

| | | | |
|---|---|---|---|
| tr\|B5L013\|B5L013_BIOGL | 1 | MKTVFFILALIVCAVVADNYRCPNPGDAFECFESDATARFCVSGKRGAYVICSKCRRKYEFCANGAKVSKRPEVECRADW | 80 |
| tr\|A0A9W3AQB7\|...A9W3AQB7_BIOGL | 1 | MKTVFIILALIVCAVVADNYRCPNPGDAFECFESDATARFCVSGKRGAYVICSKCRRKYEFCANGAKVSKRPEVECRPDW | 80 |
| tr\|B5L013\|B5L013_BIOGL | 81 | ASTECTSDNSDVPSVM | 96 |
| tr\|A0A9W3AQB7\|...A9W3AQB7_BIOGL | 81 | ASTECTSDNSDVPSVM | 96 |

**Supplementary Figure 1.** Alignment of schistosomin isoforms A78 (B5L013) and P78 (A0A9W3AQB7).

Schistosomin A78 isoform is named $BgSmin^A$. Schistosomin P78 isoform is named $BgSmin^P$.

The sequence of synthetic schistosomin has been adapted from Uniprot entry B5L013 (isoform A78).

*Sequence of synthetic schistosomin*

DNYRCPNPGDAFECFESDATAR**F-C**VSGKRGAYVICSKCRRKYE
**F-C**ANGAKVSKRPEVECRADWASTECTS**E**NSDVPSVMK(Biot)-$NH_2$

Synthetic and biotinylated schistosomin is named $BgSmin^A{}_{Synth}$, the subscript A is to recall that an alanine residue is at position 78.

The junctions utilized for assembling the full-length protein and defining the peptide segments described hereinafter are highlighted in green. The protein is modified at the C-terminus by a lysine residue with a biotin group attached to its side-chain, to enable functional studies in the future. Note that the aspartic acid residue at position 88 (Uniprot numbering) has been substituted by a glutamic residue, since preliminary studies have revealed that D88-N89 junction is highly sensitive to aspartimide formation. The protein features a carboxamide group on its C-terminus.

#### 2.2. General synthetic strategy

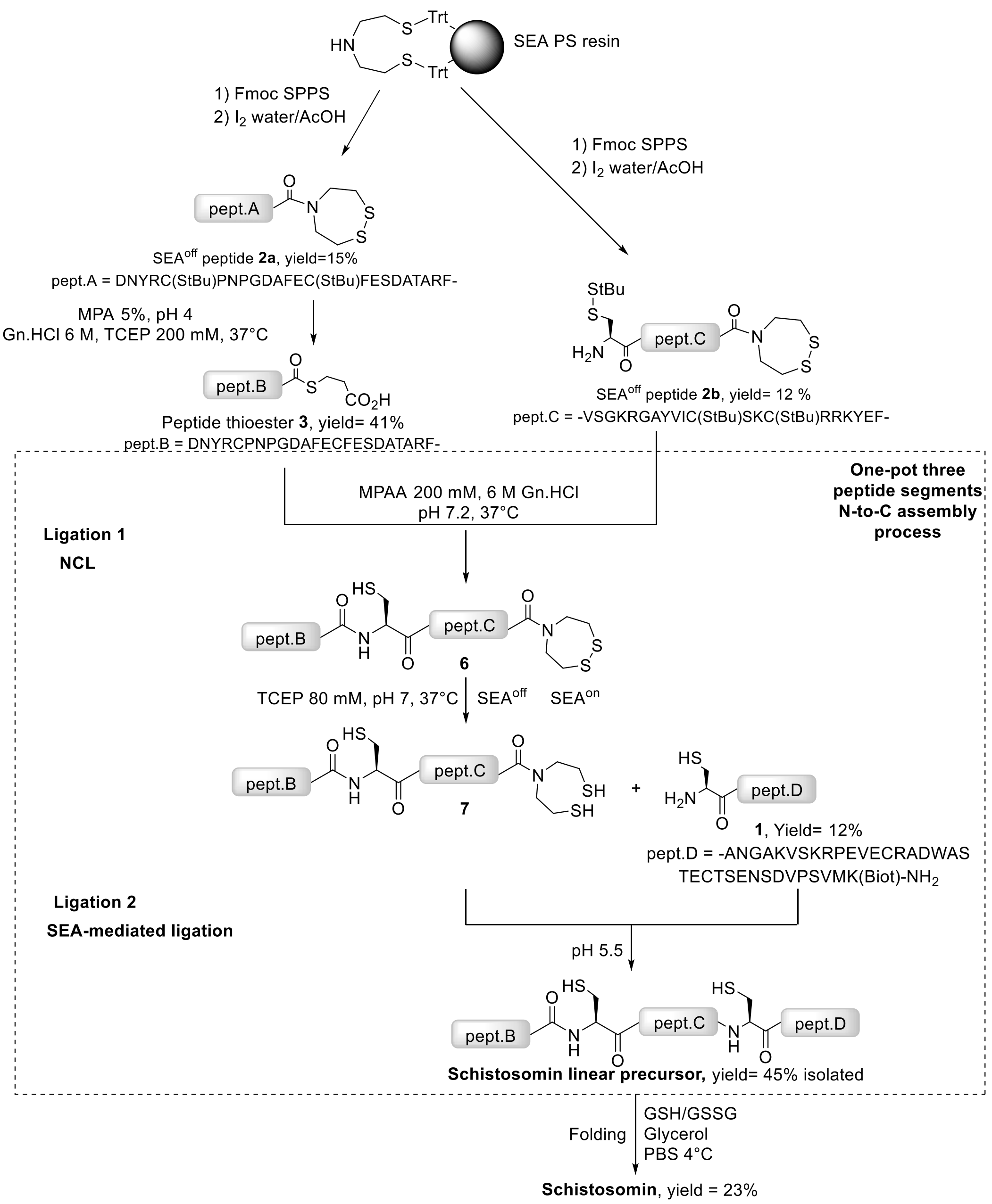

Supplementary Figure 2. General strategy for accessing schistosomin by chemical synthesis.

#### 2.3. Synthesis of the peptide segments

##### General protocols for automated peptide synthesis

Peptide elongation was performed using standard Fmoc/*tert*-butyl chemistry on an automated peptide synthesizer. Couplings were performed using 5-fold molar excess of each Fmoc-L-amino acid, 4.5-fold molar excess of TBTU, and 10-fold molar excess of DIEA. A capping step was performed after each coupling with $Ac_2O$/DIEA in DMF. At the end of the synthesis, the peptidyl solid support was washed with $CH_2Cl_2$, diethylether (3 × 2 min) and dried *in vacuo*.

Final peptide deprotection and cleavage: The elongated peptide was cleaved from the solid support using TFA with appropriate scavengers as indicated in each case. The crude peptide was precipitated by adding the TFA solution coming from the cleavage and deprotection step in ice-cold diethyl ether/heptane 1/1 v/v (20 mL per mL of TFA cocktail) and recovered by centrifugation. The solid was washed with cold diethyl ether/heptane 1/1 v/v (40 mL), dissolved in water and the solution was lyophilized.

##### Synthesis of peptide amide 1

Peptides were synthesized using standard Fmoc solid phase peptide synthesis methods on a NovaSyn TGR solid support (0.25 mmol/g).

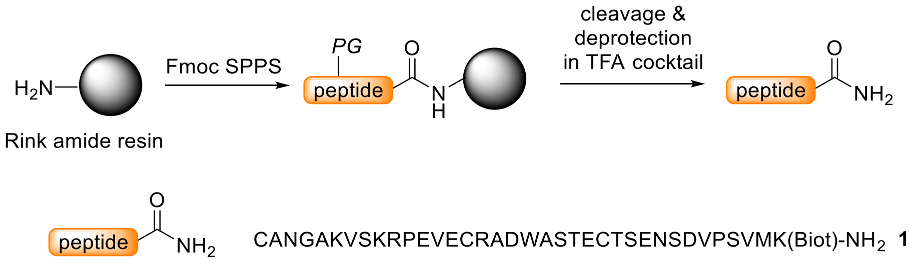

**Supplementary Figure 3.** Preparation of peptide amide **1**.

CANGAKVSKRPEVECRADWASTECTSENSDVPSVMK(Biot)-$NH_2$ **1**, 0.10 mmole scale

Cleavage cocktail: TFA/water/EDT/TIS 90/2.5/2.5/5 by vol

Yield: 56.8 mg (12.4%)

A)

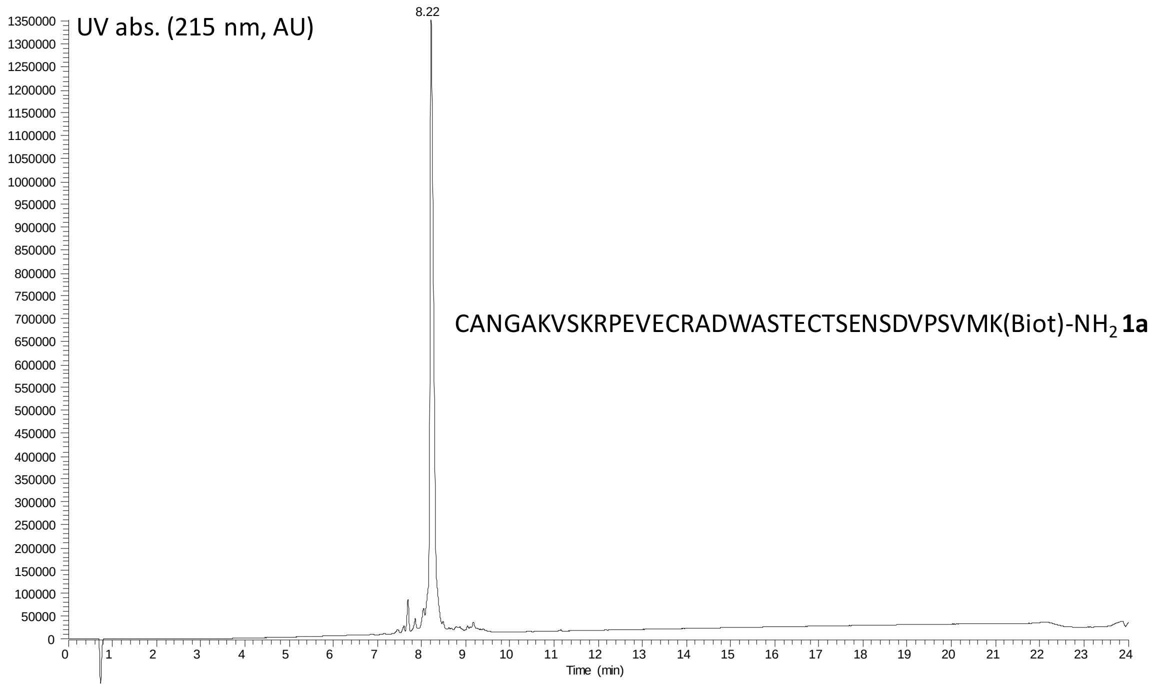

B)

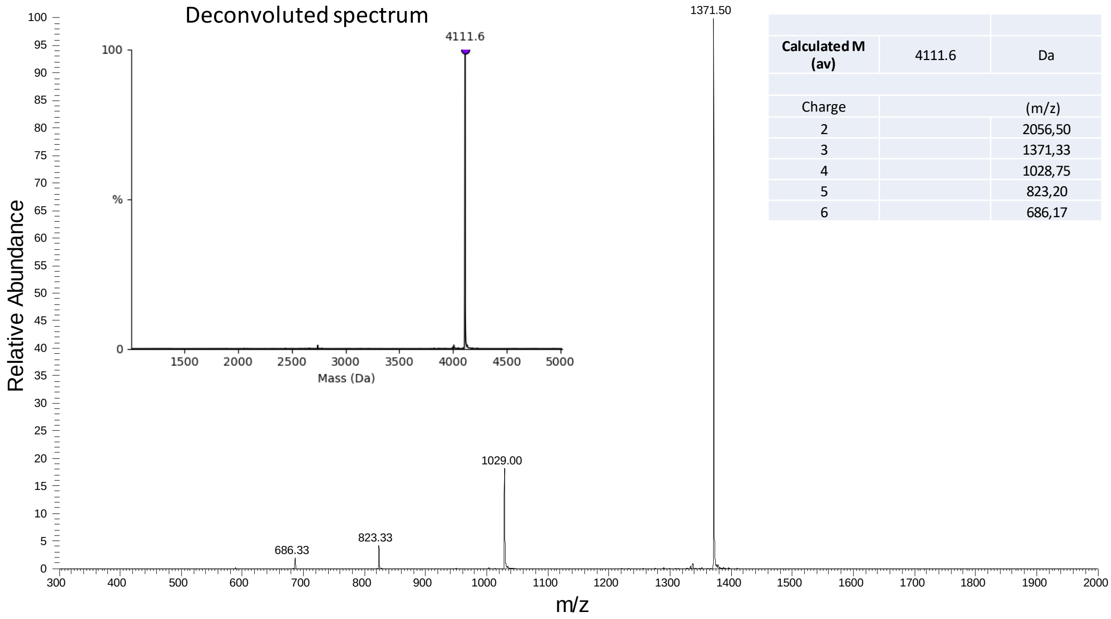

**Supplementary Figure 4.** UPLC-MS analysis of peptide **1a**. A) LC trace, eluent A 0.10% TFA in water, eluent B 0.10% TFA in $CH_3CN$. Acquity BEH C18 column, 300 Å, 1.7 µm (2.1 × 100 mm) column, gradient 0-70% B in 20 min, 70 °C, 0.4 mL/min, detection at 215 nm. B) MS trace. Deconvoluted spectrum $[M+H]^+$ m/z calcd. (av) 4111.6, found 4111.6.

##### Synthesis of SEA$^{off}$ peptide segments 2

Synthesis of *bis*(2-sulfanylethyl)aminotrityl polystyrene (SEA PS, 0.16 mmol/g) resin was carried out as described elsewhere.[1]

Peptide elongation was performed on SEA PS resin using standard Fmoc/*tert*-butyl chemistry on an automated peptide synthesizer. Typical procedures for the synthesis of SEA$^{off}$ peptide segments were described in detail in previous papers. For detailed protocols see refs [1b, 2]. The synthesis requires that all Cys residues are temporarily protected by a tert-butylsulfenyl (StBu) group, that remains on the peptide following the deprotection and cleavage step in TFA.

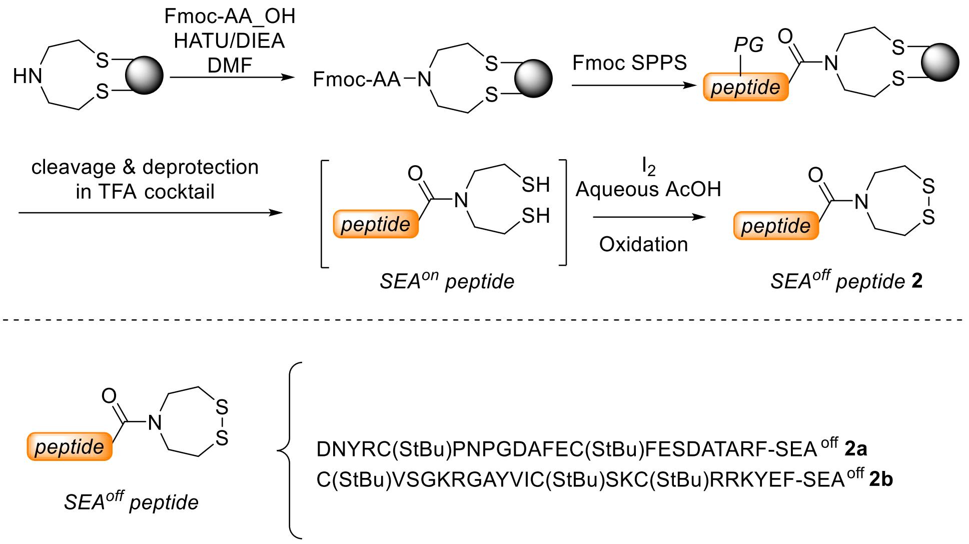

**Supplementary Figure 5.** Synthesis of SEA$^{off}$ peptides **2**.

DNYRC(StBu)PNPGDAFEC(StBu)FESDATARF-SEA$^{off}$ **2a** was prepared as described elsewhere (0.1 mmole scale).[1a]

Cleavage cocktail: TFA/water/thioanisole/thiophenol//TIS 87.5/2.5/2.5/2.5/5 by vol

Yield: 47.3 mg (14.5%)

A)

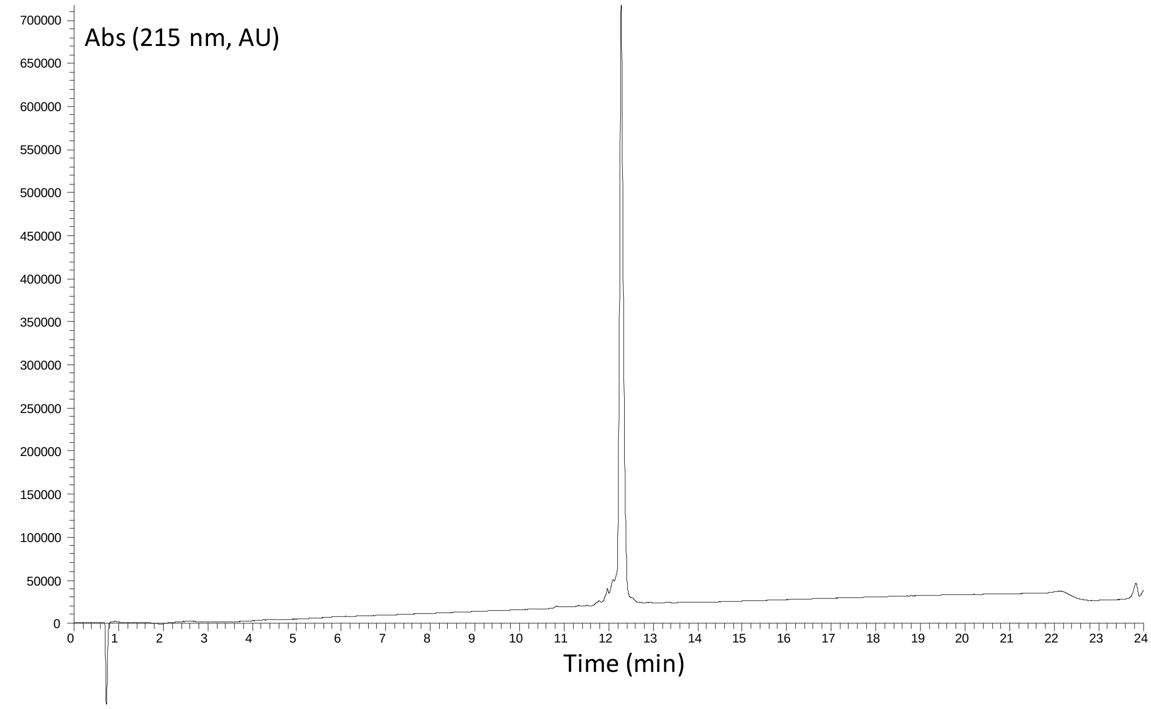

B)

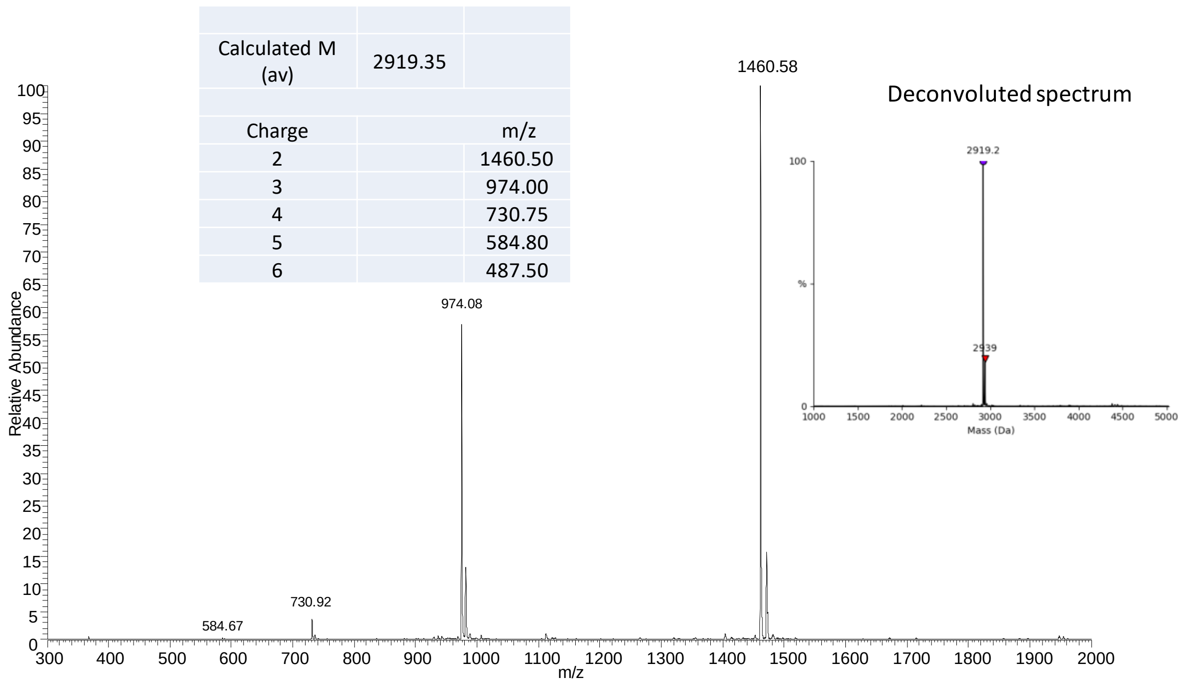

| | | |
|---|---|---|
| Calculated M (av) | 2919.35 | |
| Charge | | m/z |
| 2 | | 1460.50 |
| 3 | | 974.00 |
| 4 | | 730.75 |
| 5 | | 584.80 |
| 6 | | 487.50 |

**Supplementary Figure 6.** UPLC-MS analysis of peptide **2a**. A) LC trace, eluent A 0.10% TFA in water, eluent B 0.10% TFA in $CH_3CN$. Acquity BEH C18 column, 300 Å, 1.7 µm (2.1 × 100 mm) column, gradient 0-70% B in 20 min, 70 °C, 0.4 mL/min, detection at 215 nm. B) MS trace.

C(StBu)VSGKRGAYVIC(StBu)SKC(StBu)RRKYEF-SEA$^{off}$ **2b** was prepared as described elsewhere (0.1 mmole scale).[1a]

Cleavage cocktail: TFA/water/thioanisole/thiophenol//TIS 87.5/2.5/2.5/2.5/5 by vol

Yield: 42.5 mg (12%)

A)

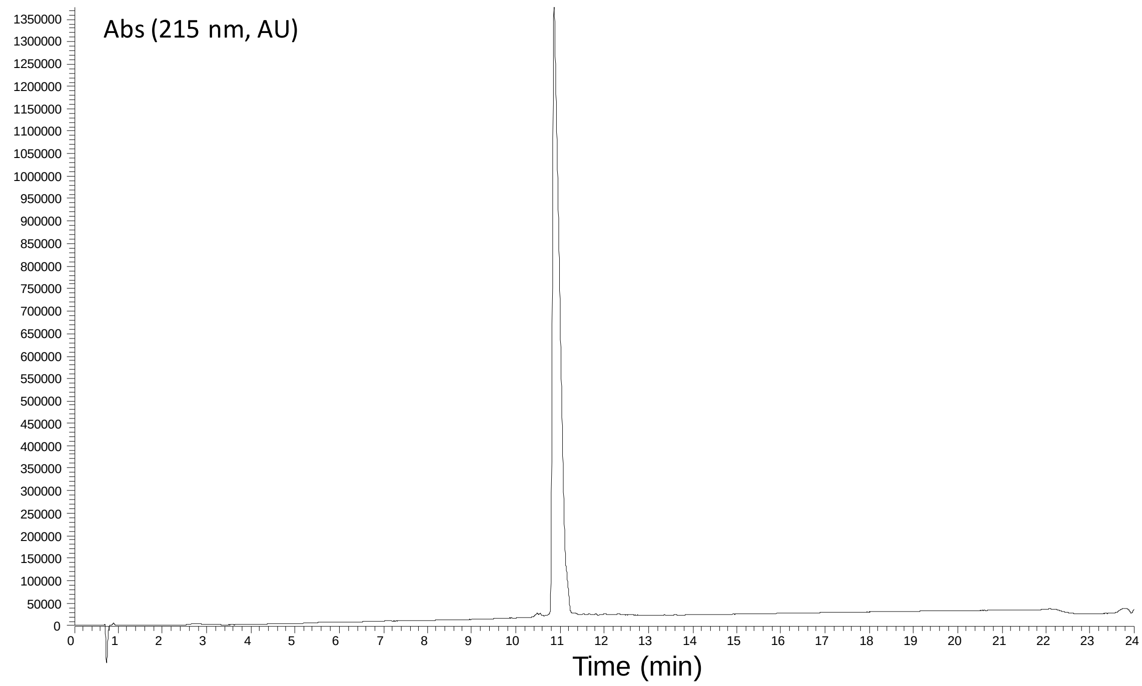

B)

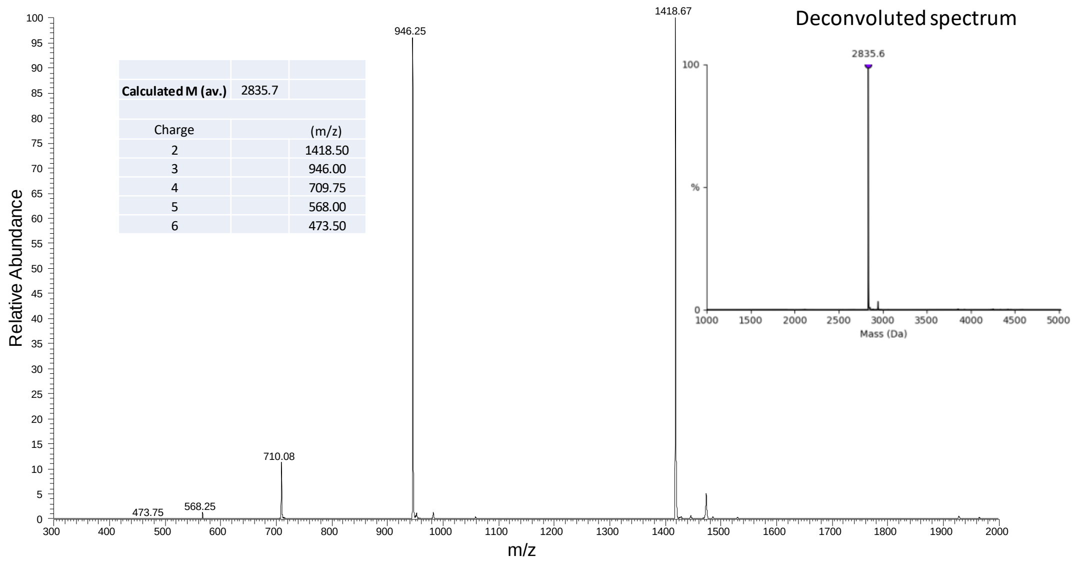

**Supplementary Figure 7.** UPLC-MS analysis of peptide **2b**. A) LC trace, eluent A 0.10% TFA in water, eluent B 0.10% TFA in $CH_3CN$. Acquity BEH C18 column, 300 Å, 1.7 µm (2.1 × 100 mm) column, gradient 0-70% B in 20 min, 70 °C, 0.4 mL/min, detection at 215 nm. B) MS trace.

**Synthesis of MPA peptide thioester 3**

Peptide thioester **3** was prepared by exchanging the SEA group of SEA peptide **2a** by 3-mercaptopropionic acid (MPA) at pH 4.0 according to published procedures (Supplementary Figure 8).[1b, 2-3]

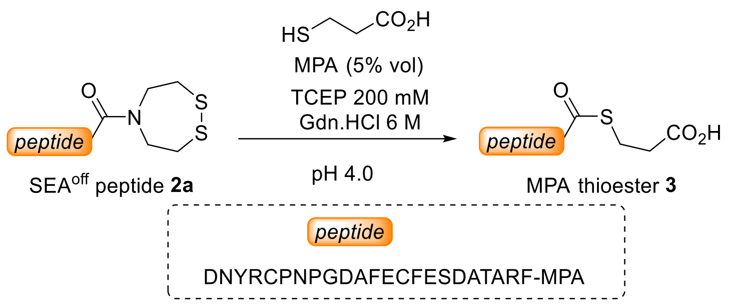

**Supplementary Figure 8.** Synthesis of peptide thioesters from SEA peptides.

The exchange has been performed on 22.3 mg (6.60 µmoles) of SEA$^{off}$ peptide **2a**. The reaction provided 8.5 mg (41 %) of peptide thioester **3** after HPLC purification.

A)

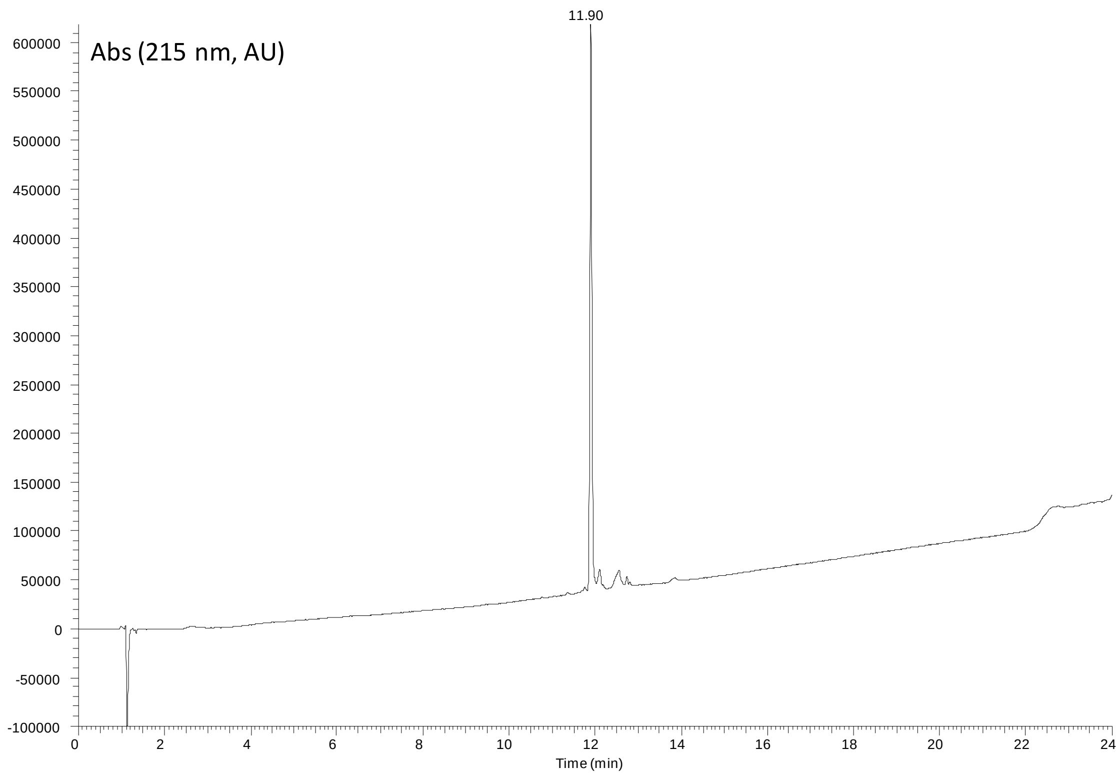

B)

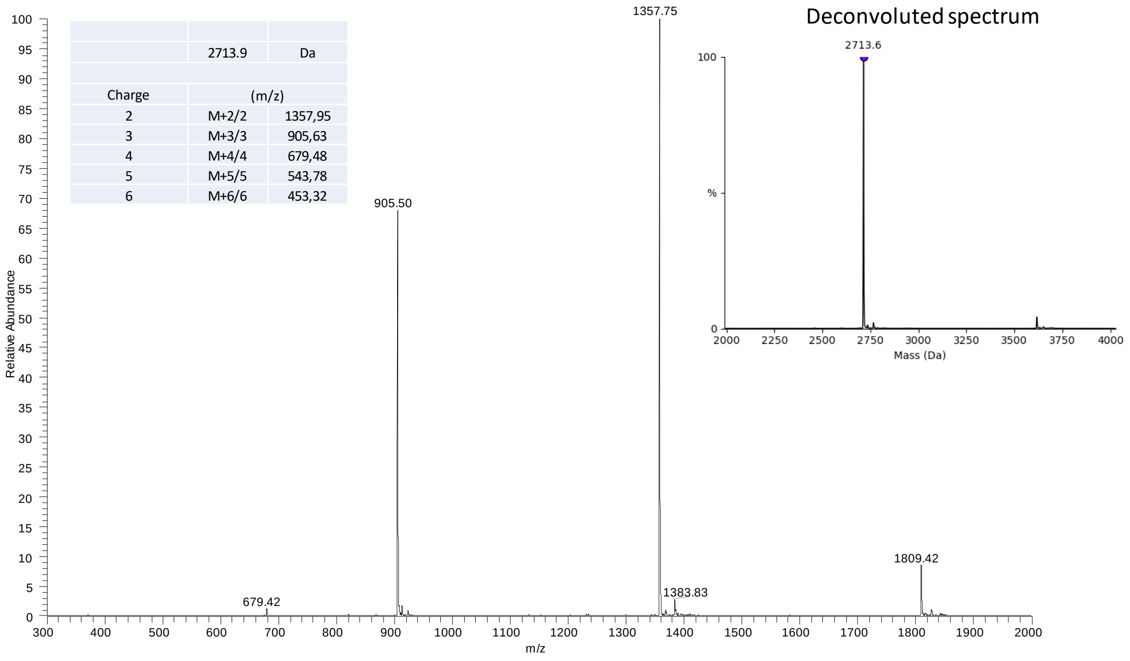

**Supplementary Figure 9.** UPLC-MS analysis of peptide **3**. A) LC trace, eluent A 0.10% TFA in water, eluent B 0.10% TFA in $CH_3CN$. Acquity BEH C18 column, 300 Å, 1.7 µm (2.1 × 100 mm) column, gradient 0-70% B in 20 min, 70 °C, 0.4 mL/min, detection at 215 nm. B) MS trace.

#### 2.4. Assembly of schistosomin

*SEA-mediated ligation (step 1)*

Peptide thioester **3** (13.0 mg , 4.2 µmoles) and SEA$^{off}$ peptide **2b** (15.5 mg soit 4.2 µmoles) were dissolved in 0.1 M pH 7.2 sodium phosphate buffer containing 6 M Gn.HCl, 200 mM 4-mercaptophenylacetic acid (MPAA) and 10 mM *N*-octyl glucoside. The reaction mixture was agitated at 37 °C overnight.

An aliquot (5 µL) was diluted with 20% aqueous acetic acid (95 µL) and extracted with diethylether three times to remove the excess of MPAA before UPLC-MS analysis. Then, TCEP.HCl (5 µL, 200 mM in water, final concentration 10mM) was added and the aliquot was kept at rt for 1 h.

UPLC-MS analysis (Supplementary Figure 10) showed the successful formation intermediate **6**.

A)

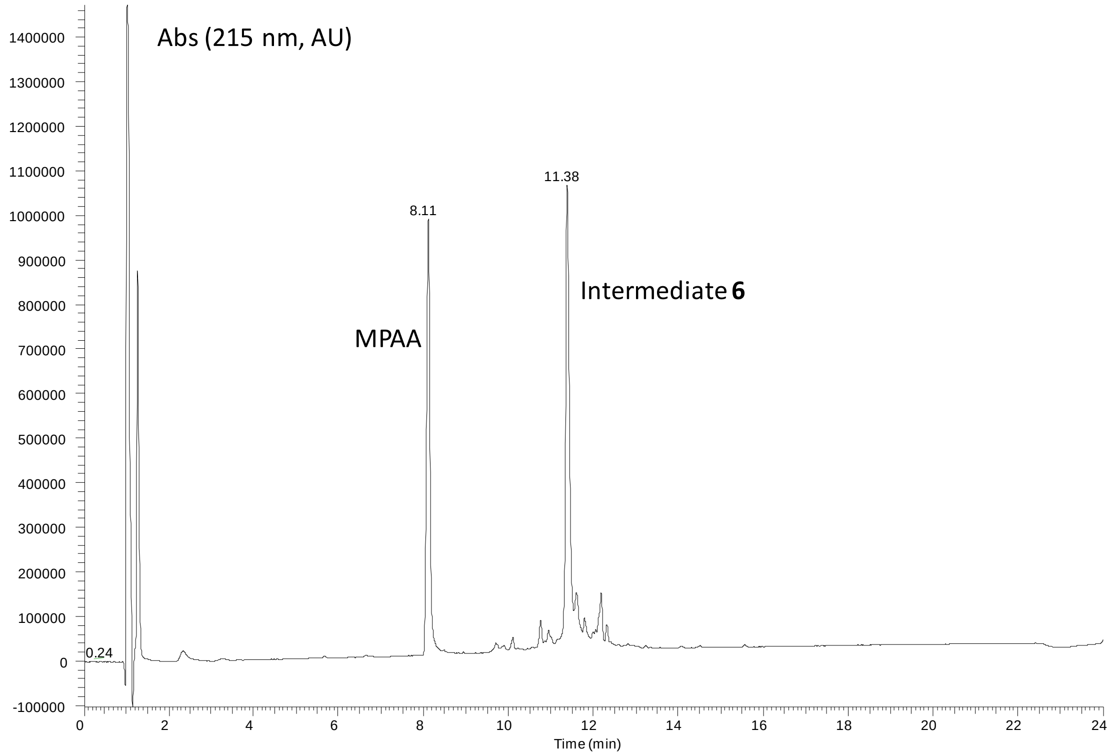

B)

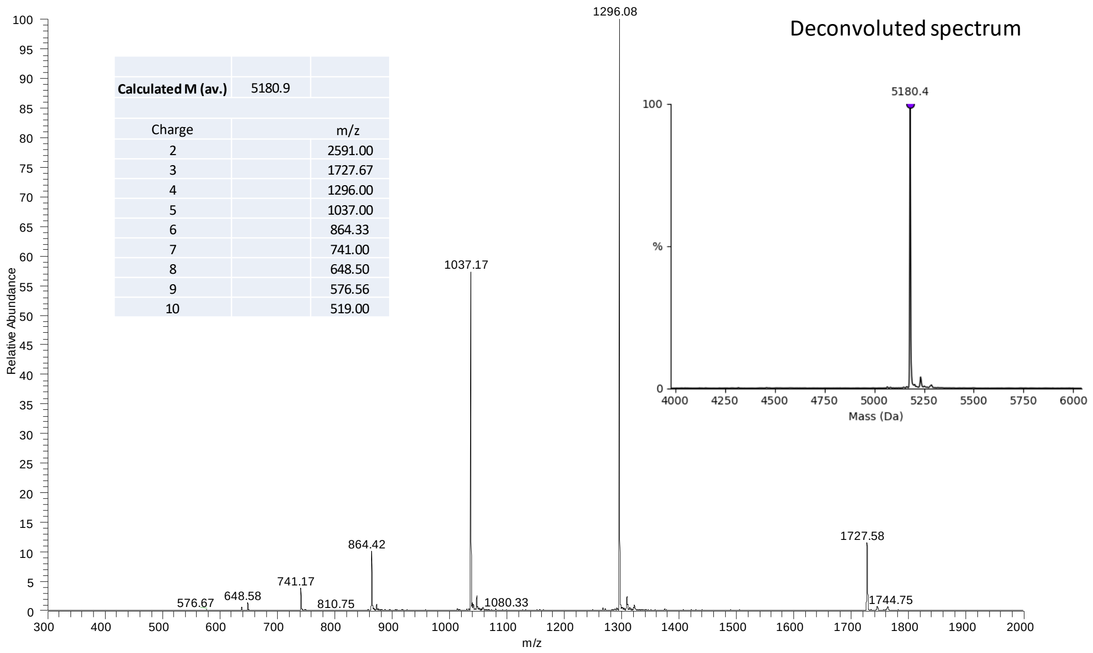

**Supplementary Figure 10.** UPLC-MS analysis of crude intermediate **6** formed during the first step of schistosomin assembly (5 µL of an extracted aliquot treated with TCEP). A) LC trace, eluent A 0.10% TFA in water, eluent B 0.10% TFA in $CH_3CN$. Acquity BEH C18 column, 300 Å, 1.7 µm (2.1 × 100 mm) column, gradient 0-70% B in 20 min, 70 °C, 0.4 mL/min, detection at 215 nm. B) MS trace.

*NCL (step 2)*

To the above mixture was added peptide segment **1** (21.8 mg, 4.65 µmoles) solubilized in 0.1 M pH 5.5 sodium phosphate buffer containing 6 M Gn.HCl, 200 mM MPAA and 10 mM *N*-octyl glucoside (840 µL). The pH of the reaction mixture reaction mixture was adjusted to 5.5 using 6 M HCl and agitated at 37 °C for 44 h.

Aliquots (10 µL) were diluted with 20% aqueous acetic acid (95 µL) and extracted with diethylether three times to remove the excess of MPAA before UPLC-MS analysis.

A)

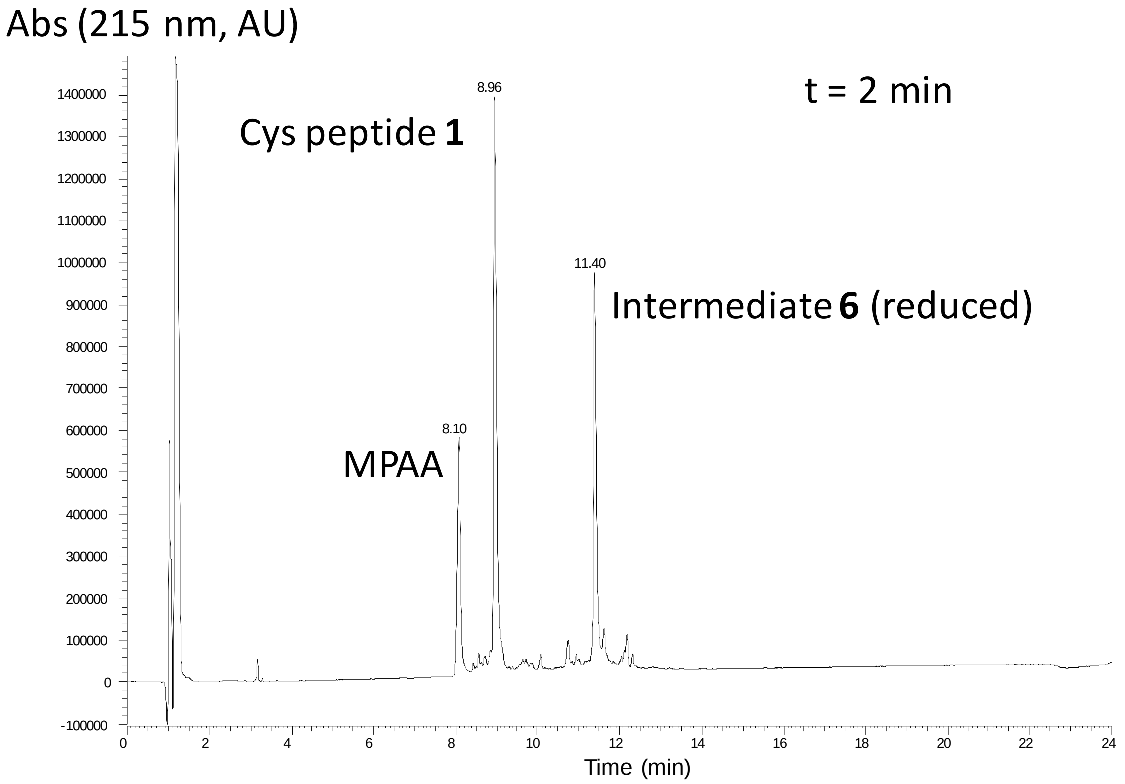

B)

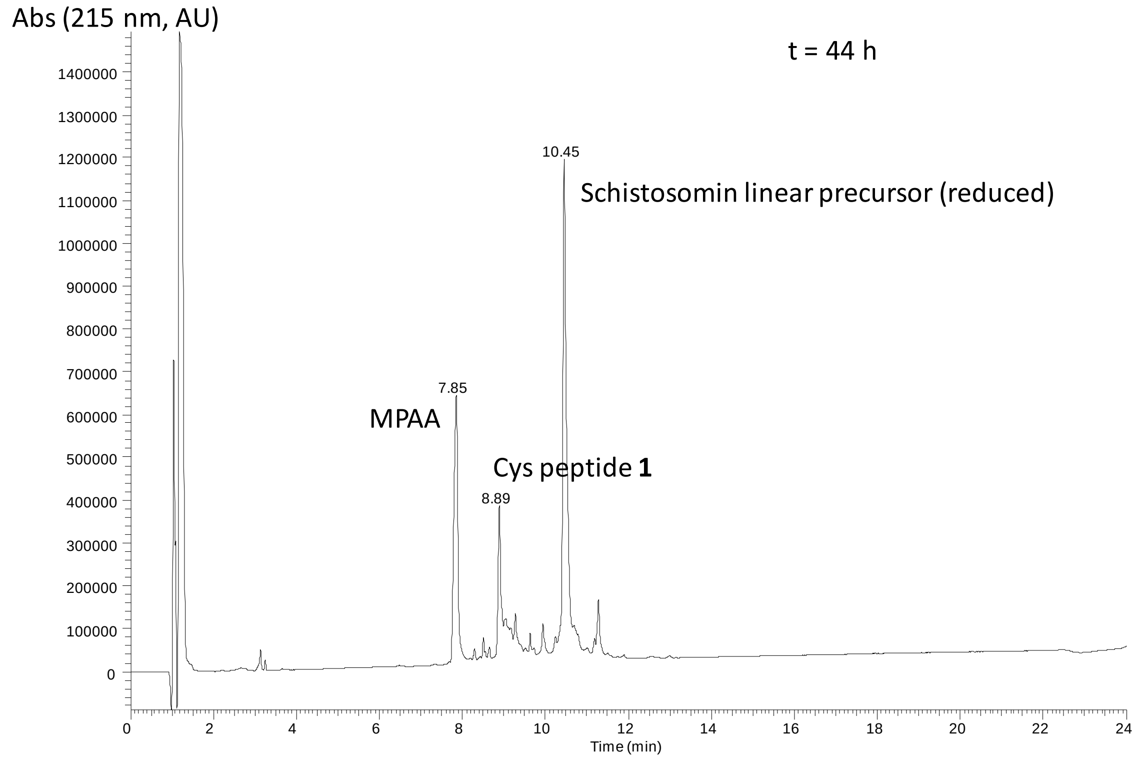

C)

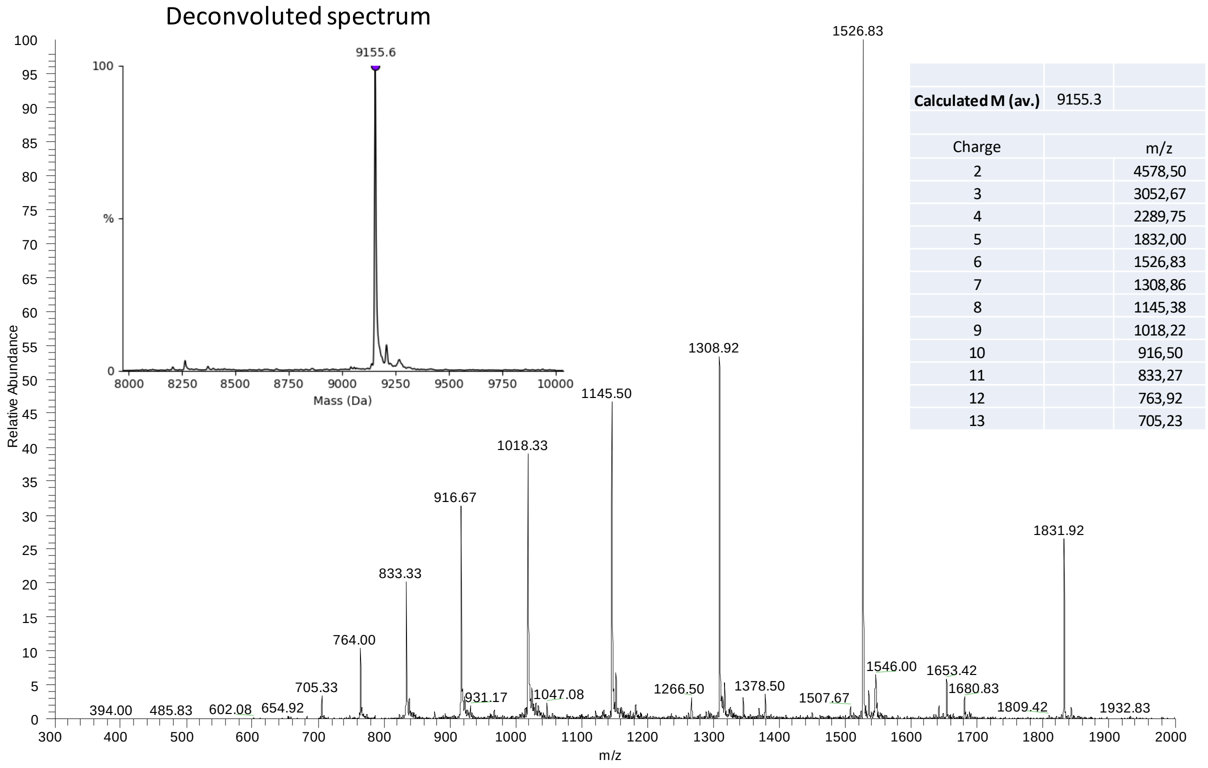

| | |
|---|---|
| **Calculated M (av.)** | 9155.3 |
| Charge | m/z |
| 2 | 4578,50 |
| 3 | 3052,67 |
| 4 | 2289,75 |
| 5 | 1832,00 |
| 6 | 1526,83 |
| 7 | 1308,86 |
| 8 | 1145,38 |
| 9 | 1018,22 |
| 10 | 916,50 |
| 11 | 833,27 |
| 12 | 763,92 |
| 13 | 705,23 |

**Supplementary Figure 11.** UPLC-MS analysis of the second step of schistosomin assembly leading to the formation of schistosomin linear precursor (10 µL of an extracted aliquot). A) LC trace after 2 min., eluent A 0.10% TFA in water, eluent B 0.10% TFA in $CH_3CN$. Acquity BEH C18 column, 300 Å, 1.7

µm (2.1 × 100 mm) column, gradient 0-70% B in 20 min, 70 °C, 0.4 mL/min, detection at 215 nm. B) MS trace of the peak at 10.45 min (t = 44 h).

*HPLC purification*

The reaction mixture was diluted with aqueous acetic acid (10 mL, 20% in deionized water) and then extracted with diethylether to remove the excess of MPAA (2 mL × 16). The excess of diethylether was removed by bubbling argon into the solution for 15 min. The crude peptide was purified using a Xbridge BEH C18 300 Å 5 µm 10 mm × 250 mm column, flow rate 6 mL/min, 65 °C, eluent A 0.10% TFA in water, eluent B 0.10% TFA in CH3CN, gradient 0-10% B in 5 min then 10-40% B in 60 min.

The purified fractions were collected, frozen and lyophilized to provide 19.98 mg (45.2 %) of schistosomin linear precursor.

A)

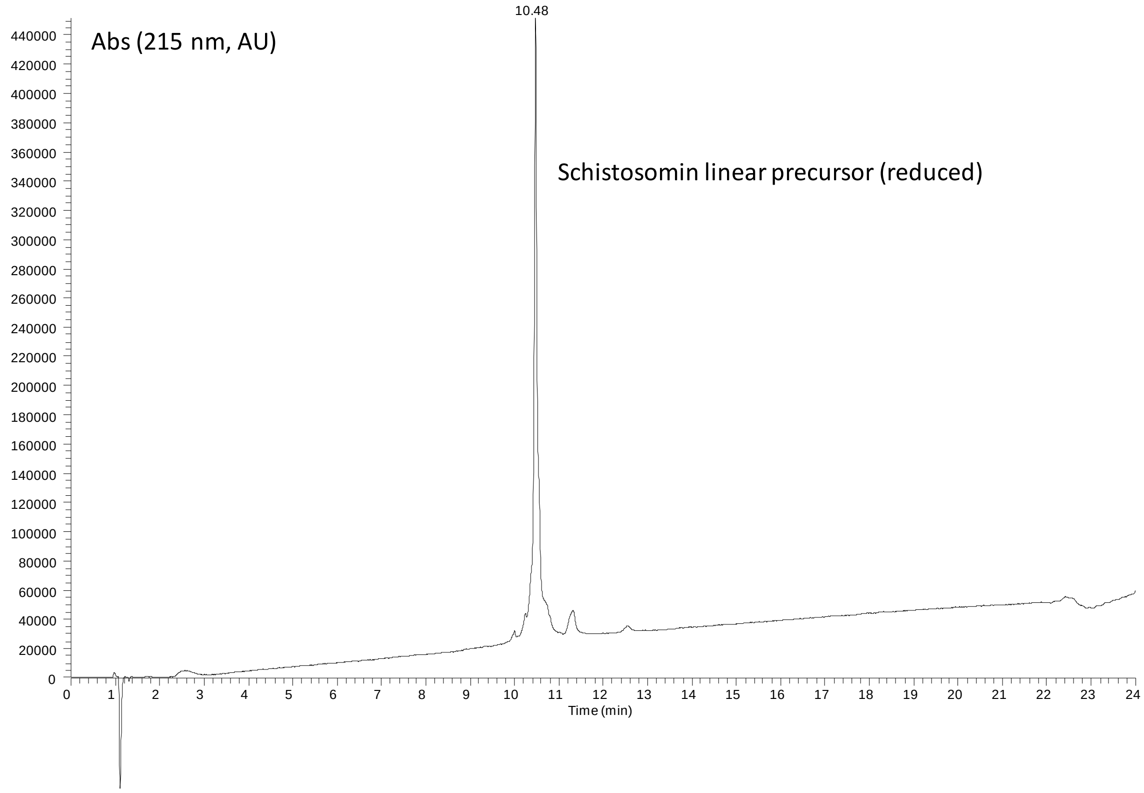

B)

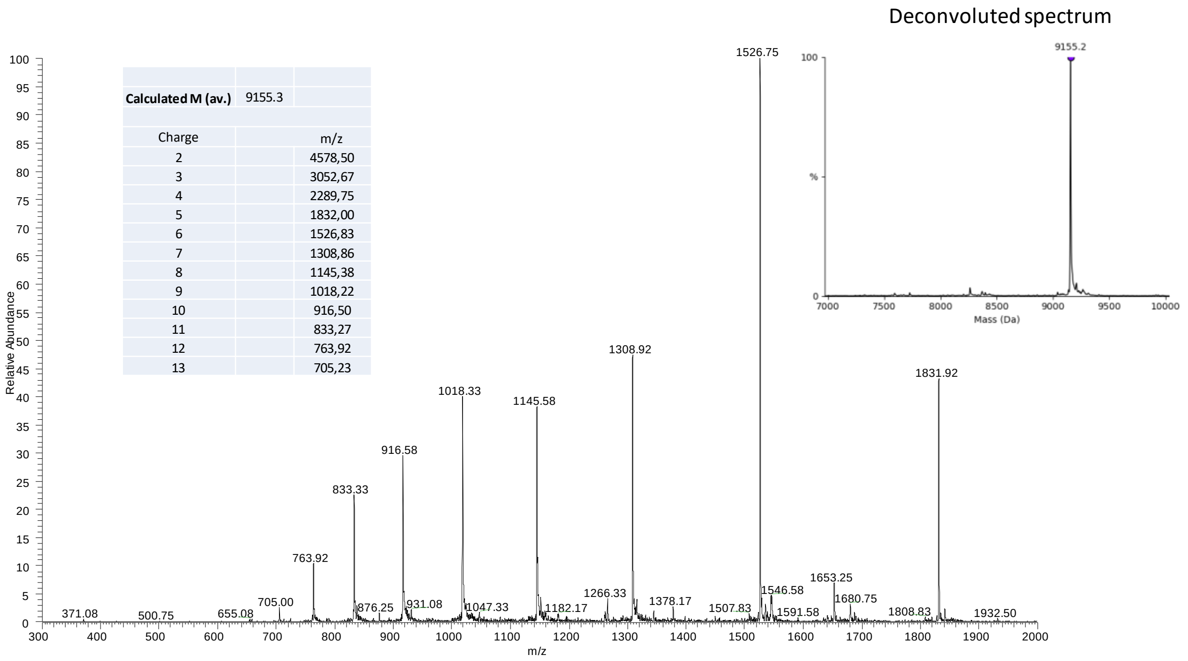

**Supplementary Figure 12.** UPLC-MS analysis of purified schistosomin linear precursor. A) LC trace, eluent A 0.10% TFA in water, eluent B 0.10% TFA in $CH_3CN$. Acquity BEH C18 column, 300 Å, 1.7 µm (2.1 × 100 mm) column, gradient 0-70% B in 20 min, 70 °C, 0.4 mL/min, detection at 215 nm. B) MS trace.

##### 2.5. Folding of schistosomin

The folding of schistosomin linear precursor was performed at 4 °C. Extensive experimentations enabled us to develop a robust procedure minimizing protein precipitation during folding, which requires 29 days.

Schistosomin linear precursor (11 mg) was dissolved in PBS buffer containing 6 M Gn.HCl (1.1 mL). This solution was diluted with a PBS buffer containing glycerol as a co-solvent (10% by vol), N-octyl glucoside (2 mM), and redox system GSH (1 mM)/GSSG (0.2 mM). The folding mixture was gently agitated for 29 days.

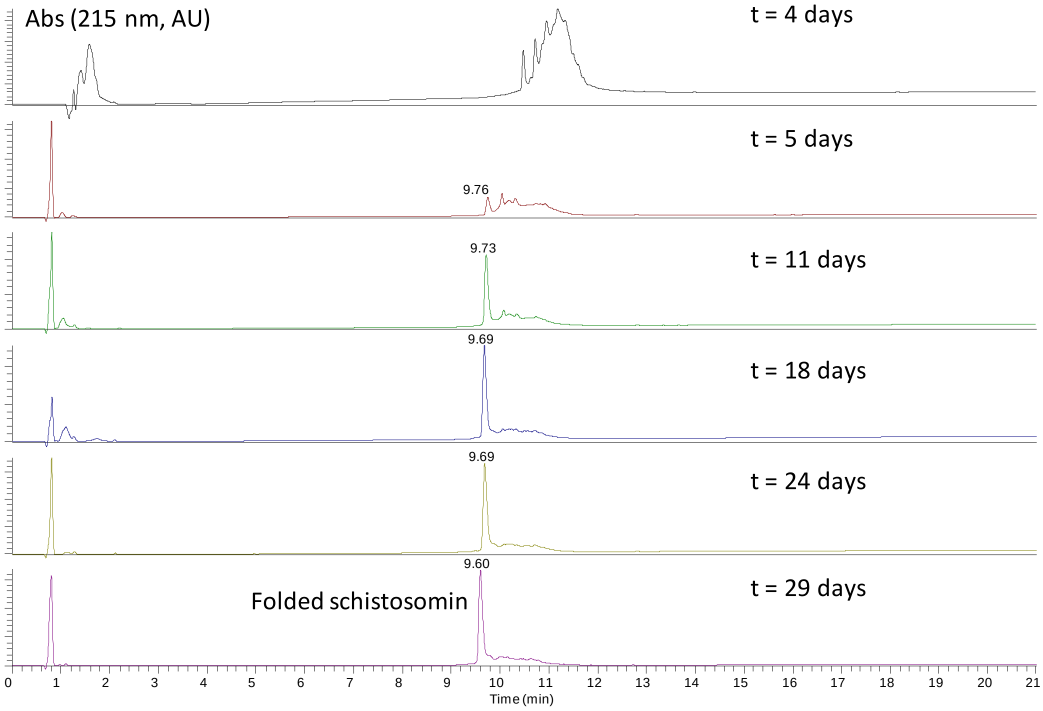

**Supplementary Figure 13.** UPLC-MS analysis of the folding schistosomin mixture. A) LC traces, eluent A 0.10% TFA in water, eluent B 0.10% TFA in $CH_3CN$. Acquity BEH C18 column, 300 Å, 1.7 µm (2.1 × 100 mm) column, gradient 0-70% B in 20 min, 70 °C, 0.4 mL/min, detection at 215 nm. B) MS trace.

The folded schistosomin product was purified by HPLC as described above for the linear precursor. The collected fractions were pooled, frozen and lyophilized to provide 2.3 mg (23 %) of folded schistosomin.

##### 2.6. Characterization of synthetic schistosomin

A)

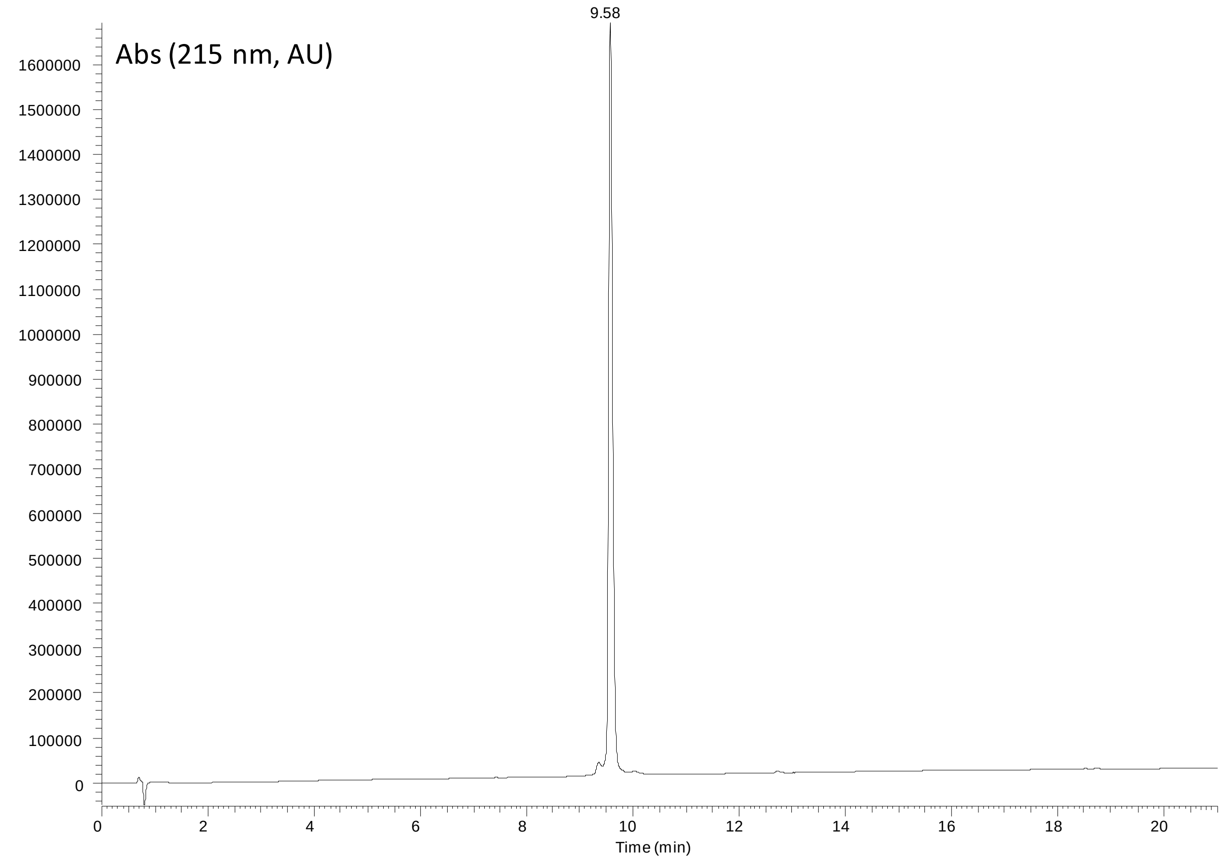

B)

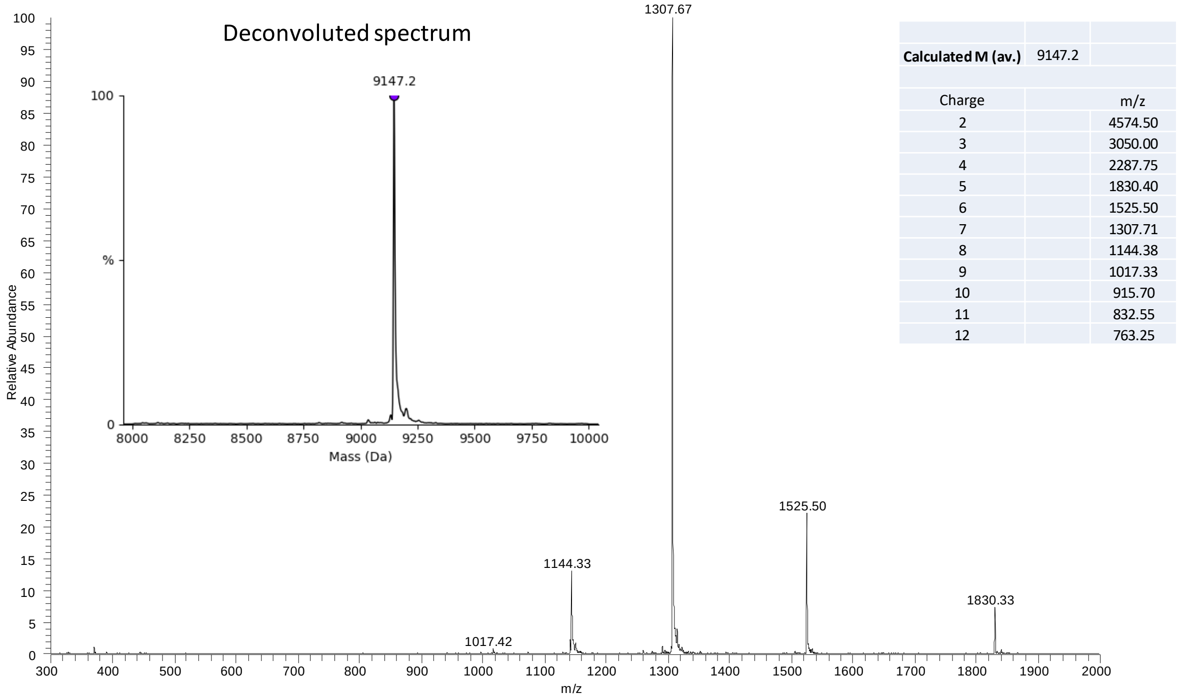

| | | |
|---|---|---|
| **Calculated M (av.)** | 9147.2 | |
| Charge | | m/z |
| 2 | | 4574.50 |
| 3 | | 3050.00 |
| 4 | | 2287.75 |
| 5 | | 1830.40 |
| 6 | | 1525.50 |
| 7 | | 1307.71 |
| 8 | | 1144.38 |
| 9 | | 1017.33 |
| 10 | | 915.70 |
| 11 | | 832.55 |
| 12 | | 763.25 |

**Supplementary Figure 14.** UPLC-MS analysis of folded schistosomin after HPLC purification. A) LC trace, eluent A 0.10% TFA in water, eluent B 0.10% TFA in $CH_3CN$. Acquity BEH C18 column, 300 Å, 1.7 µm (2.1 × 100 mm) column, gradient 0-70% B in 20 min, 70 °C, 0.4 mL/min, detection at 215 nm. B) MS trace.

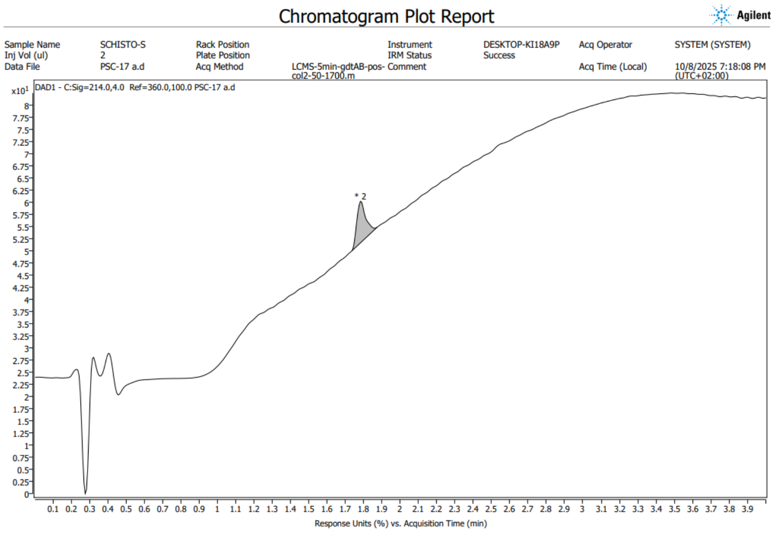

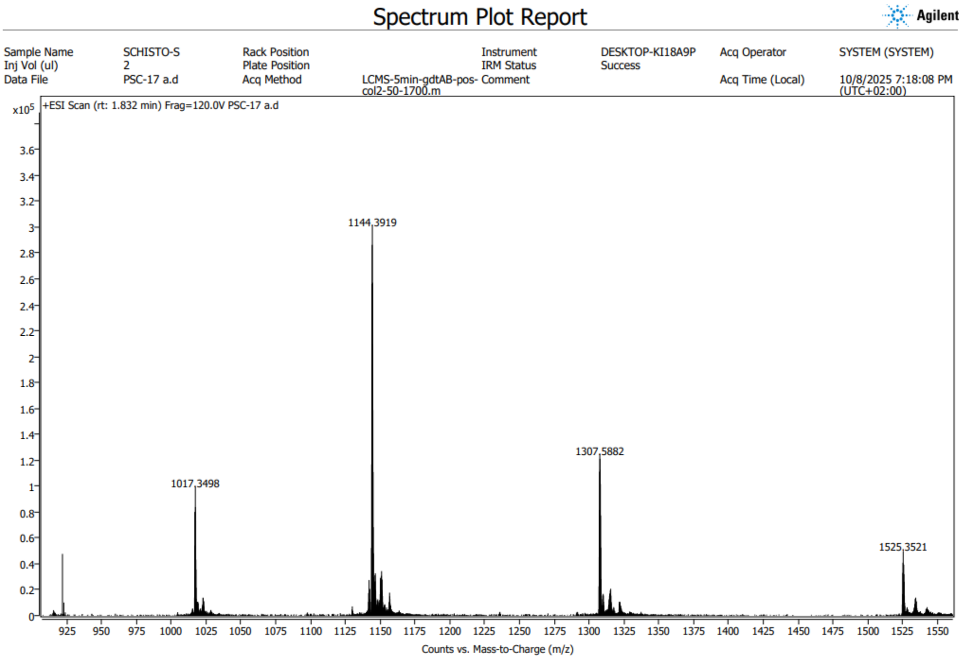

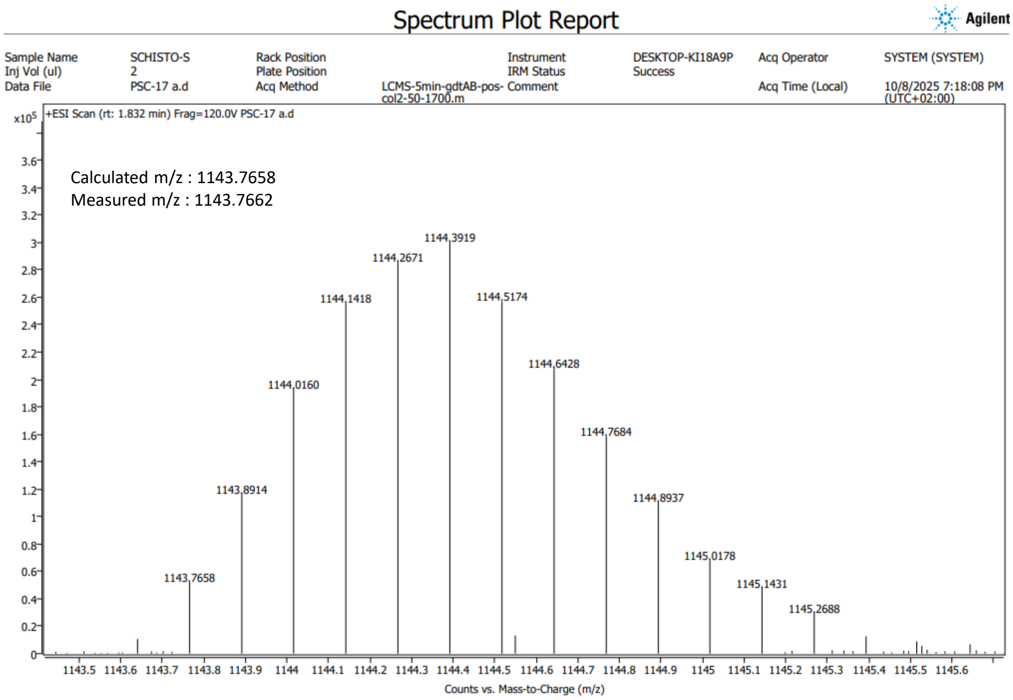

**Supplementary Figure 15.** HRMS analysis of synthetic schistosomin. HRMS data were aquired using An Orbitrap ID-X (Thermo) with ESI source coupled with an UPLC Vanquish (Thermo), with a Kinetex EVO C18 50 x 2.1 mm, 1.7µm column (Phenomenex). UV-Vis chromatogram is acquired on a 200-400 nm range. Mass spectra are acquired in a 100-2000 Da range in positive mode with a 120000 resolution. ESI, *m/z* for $[C_{386}H_{599}N_{115}O_{124}S_{10}]^{8+}$ calculated m/z for A+1 ion: 1143.7658, measured: 1143.7662

#### 3. Crystallization of the synthetic schistosomin and structure determination

Crystallization conditions for schistosomin at 10 mg/mL were screened using Qiagen kits (Valencia, CA, USA) with a Mosquito (SPT Labtech, Melbourn, UK). The crystals were manually reproduced in hanging drops experiments by mixing equal volumes of protein solution and precipitant solution mentioned in Table S1. The same crystallization condition led to two crystal forms (Table S1). Crystals were transferred to a cryo-protectant solution (mother liquor supplemented with 25% sucrose) and flash-frozen in liquid nitrogen.

Diffraction data were collected at 100 K on the PROXIMA 1 and 2 beamlines at synchrotron SOLEIL (Saint-Aubin, France). Diffraction intensities were integrated by the XDS program[4] using autoPROC (www.globalphasing.com) including STARANISO.[5] Both crystal structures were determined by molecular replacement with PHASER[6] using an AlphaFold2[7] model as a search model. Refinement of each structure was performed with BUSTER-2.10.4[8] using TLS group and NCS restraints when necessary. Inspection of the density maps and manual rebuilding were performed using COOT.[9] Refinement details of each structure are shown in Table 1.

**Supplementary Table 1.** Crystallographic data and refinement parameters for schistosomin.

| PDB code | **9RT6 (9FDM)** | **9FDO** |
|---|---|---|
| Crystallization conditions | 2 M AS/ 0.1 M Tris-HCl pH 8/ 0.2 M NaBr | |
| Space group<br>Cell parameters (Å,°) | $P2_1$<br>a= 21.8<br>b= 38.4<br>c=87.8<br>$\beta$= 91.2 | C2<br>a= 87.7<br>b= 38.0<br>c= 43.2<br>$\beta$= 95.3 |
| Resolution (Å) | 19.7-1.897 (2-1.897) | 43.6-2.056 (2.166-2.056) |
| Estimated resolution limit (Å) STARANISO | 1.8; 1.977 ; 1.937 | 2.122; 2.06; 2.055 |
| No. of observed reflections | 77793 (3325) | 43741 (856) |
| No. of unique reflections | 10239 (512) | 7585 (379) |
| Completeness (%) | 87.8 (30.3) | 84.5 (29.4) |
| $R_{merge}$ (%) | 26 (133.6) | 13.3 (115.8) |
| $R_{pim}$ (%) | 10 (56.6) | 5.8 (94.6) |
| $I/\sigma(I)$ | 5.6 (1.4) | 7.7 (0.8) |
| $CC_{1/2}$ | 0.985 (0.483) | 0.994 (0.313) |
| $R_{work}$ (%) | 20.7 | 22.9 |
| $R_{free}$ (%) | 25 | 27.5 |
| rms bond deviation (Å) | 0.008 | 0.008 |
| rms angle deviation (°) | 0.96 | 0.93 |
| Average B (Å$^2$)<br>Protein A/B<br>solvent | <br>18.7/27<br>25.7 | <br>33.5/35.5<br>35.2 |
| [a]Clashscore<br>MolProbity score | 1.68<br>1.46 | 0.82<br>1.51 |
| [a]Ramachandran plot (%)<br>Favoured<br>Outliers | <br>94.00<br>1.33 | <br>94.16<br>0.65 |

Values for the highest resolution shell are in parentheses. $CC_{1/2}$= percentage of correlation between intensities from random hall-dataset. [a]Calculated with MolProbity
STARANISO applies ellipsoidal mask: estimated resolution limits along the three crystallographic directions a*, b*, c*.

The atomic coordinates and structure factors have been deposited at the Protein Data Bank under 9RT6 and 9FDO for the synthetic schistosomin structures in P21 and C2 space group respectively.

A)

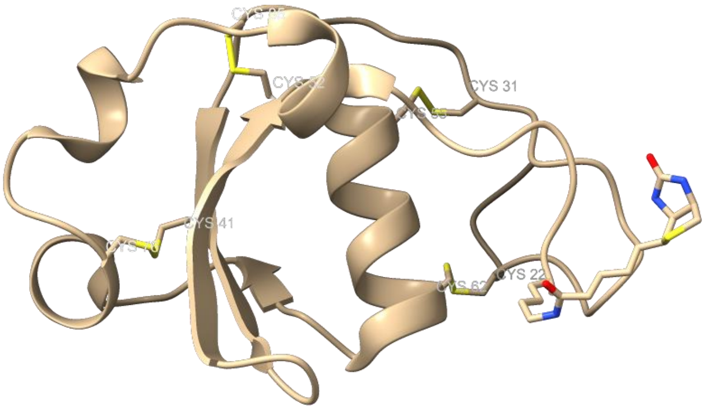

B)

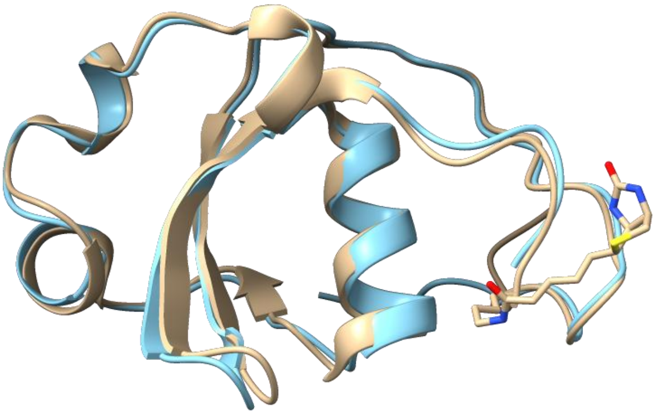

**Supplementary Figure 16.** X-ray structure of schistosomin. A) pdb entry 9RT6 showing the four disulfide bonds. B) X-ray structure 9RT6 (light orange) aligned with Alphafold2 model (blue). Pairwise structure alignment was performed using the structural alignment tool of the pdb.[10] The figures were created using ChimeraX.[11]

#### 4. Proscan analysis of synthetic schistosomin X-ray crystal structure (9FDO, chain A)

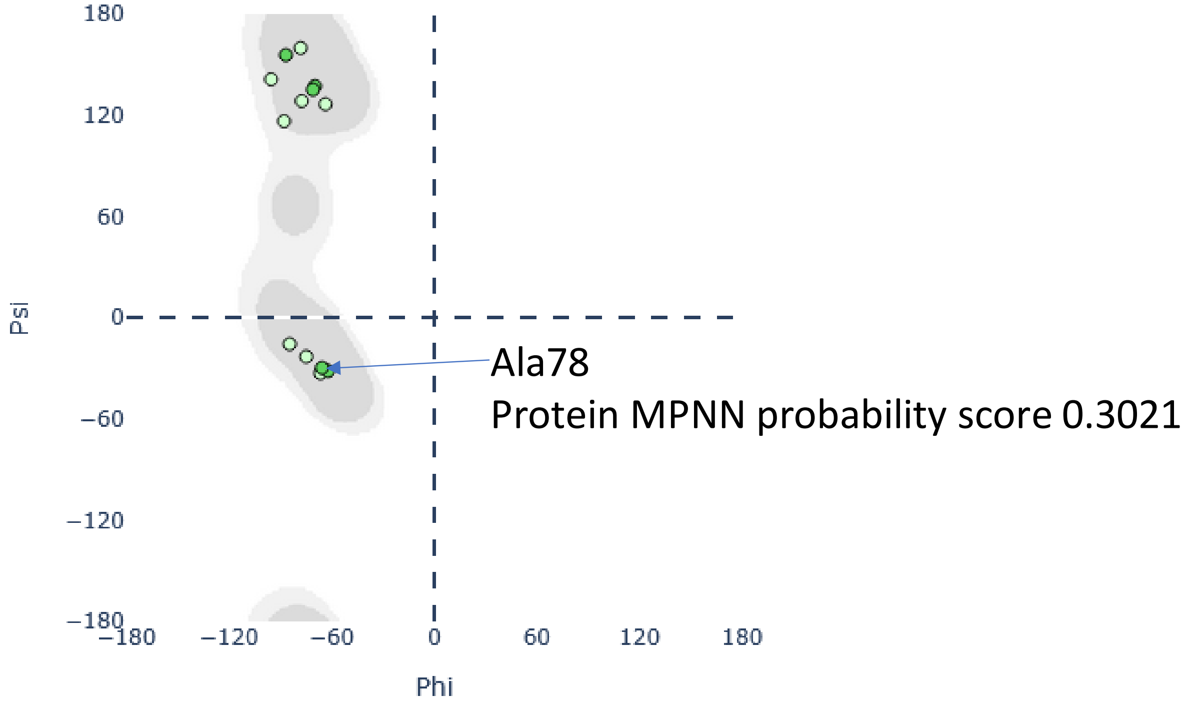

**Supplementary Figure 17.** Proscan analysis of schistosomin X-ray crystal structure (9FDO, chain A) yields residues compatible with XX →Pro mutation. Ramachandran plot showing input structure residue Φ/Ψ backbone angles plotted as circles, which are colored based on predicted proline favorability and score criteria. Only Proscan high-scoring residues are shown in this Ramachandran plot. A78 is indicated by an arrow. The figure was prepared using output data from Proscan server (https://proscan.ibbr.umd.edu/).[12]

| Residue | ID | Chain | Phi | Psi | Sec Struct | ProteinMPNN Probability | Pro Angle | Pre-Pro Angle | Notes |
|---|---|---|---|---|---|---|---|---|---|
| GLU | 75 | A | -65.54 | -39.283 | A_Helix | 0.0032 | Preferable | Preferable | H-bond_A0043 H-bond_A0071 |
| CYS | 76 | A | -105.562 | -96.265 | Turn | 0.0034 | Questionable | Preferable | H-bond_A0072 Disulfide_A0041 |
| ARG | 77 | A | -146.529 | 163.154 | No_Sec | 0.0071 | Questionable | Questionable | - |
| ALA | 78 | A | -65.753 | -29.843 | Turn | 0.3021 | Preferable | Preferable | - |
| ASP | 79 | A | -103.713 | 38.895 | Turn | 0.004 | Acceptable | Preferable | - |
| TRP | 80 | A | -61.499 | -29.466 | 3-10_Helix | 0.0034 | Preferable | Questionable | H-bond_A0077 |
| ALA | 81 | A | -79.848 | -19.025 | 3-10_Helix | 0.0057 | Preferable | Preferable | - |
| SER | 82 | A | -91.633 | 4.231 | 3-10_Helix | 0.0043 | Preferable | Acceptable | H-bond_A0079 |
| THR | 83 | A | -58.482 | 131.55 | Poly_Proline | 0.004 | Preferable | Questionable | H-bond_A0080 |

**Supplementary Figure 18.** Details of Proscan output centered on Ala78. The figure was prepared using output data from Proscan server.[12]

#### 5. Analysis of the synthetic schistosomin by circular dichroism and nanoDSF

The concentration of synthetic BgSminmine (BgSmin_A78) solubilized in 10 mM sodium phosphate buffer (pH 7.2) was determined using a NanoDrop spectrophotometer by measuring absorbance at 280 nm, based on its specific extinction coefficient (ε = 10470 $L \cdot mol^{-1} \cdot cm^{-1}$). The stock solution (~4 mg/mL) was diluted in the same buffer to a final concentration of 0.1 mg/mL for circular dichroism (CD) analysis.

CD spectra were recorded on a Jasco J-815 spectropolarimeter coupled with PFD-425S/15 pelletier heating unit at 25 °C over the wavelength range of 185-260 nm, using a 0.1 cm path-length quartz cuvette (HELMA). Spectra were acquired with a bandwidth of 2 nm, a data pitch of 1 nm, and an integration time of 1 s. Each spectrum corresponds to the average of eight accumulations, and data were smoothed using a five-point smoothing algorithm.

CD data are presented as mean residue weight ellipticity [Θ]MRW (mean ± 95% confidence interval, ((mDeg x (MW / nAA-1)) / (10 x $mg.mL^{-1}$ x l), with MW= 10629 $g.mol^{-1}$, n=80, l=0.1 cm)as a function of wavelength between 185 and 260 nm [12].

Thermal denaturation experiments were carried out using the same acquisition parameters, using six accumulations for averaging and smoothing. The temperature was increased from 15 °C to 90 °C in 5 °C increments, with a 1-min equilibration time at each temperature prior to data acquisition. Ellipticity values at 208 and 222 nm were monitored and plotted as a function of temperature. The melting temperature (Tm) was determined by fitting the data using single- or double-slope four-parameter sigmoid regression models implemented in SigmaPlot v14.5.

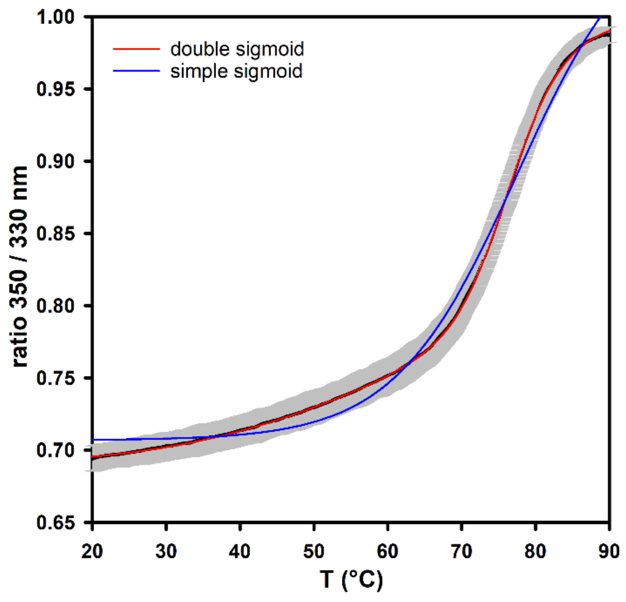

**Supplementary Figure 19**. Comparison between simple and double sigmoidal fittings of nanoDSF signals

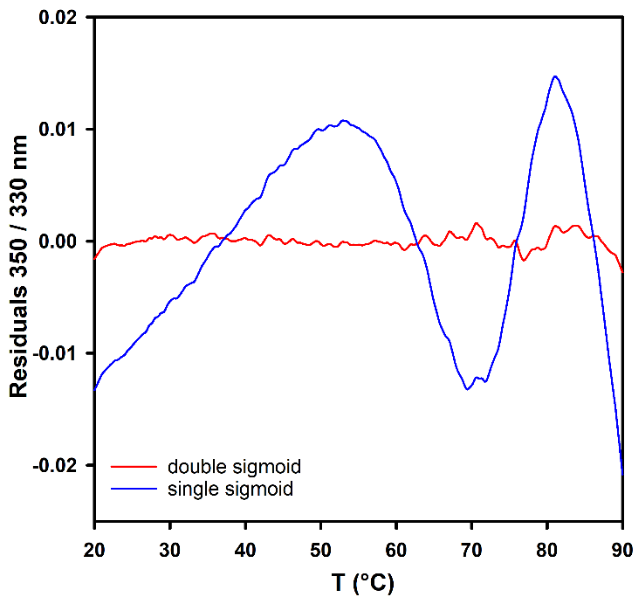

**Supplementary Figure 20**. Residual of simple and double sigmoidal fittings of nanoDSF signals

#### 6. Proteomic analysis of synthetic schistosomin

Stock solutions

A total of 116 µg of synthetic schistosomin was dissolved in 58.8 µL of 20 mM ammonium bicarbonate (ABC) buffer, pH 8.0, to obtain a 2 mg/mL stock solution. 12 µg of the A-isoform of isolated schistosomin was dissolved in 12 µL of 20 mM ABC buffer to prepare a 1 mg/mL stock solution. Similarly, 6 µg of the P-isoform of isolated schistosomin was dissolved in 6 µL of 20 mM ABC buffer to generate a 1 mg/mL stock solution.

Schistosomine digestion

From stock solutions, 10 µL of a 0.1 mg/mL protein solution were prepared in 20 mM ammonium bicarbonate (ABC) buffer. Then, 0.5 µL of a 0.1 mg/mL trypsin solution was added to achieve a 1:20 trypsin-to-protein ratio, and the mixture was incubated overnight at 37 °C with agitation.

To stop the digestion, 1 µL of a 1% trifluoroacetic acid (TFA) solution was added. The resulting samples were analyzed in MALDI mass spectrometry.

MALDI mass analysis

On a small AnchorChip Bruker MALDI plate, a 0.6 µL droplet of α-cyano-4-hydroxycinnamic acid (CHCA) solution (10 mg/mL in 50% acetonitrile with 0.1% TFA) was deposited as the matrix. Subsequently, 0.6 µL of the analyte solution was deposited on top of the matrix spot. After co-crystallization, the samples were analyzed using MALDI-MS, in positive mode with refletron.

Apparatus :

MALDI data were acquired on a rapifleX MALDI-TOF/TOF mass spectrometer (Bruker Daltonics, Billerica, MA, USA) equipped with a smartbeam 3D laser operating at 10 kHz. The instrument was operated in positive ion mode, and data were acquired in MS The mass range typically covered m/z 500–8000, suitable for peptide and small protein analysis. Spectra were recorded using flexControl software, and data processing was performed with flexAnalysis (Bruker Daltonics). Sequence (Bruker Daltonics) was used to generate theoretical fragment with multiples disulfide bridges combinations, Biotools (Bruker Daltonics) was used for identifications and attibution of peaks.

#### 7. Molecular dynamics simulations

##### 7.1. Starting structures

The starting structure of BgSmin$^{A}$ D88E (Schistosomin A78 isoform) used for the molecular dynamics simulations was derived from the X-ray crystal structure of BgSmin$^{A\ Synth}$ (space group C2, chain B). This structure was selected because it is the only experimentally determined X-ray structure among those reported in this work that includes all N-terminal residues. The C-terminal Lys(Biot)-$NH_2$ residue was removed using ChimeraX.

The starting BgSmin$^{P}$ D88E structure was generated from the BgSmin$^{A}$ D88E model by substituting Ala78 with Pro78 using ChimeraX.

##### 7.2 Molecular dynamics simulations

Molecular dynamics simulations were performed using the GROMACS/2023.4[13] and GROMACS/2023.4-mpi-cuda package of programs with the Charmm36 force field[14] and tip3p water model. The molecular dynamic simulations were performed in triplicate.

The protein was placed in a dodecahedral simulation box with a minimum distance of 2.0 nm between the solute and the box boundaries (box volume ≈ 367 $nm^3$). The system was solvated with water and subsequently neutralized and adjusted to an ionic strength of 70 mM by adding NaCl. This ionic strength was chosen to match reported values for the hemolymph of freshwater pulmonate mollusks (*Lymnaea stagnalis* and *Lymnaea truncatula*), which belong to the same Heterobranchia group as *Biomphalaria glabrata*.

The N-terminus was protonated ($NH_3^+$), while the C-terminus was treated as deprotonated ($COO^-$). The system was energy-minimized until the maximum force was below 1000 $kJ \cdot mol^{-1} \cdot nm^{-1}$ and subsequently equilibrated. Temperature (300 K, 338 K or 345 K) and pressure (1 bar) equilibration were performed under position restraints for 400 ps.

Production molecular dynamics simulations were carried out under NPT conditions using the velocity-rescale thermostat and the Parrinello–Rahman (C-rescale) barostat to maintain constant temperature and pressure (T = 300 K, $\tau_T$ = 0.1 ps; P = 1 bar, $\tau_P$ = 2 ps). Van der Waals interactions were treated using a force-switch cutoff scheme with a cutoff distance of 1.2 nm and a switching distance of 1.0 nm. Long-range electrostatic interactions were calculated using the particle mesh Ewald (PME) method with a Fourier grid spacing of 0.12 nm, interpolation order 4, and a relative tolerance of $10^{-5}$.

A time step of 2 fs was used. Coordinates were saved every 10 ps, and each simulation replicate was run for 160 ns.

##### 7.3 BgSmin$^A$ D88E

###### 7.3.1 Root-mean-square deviation (RMSD)

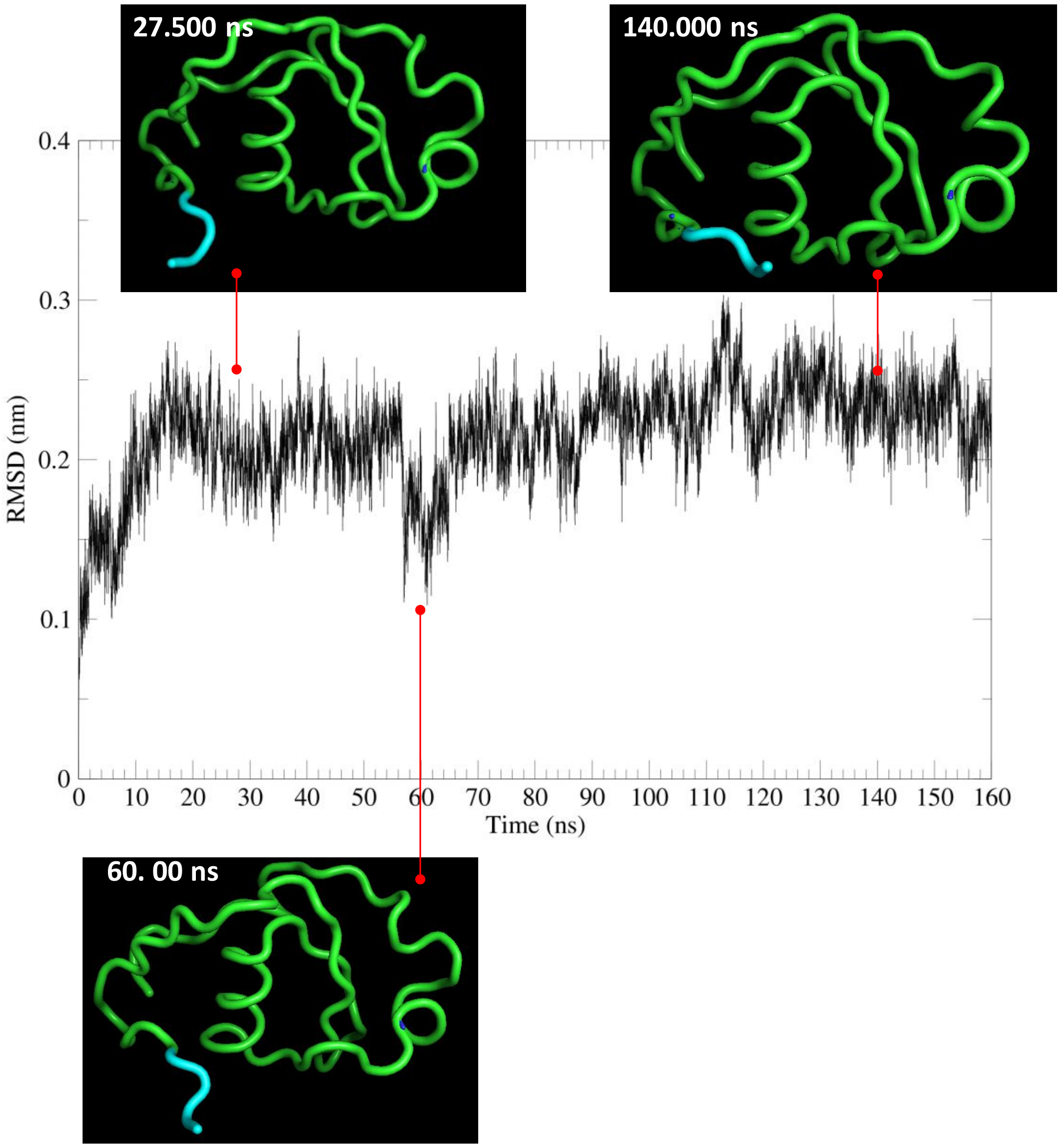

**Supplementary Figure 21.** Time evolution of the root-mean-square deviation (RMSD) of BgSmin$^A$ D88E backbone atoms for trajectory 1. Temperature: 300 K

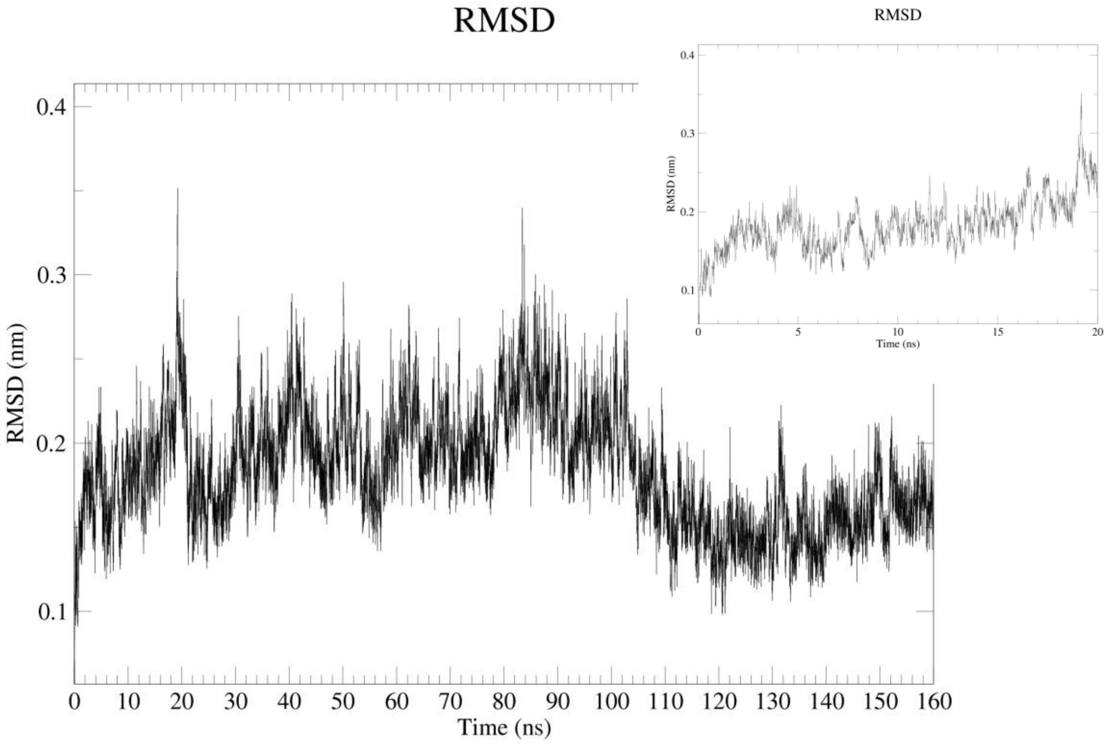

**Supplementary Figure 22.** Time evolution of the root-mean-square deviation (RMSD) of $BgSmin^A$ D88E backbone atoms for trajectory 2. Temperature: 300 K

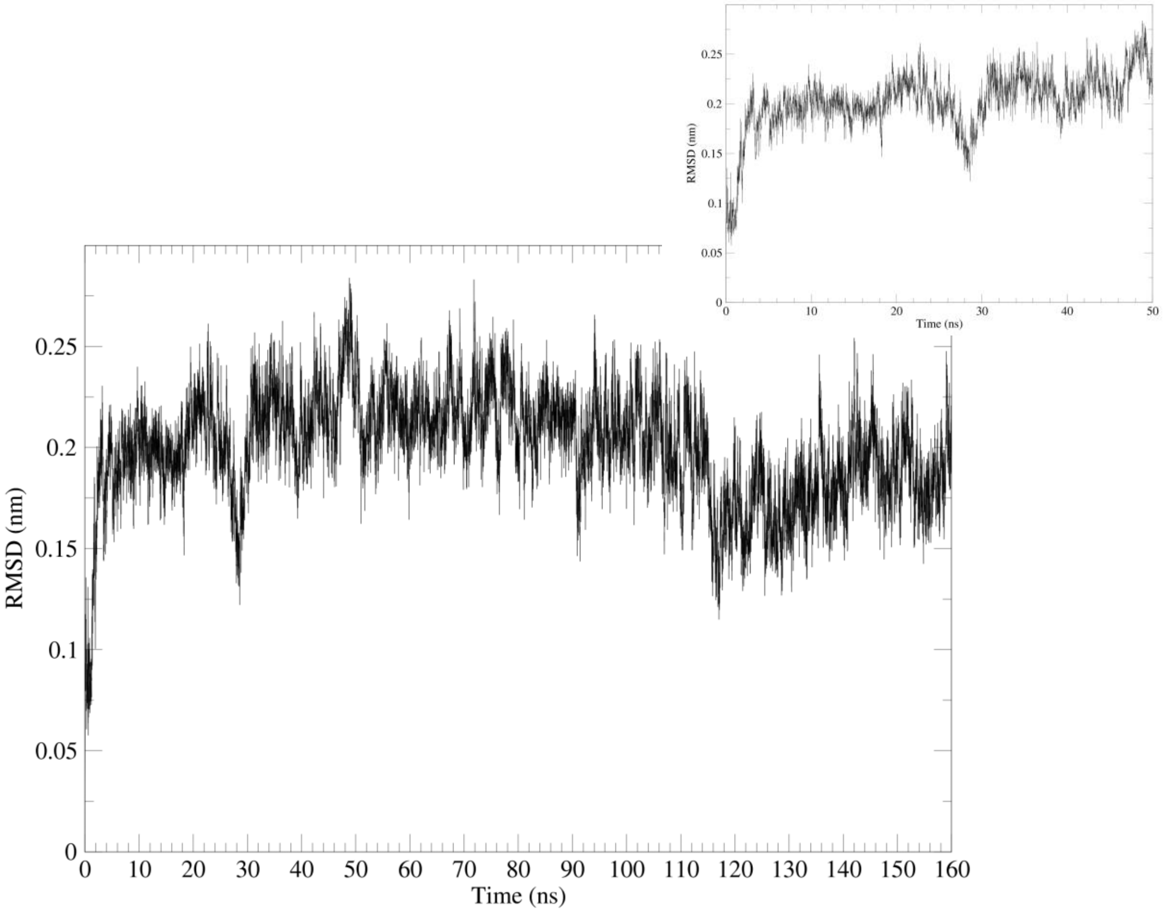

**Supplementary Figure 23.** Time evolution of the root-mean-square deviation (RMSD) of BgSmin[A] D88E backbone atoms for trajectory 3. Temperature: 300 K

**7.3.2** Radius of gyration

A)

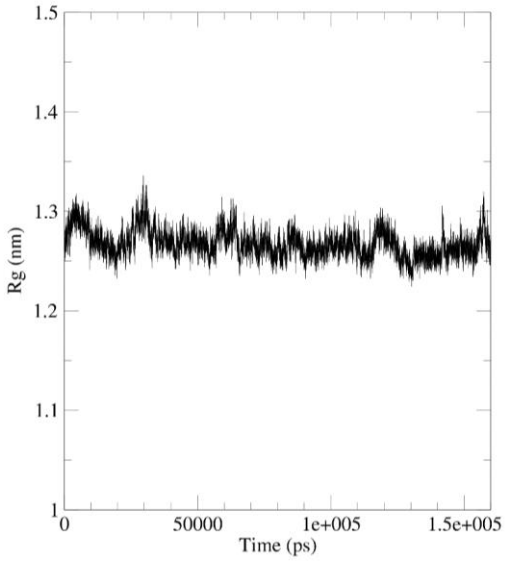
1.5
1.4
1.3
1.2
1.1
1
Rg (nm)
0
50000
1e+005
1.5e+005
Time (ps)

B)

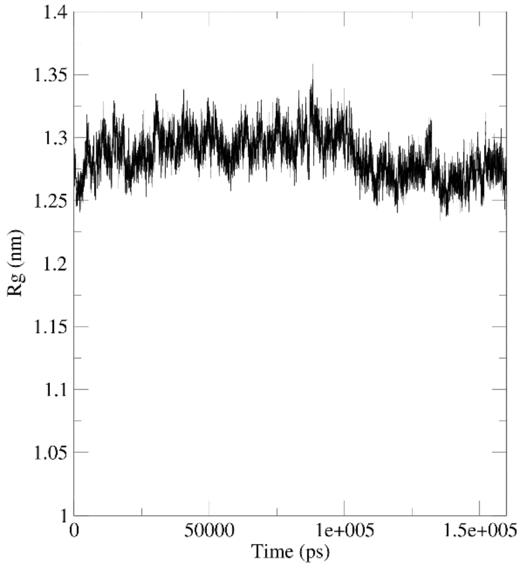
1.4
1.35
1.3
1.25
1.2
1.15
1.1
1.05
1
Rg (nm)
0
50000
1e+005
1.5e+005
Time (ps)

C)

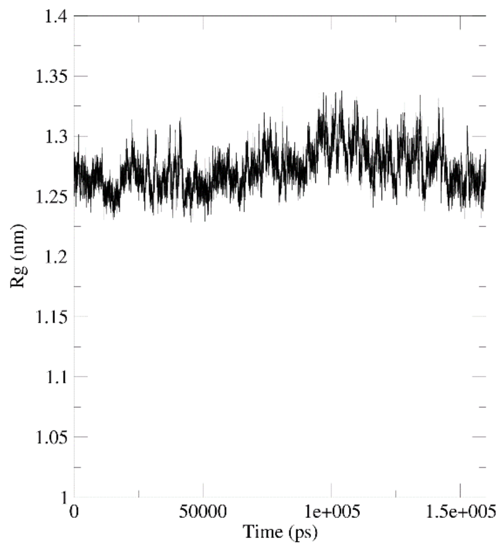

**Supplementary Figure 24.** Time evolution of the radius of gyration of BgSmin$^A$ D88E during trajectory 1 (A), trajectory 2 (B) and trajectory 3 (C), temperature 300 K.

A)

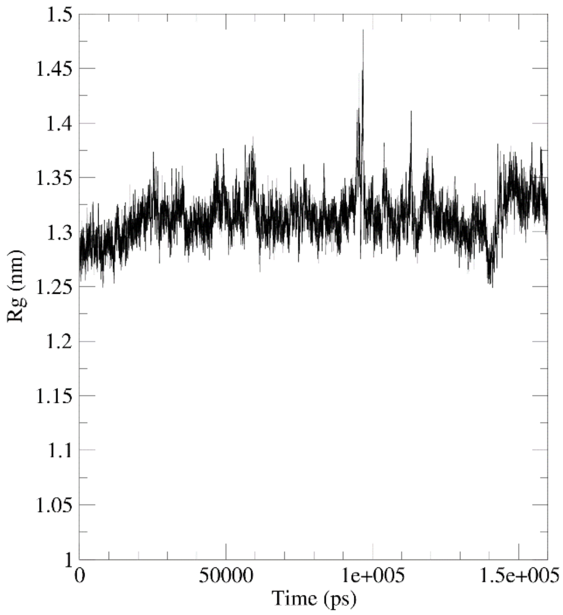

B)

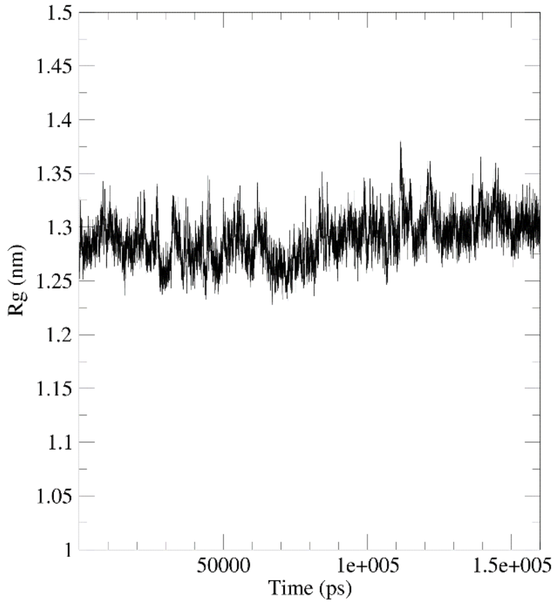

C)

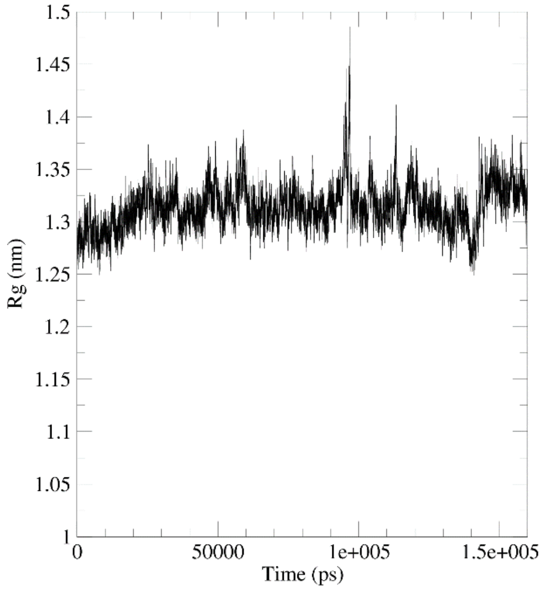

**Supplementary Figure 25.** Time evolution of the radius of gyration of BgSmin$^{A}$ D88E during trajectory 1 (A), trajectory 2 (B) and trajectory 3 (C), temperature 345 K.

**7.3.3** Root-mean-square fluctuation (RMSF)

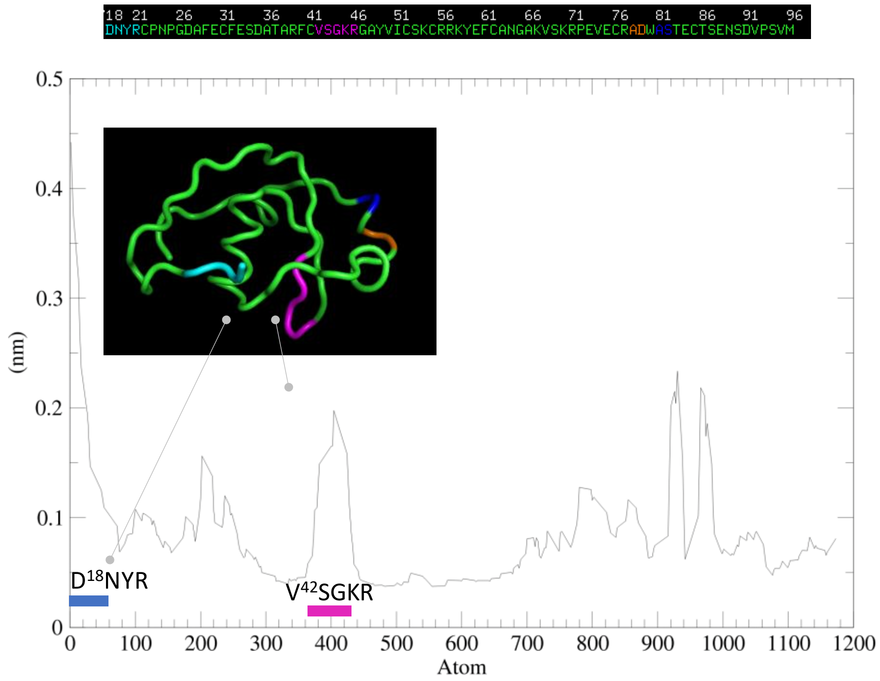

A)

B)

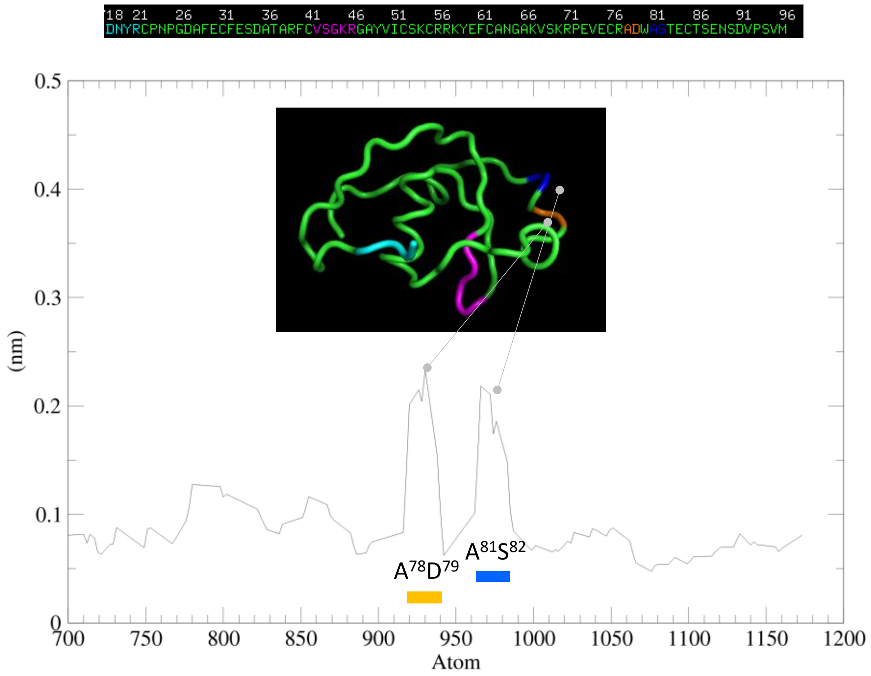

**Supplementary Figure 26.** Root-mean-square fluctuation (RMSF) of BgSmin$^{A}$ D88E backbone atoms for trajectory 1 (15-160 ns). (A) All backbone atoms. (B) Backbone atoms from atom index 700 onward. Temperature: 300 K.

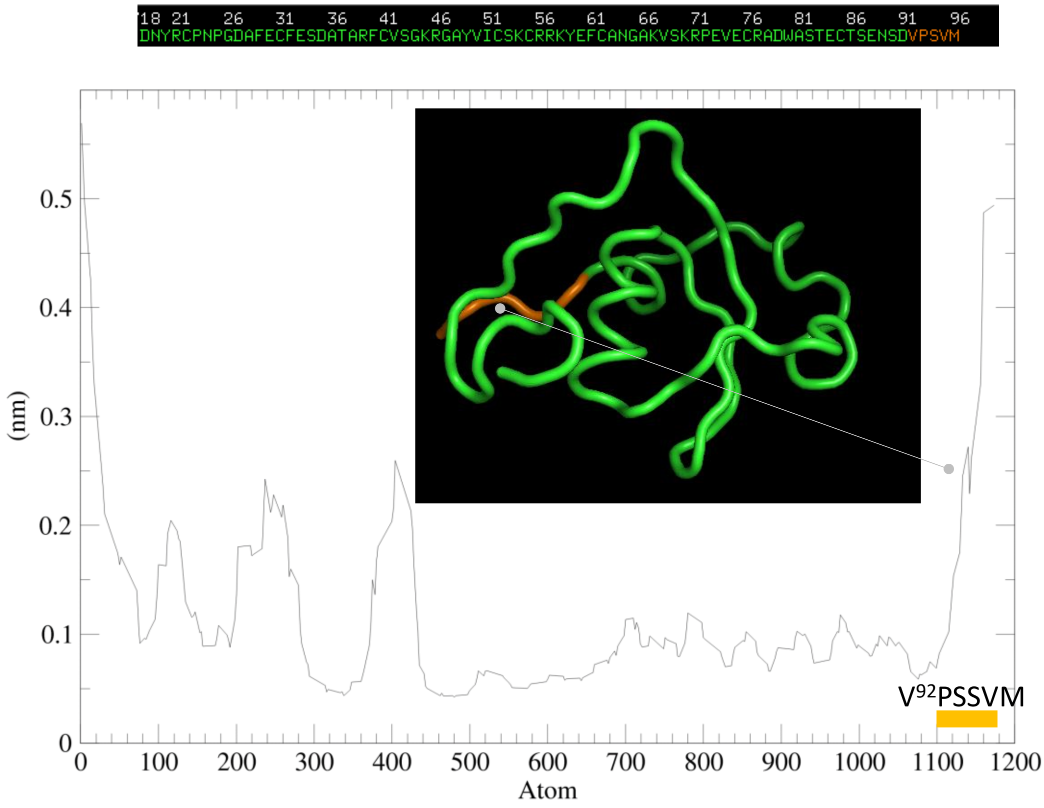

**Supplementary Figure 27.** Root-mean-square fluctuation (RMSF) of BgSmin$^A$ D88E backbone atoms for trajectory 2 (2-160 ns). All backbone atoms, temperature 300 K.

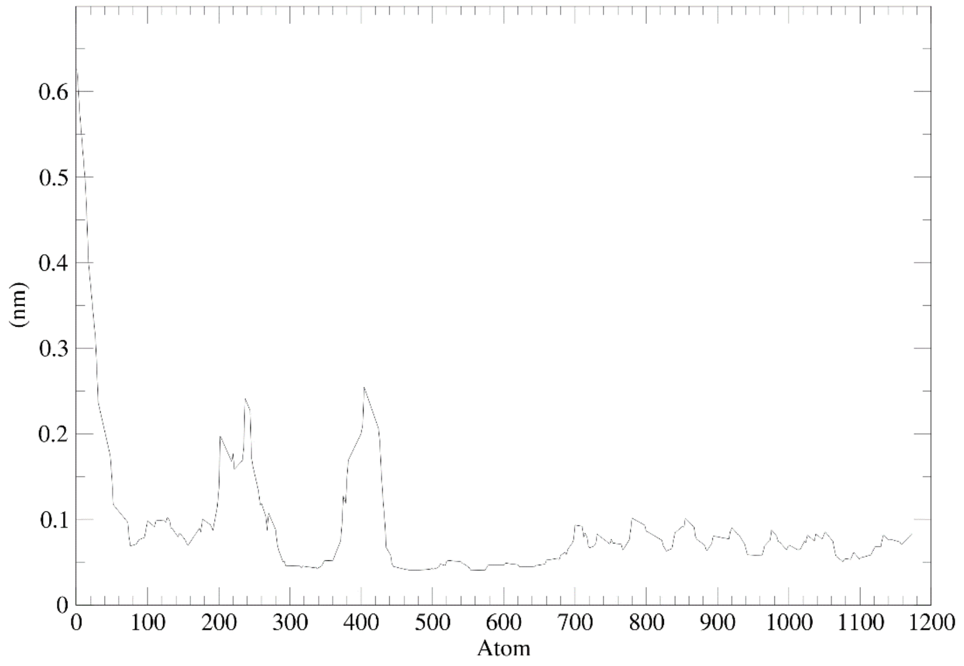

**Supplementary Figure 28.** Root-mean-square fluctuation (RMSF) of BgSmin$^A$ D88E backbone atoms for trajectory 3 (2-160 ns). All backbone atoms, temperature 300 K.

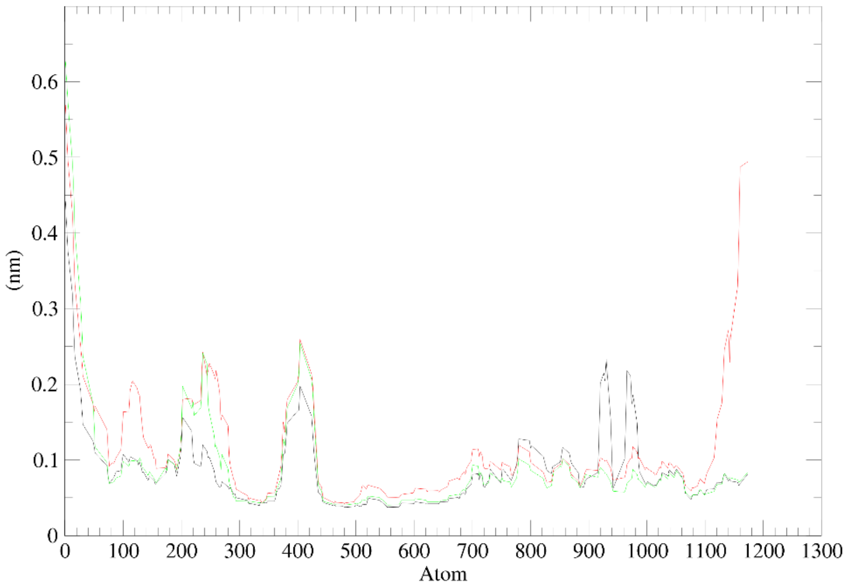

**Supplementary Figure 29.** Root-mean-square fluctuation (RMSF) of BgSmin$^{A}$ D88E backbone atoms for all trajectories at 300 K. 15-160 ns trajectory 1 (black), 2-160 ns trajectories 2 (red) and 3 (green). All backbone atoms.

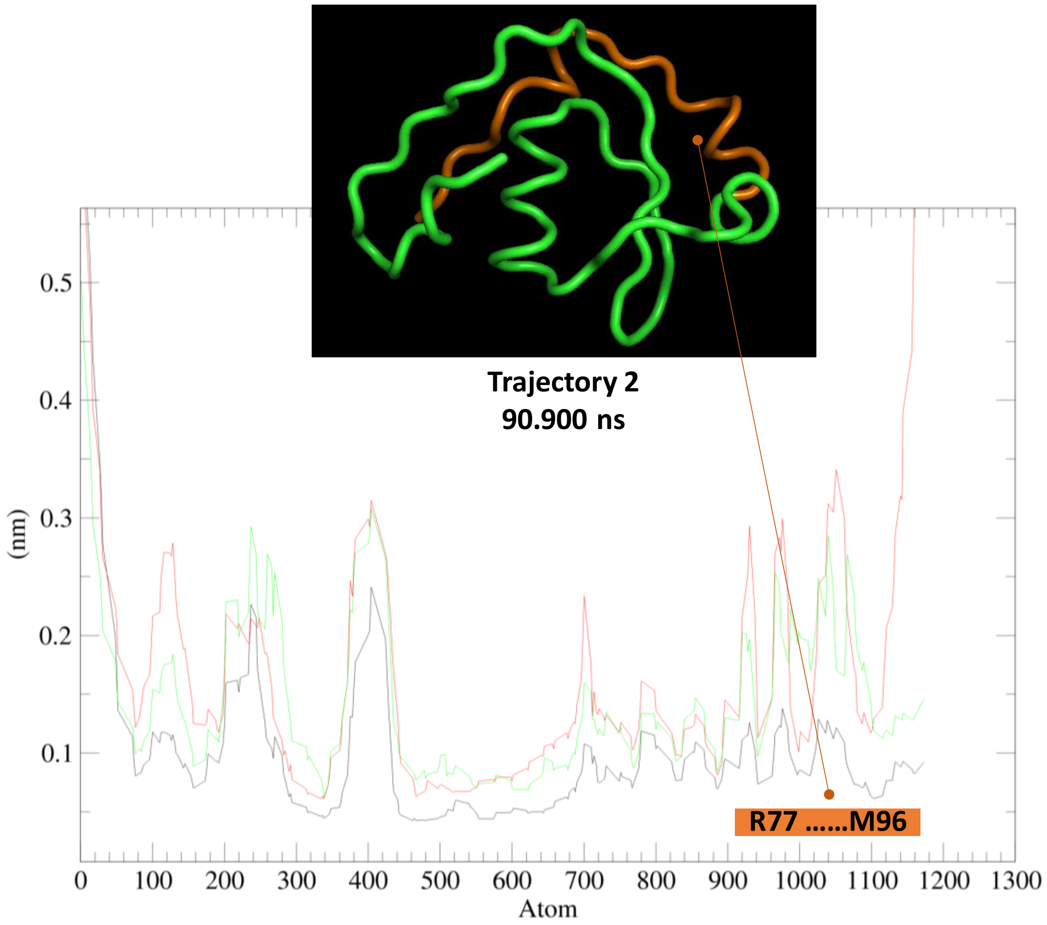

**Supplementary Figure 30.** Root-mean-square fluctuation (RMSF) of BgSmin$^{A}$ D88E backbone atoms for all trajectories an 338 K. 2-160 ns for all trajectories, trajectory 1 (black), trajectory 2 (red) and trajectory 3 (green). All backbone atoms.

**Starting structure**

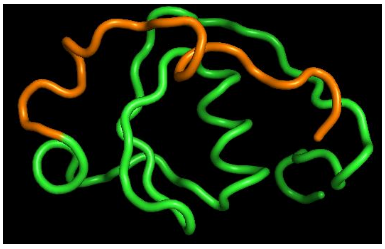

**Trajectory 2**

42.800 ns 146.000 ns 160.000 ns

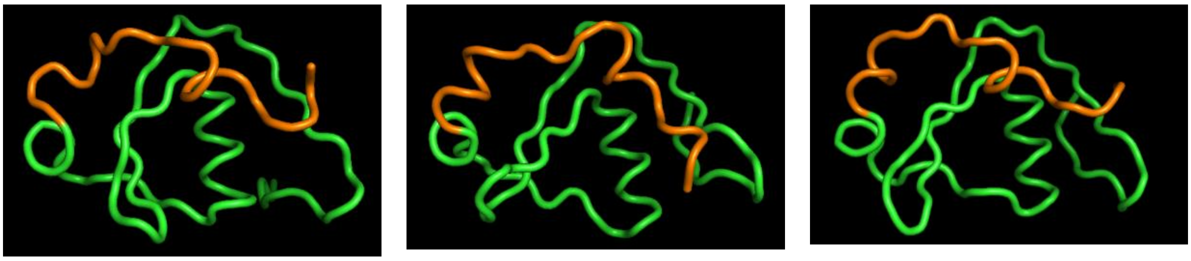

**Trajectory 3**

87.700 ns | 14.810 ns | 160.000 ns

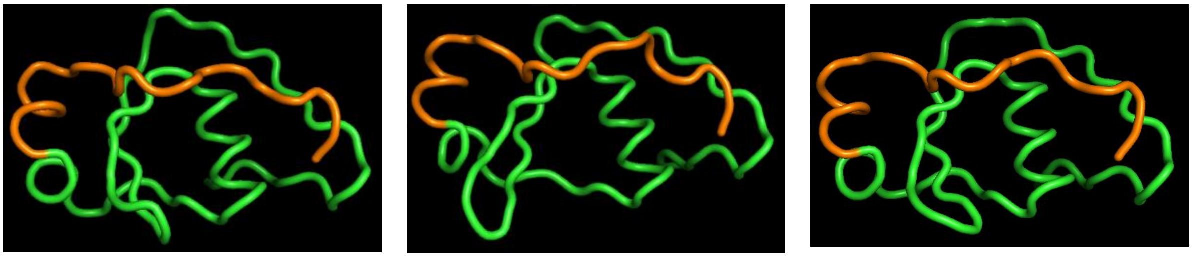

**Supplementary Figure 31.** Representative structural snapshots from trajectories 2 and 3 at 338 K at the indicated time points. The region spanning residues Arg77 to Met96 is shown in orange (cartoon representation).

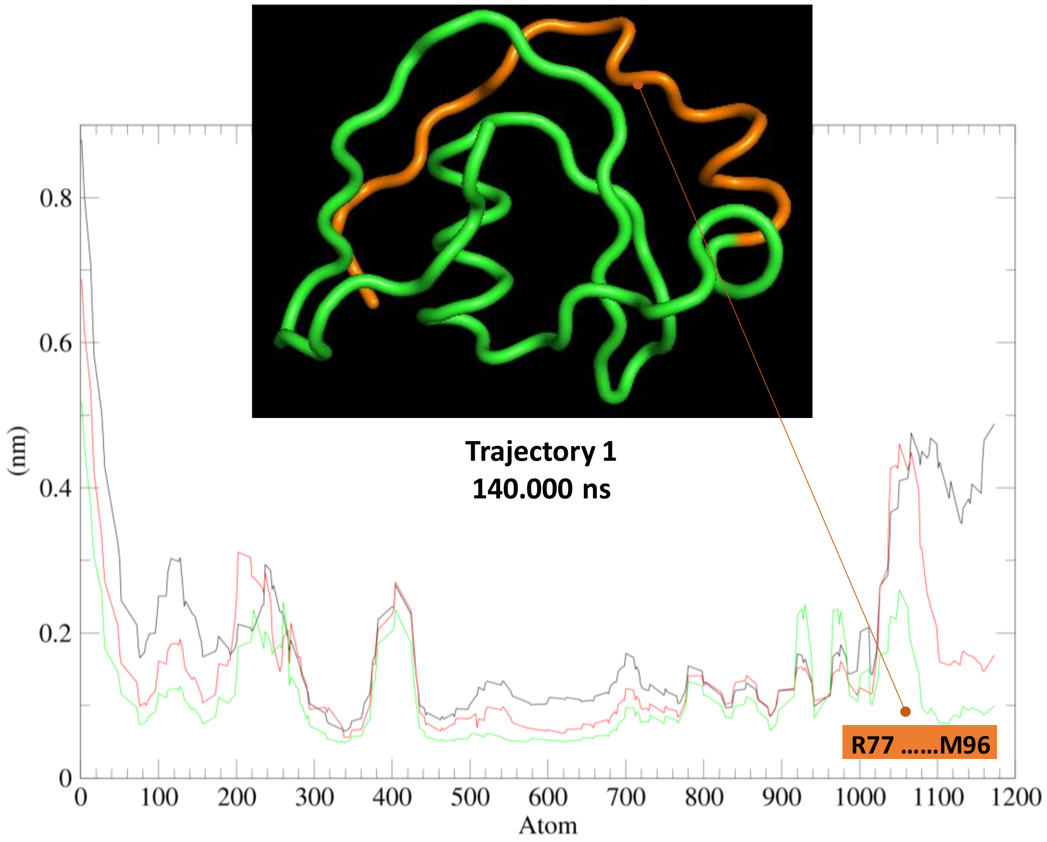

**Supplementary Figure 32.** Root-mean-square fluctuation (RMSF) of BgSmin$^{A}$ D88E backbone atoms for all trajectories an 345 K. 2-160 ns for all trajectories, trajectory 1 (black), trajectory 2 (red) and trajectory 3 (green). All backbone atoms.

**Starting structure**

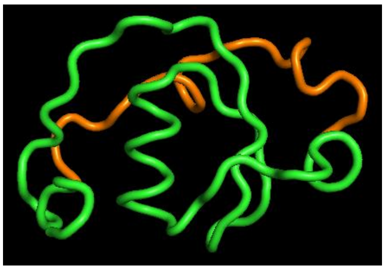

**Trajectory 1**

28.000 ns | 96.800 ns | 96.800 ns

Rotated 90 °

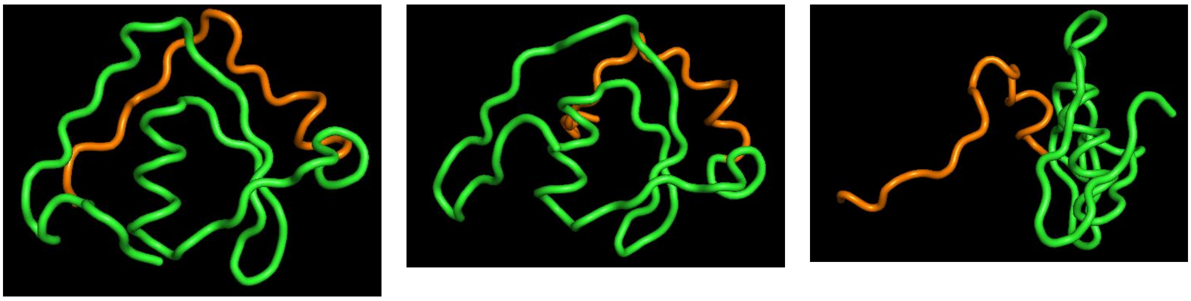

**Trajectory 2**

95.400 ns | 10.930 ns | 160.000 ns

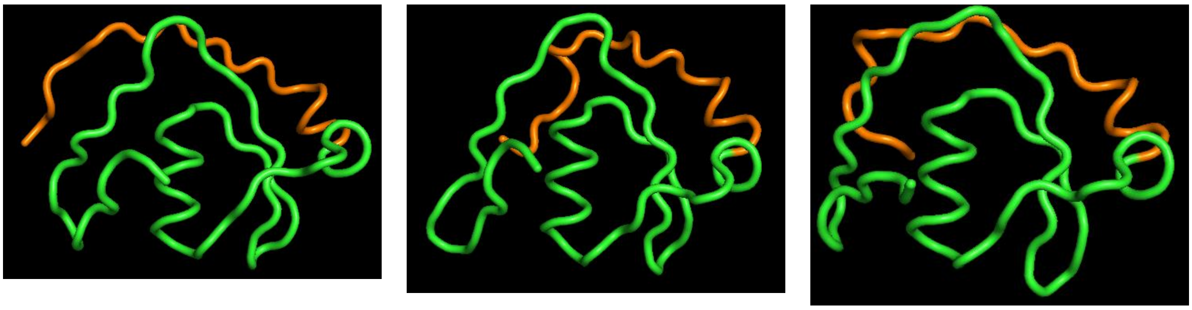

**Supplementary Figure 33.** Representative structural snapshots from trajectories 1 and 2 at 345 K at the indicated time points. The region spanning residues Arg77 to Met96 is shown in orange (cartoon representation).

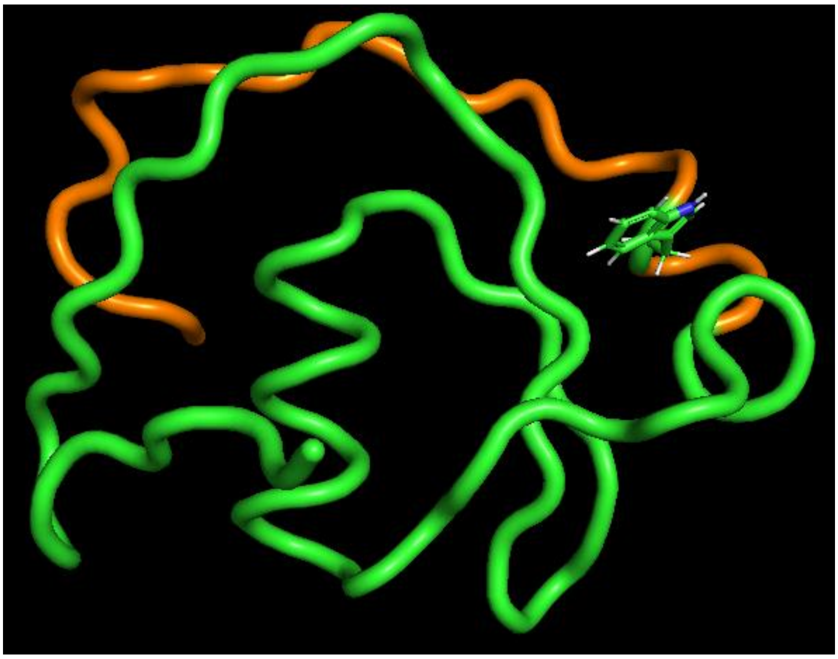

**Supplementary Figure 34.** Snapshot of trajectory 1 (345 K, 160.000 ns) showing W80 (atoms 940-963).

**7.3.4** Ramachandran plots

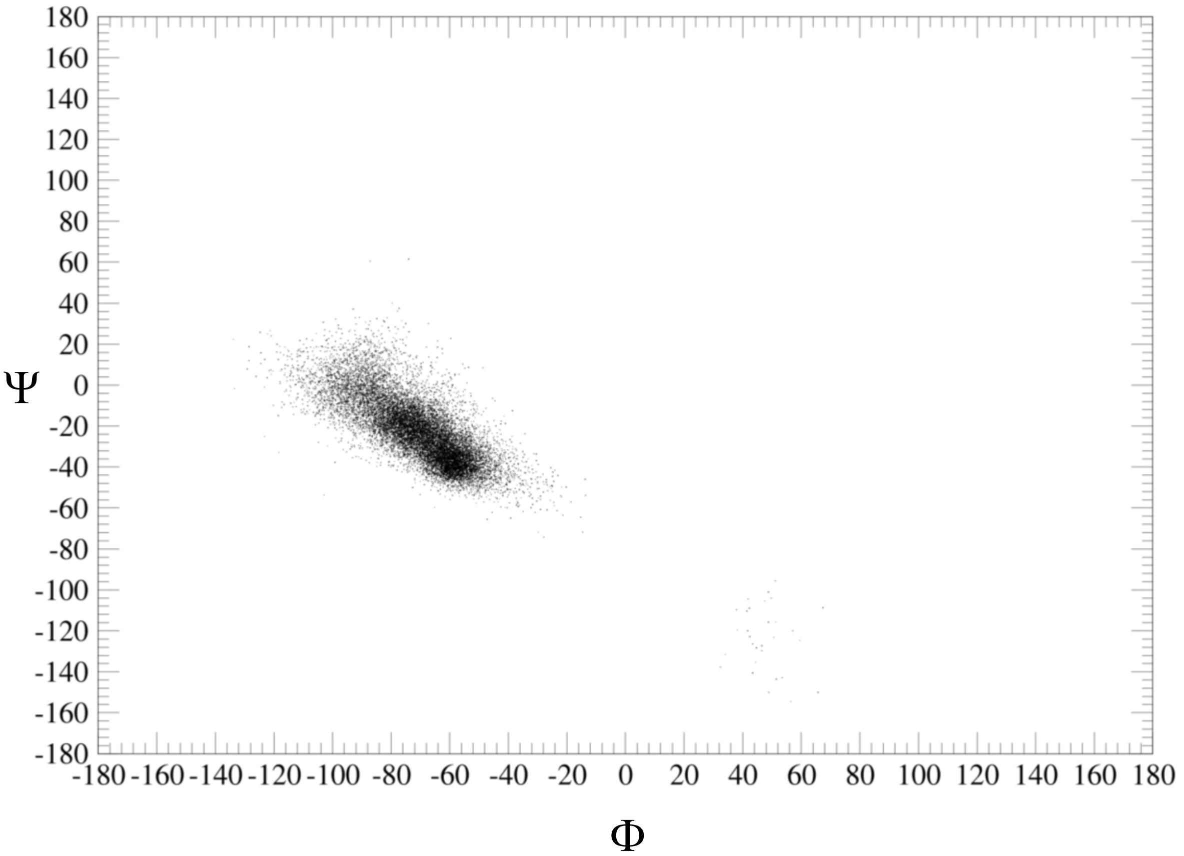

**Supplementary Figure 35.** Ramachandran plot for Ala78 over trajectory 1, 300 K.

**7.3.5 Secondary structure analysis (DSSP)**

Secondary structure analysis of the trajectories has been performed with the DSSP algorithm implemented in gromacs/2024.3-mpi-cuda- cufftmp (hmode dssp, hbond geometry).

**Supplementary Table 2.** DSSP analysis of BgSmin$^{A}$ D88E at 300 K.

| | Loop | Breaks | Bends | Turns | PP helices | π–helices | 3/10-helices | β-strands | β-bridges | α-helices |
|---|---|---|---|---|---|---|---|---|---|---|
| **Trajectory 1** | 33,0 | 0 | 15,5 | 10,5 | 4,8 | 0 | 2,9 | 13,3 | 3,1 | 16,8 |
| **Trajectory 2** | 30,9 | 0 | 17,5 | 10,5 | 7,5 | 0 | 2,2 | 13,3 | 1,6 | 16,4 |
| **Trajectory 3** | 33,4 | 0 | 16,7 | 9,6 | 4,5 | 0 | 2,7 | 14,1 | 2,8 | 16,2 |
| **Mean** | 32,2 | 0,0 | 16,6 | 10,2 | 5,6 | 0,0 | 2,6 | 13,6 | 2,5 | 16,5 |
| **SD** | 1,8 | 0,0 | 1,0 | 0,5 | 1,7 | 0,0 | 0,4 | 0,5 | 0,8 | 0,3 |
| **SE (95%)** | 4,4 | 0,0 | 2,5 | 1,3 | 4,1 | 0,0 | 0,9 | 1,1 | 2,0 | 0,8 |

**Supplementary Table 3.** DSSP analysis of BgSmin$^{A}$ D88E at 338 K.

| | Loop | Breaks | Bends | Turns | PP helices | π–helices | 3/10-helices | β-strands | β-bridges | α-helices |
|---|---|---|---|---|---|---|---|---|---|---|
| **Trajectory 1** | 32,9 | 0,0 | 17,6 | 9,9 | 5,1 | 0,0 | 2,5 | 13,6 | 2,8 | 15,6 |
| **Trajectory 2** | 34,3 | 0,0 | 16,6 | 9,8 | 7,5 | 0,0 | 1,8 | 11,7 | 1,4 | 16,8 |
| **Trajectory 3** | 37,1 | 0,0 | 15,6 | 8,4 | 5,3 | 0,0 | 1,1 | 11,1 | 2,6 | 18,8 |
| **Mean** | 34,8 | 0,0 | 16,6 | 9,4 | 6,0 | 0,0 | 1,8 | 12,1 | 2,3 | 17,1 |
| **SD** | 2,1 | 0,0 | 1,0 | 0,8 | 1,4 | 0,0 | 0,7 | 1,3 | 0,7 | 1,6 |
| **SE (95%)** | 5,3 | 0,0 | 2,4 | 2,1 | 3,4 | 0,0 | 1,8 | 3,3 | 1,8 | **4,0** |

**Supplementary Table 4.** DSSP analysis of BgSmin$^{A}$ D88E at 345 K.

| | Loop | Breaks | Bends | Turns | PP helices | π–helices | 3/10-helices | β-strands | β-bridges | α-helices |
|---|---|---|---|---|---|---|---|---|---|---|
| **Trajectory 1** | 37,2 | 0,0 | 16,7 | 7,9 | 5,2 | 0,0 | 0,5 | 12,5 | 1,9 | 18,1 |
| **Trajectory 2** | 35,4 | 0,0 | 18,5 | 8,6 | 4,8 | 0,0 | 1,6 | 12,7 | 2,5 | 15,8 |
| **Trajectory 3** | 37,2 | 0,0 | 16,7 | 7,9 | 5,2 | 0,0 | 0,5 | 12,5 | 1,9 | 18,1 |
| **Mean** | 36,6 | 0,0 | 17,3 | 8,1 | 5,1 | 0,0 | 0,9 | 12,6 | 2,1 | 17,3 |
| **SD** | 1,0 | 0,0 | 1,0 | 0,4 | 0,2 | 0,0 | 0,6 | 0,1 | 0,3 | 1,3 |
| **SE (95%)** | 2,6 | 0,0 | 2,6 | 1,0 | 0,5 | 0,0 | 1,6 | 0,2 | 0,8 | 3,2 |

##### 7.4 BgSmin$^{P}$ D88E

Across all simulations, both isoforms rapidly reached stable conformational states, as reflected by the early convergence of backbone root-mean-square deviation (RMSD) values and the absence of large-scale structural rearrangements over the full simulation time (Figures S19-S21 and S29-S31). Consistently, the radius of gyration showed limited variation throughout the trajectories (Figures S22 and S32), indicating preservation of the compact global fold.

###### **7.4.1** Root-mean-square deviation (RMSD)

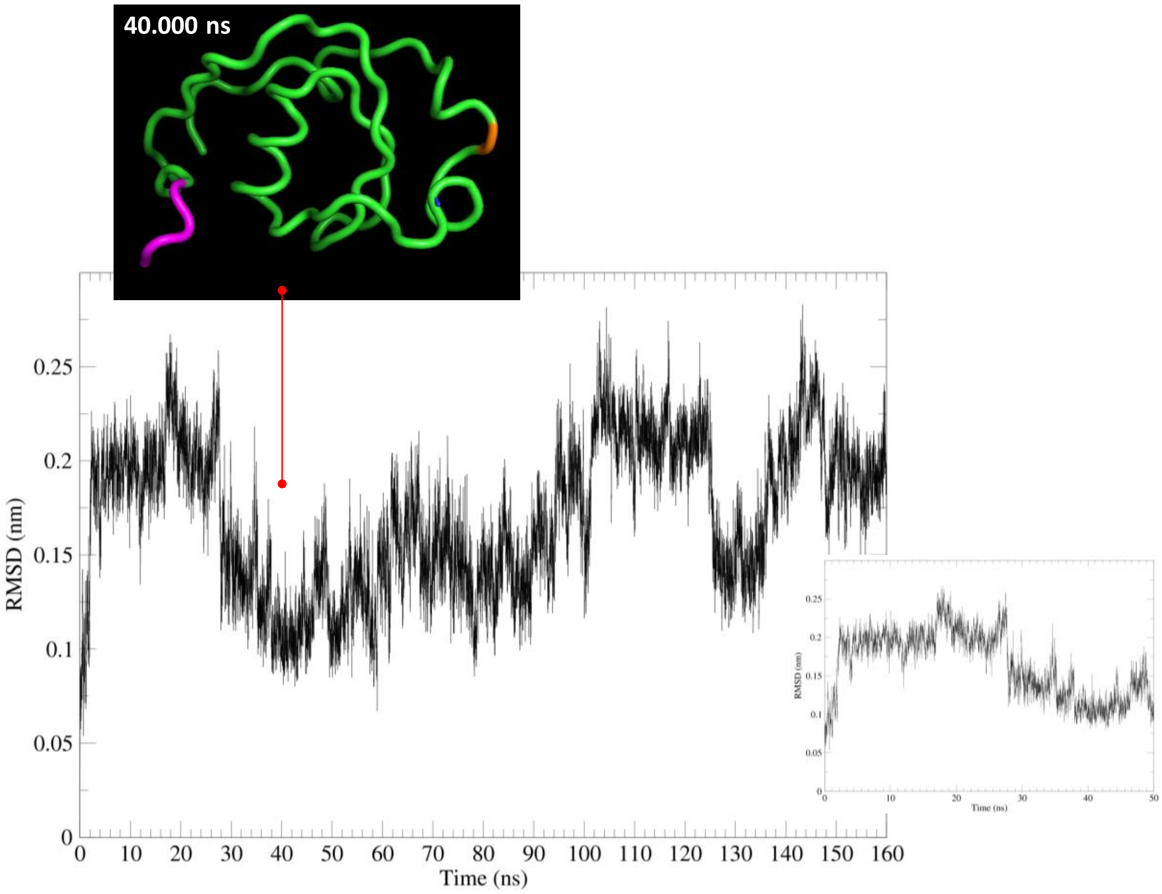

**Supplementary Figure 36.** Time evolution of the root-mean-square deviation (RMSD) of BgSmin$^{P}$ **D88E** backbone atoms for trajectory 1. Temperature: 300 K

**40.000 ns**

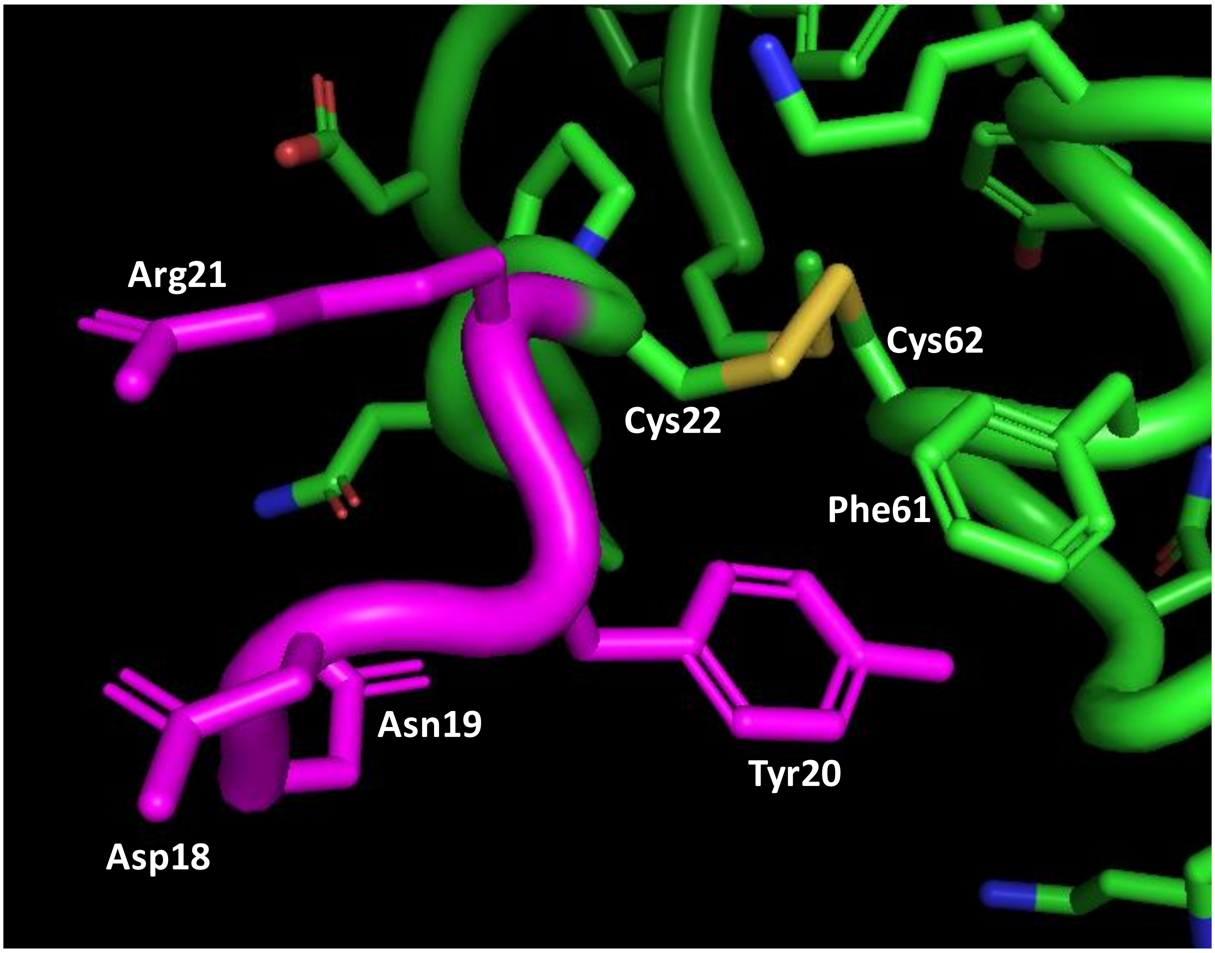

**Supplementary Figure 37.** Detailed view of the N-terminal region of BgSmin$^{P}$ (D88E) at 40 ns (trajectory 1). Temperature: 300 K

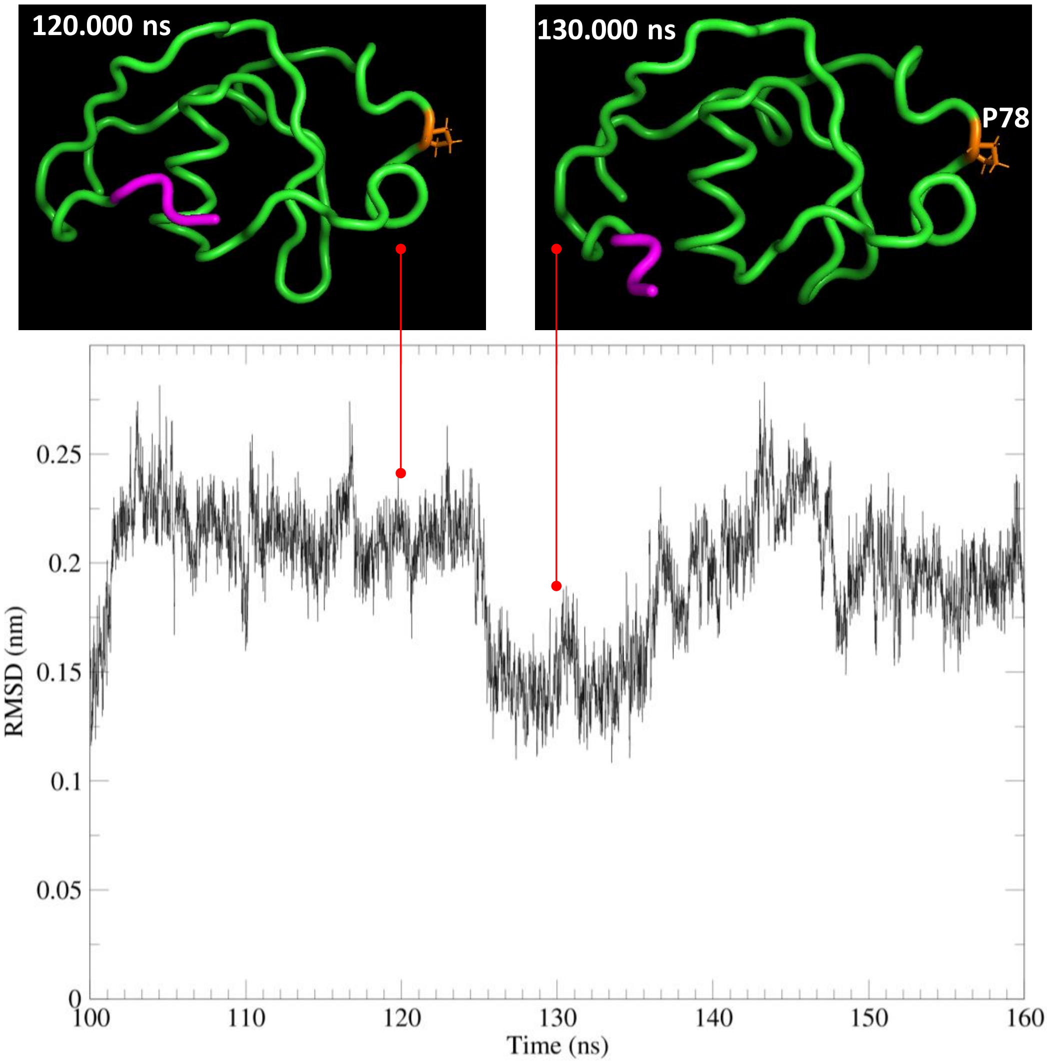

**Supplementary Figure 38.** Time evolution of the root-mean-square deviation (RMSD) of BgSmin[P] **D88E** backbone atoms for trajectory 1 between 100 and 160 ns. Temperature: 300 K

**120.000 ns**

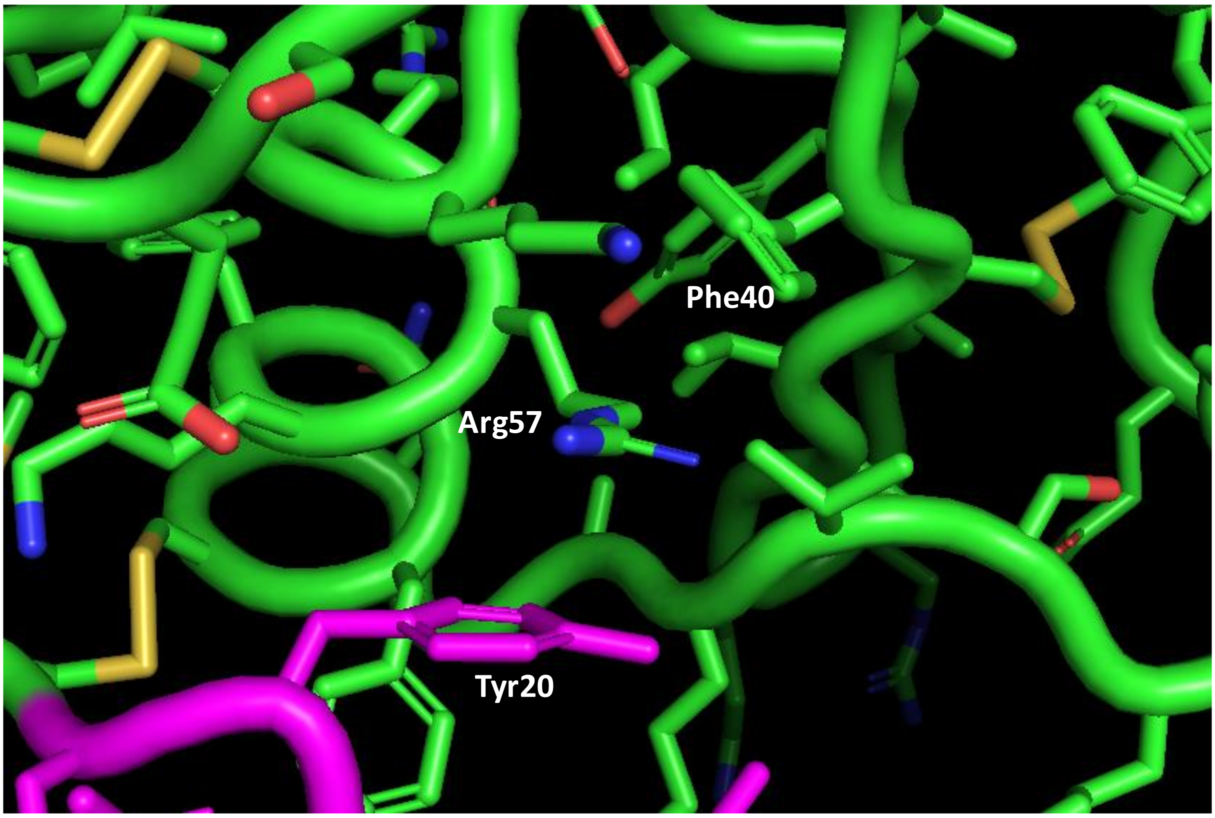

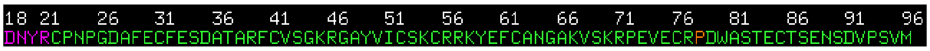

**Supplementary Figure 39.** Detailed view of the N-terminal region of BgSmin$^{P}$ (D88E) at 120 ns (trajectory 1). Temperature: 300 K

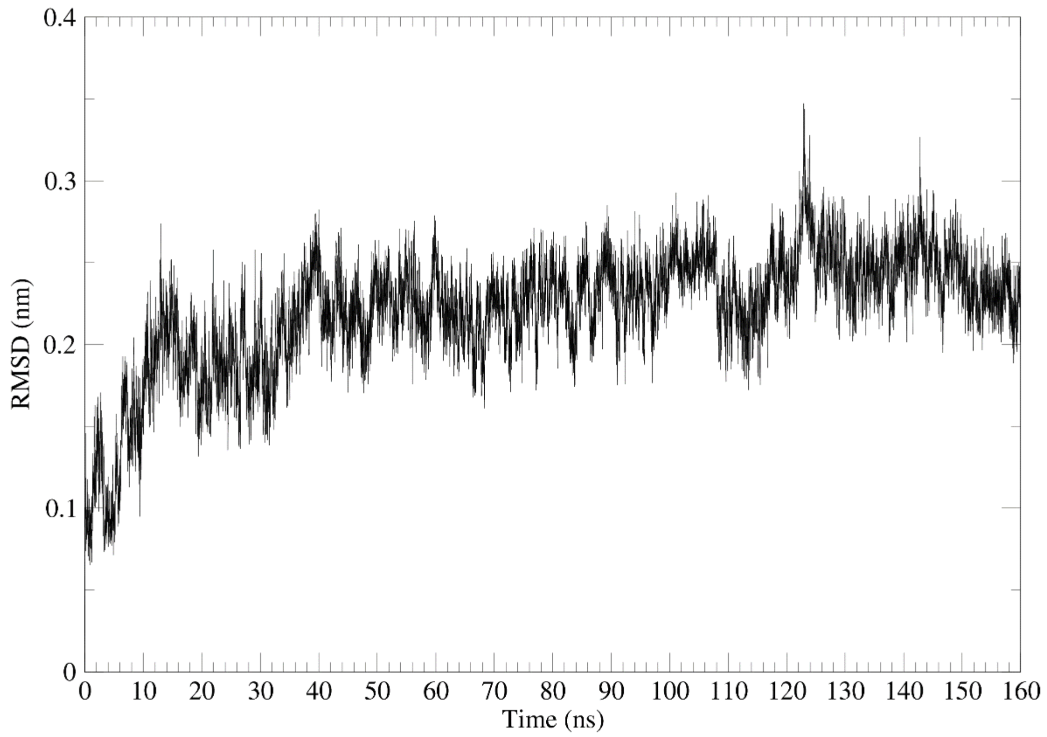

**Supplementary Figure 40.** Time evolution of the root-mean-square deviation (RMSD) of BgSmin[P] **D88E** backbone atoms for trajectory 2. Temperature: 300 K

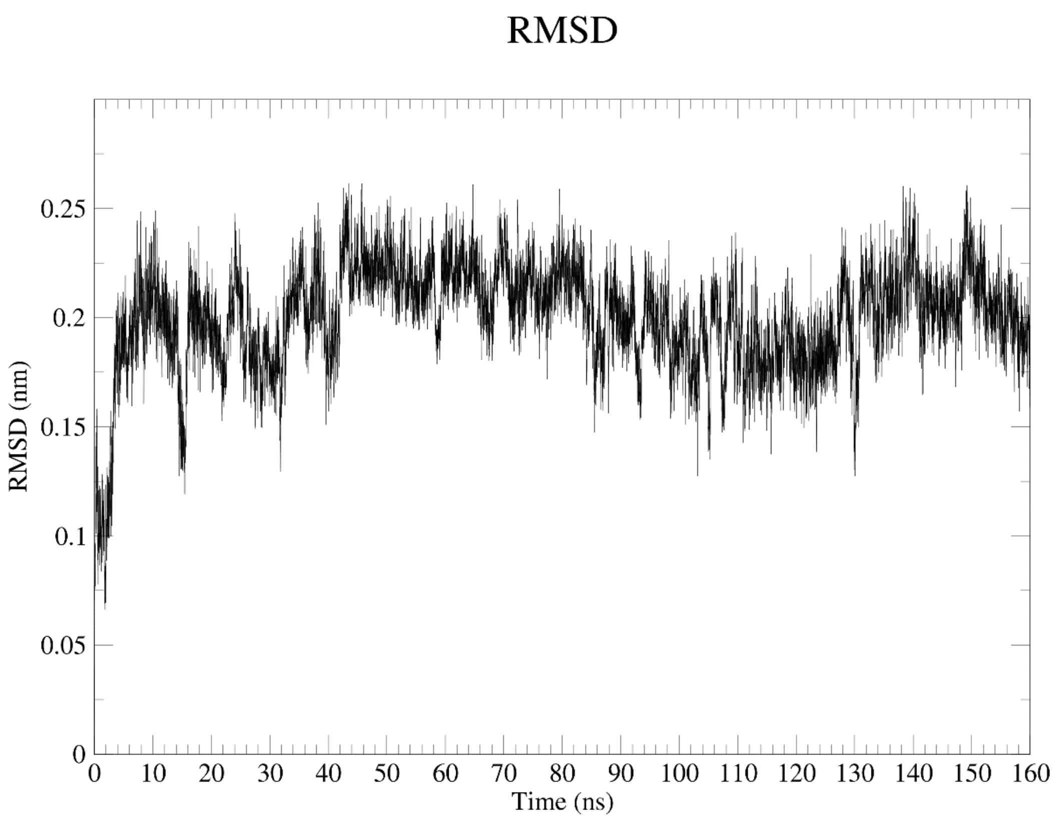

**Supplementary Figure 41.** Time evolution of the root-mean-square deviation (RMSD) of $BgSmin^{P}$ **D88E** backbone atoms for trajectory 3. Temperature: 300 K

**7.4.2** Radius of gyration

A)

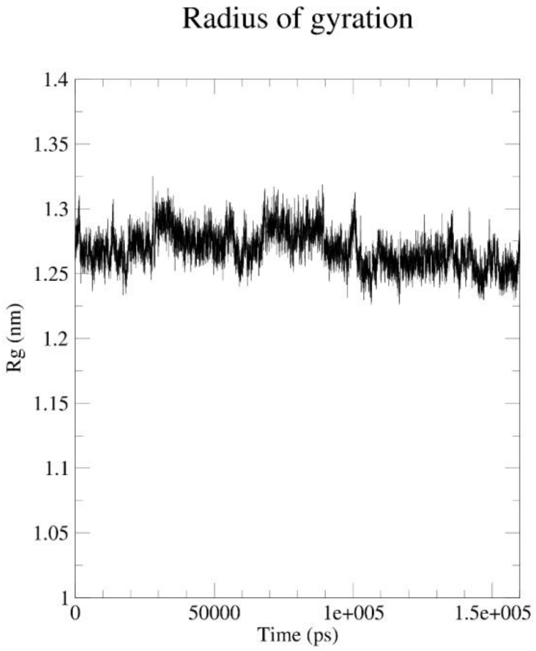

Radius of gyration
Rg (nm)
Time (ps)
1.4
1.35
1.3
1.25
1.2
1.15
1.1
1.05
1
0
50000
1e+005
1.5e+005

B)

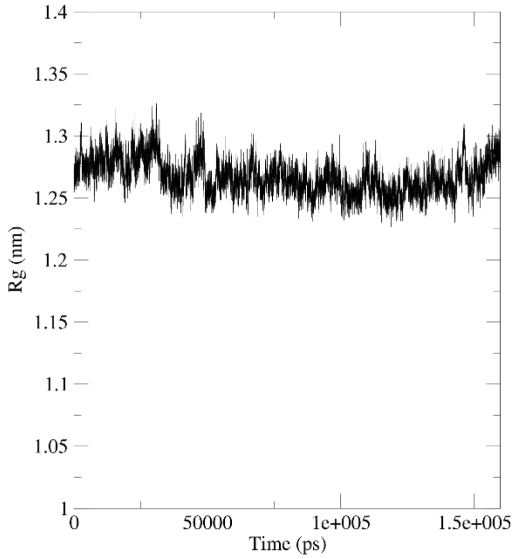

Rg (nm)
Time (ps)
1.4
1.35
1.3
1.25
1.2
1.15
1.1
1.05
1
0
50000
1e+005
1.5e+005

C)

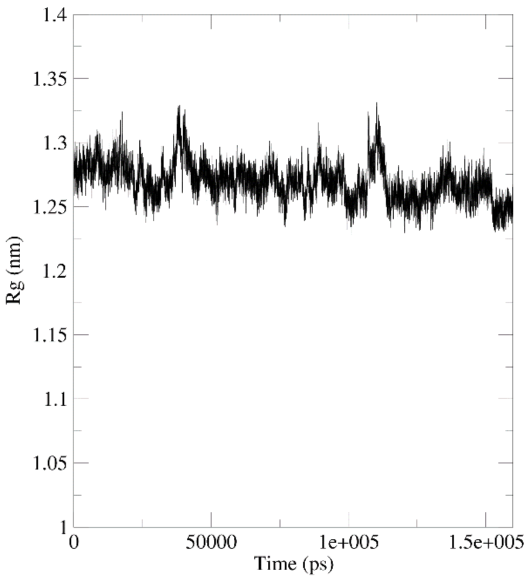

**Supplementary Figure 42.** Time evolution of the radius of gyration of BgSmin$^P$ D88E during trajectory 1 (A), trajectory 2 (B) and trajectory 3 (C). Temperature: 300 K

##### 7.4.3 Root-mean-square fluctuation (RMSF)

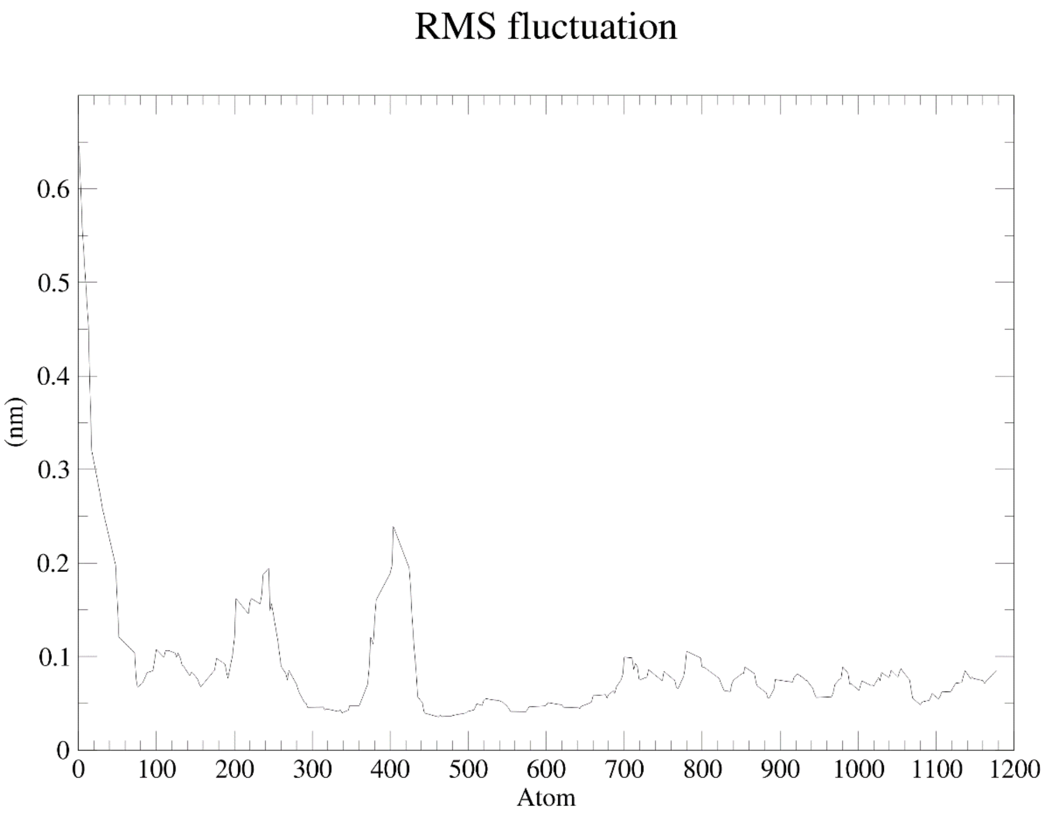

**Supplementary Figure 43.** Root-mean-square fluctuation of BgSmin$^P$ D88E backbone atoms for trajectory 1 (2-160 ns). Temperature: 300 K

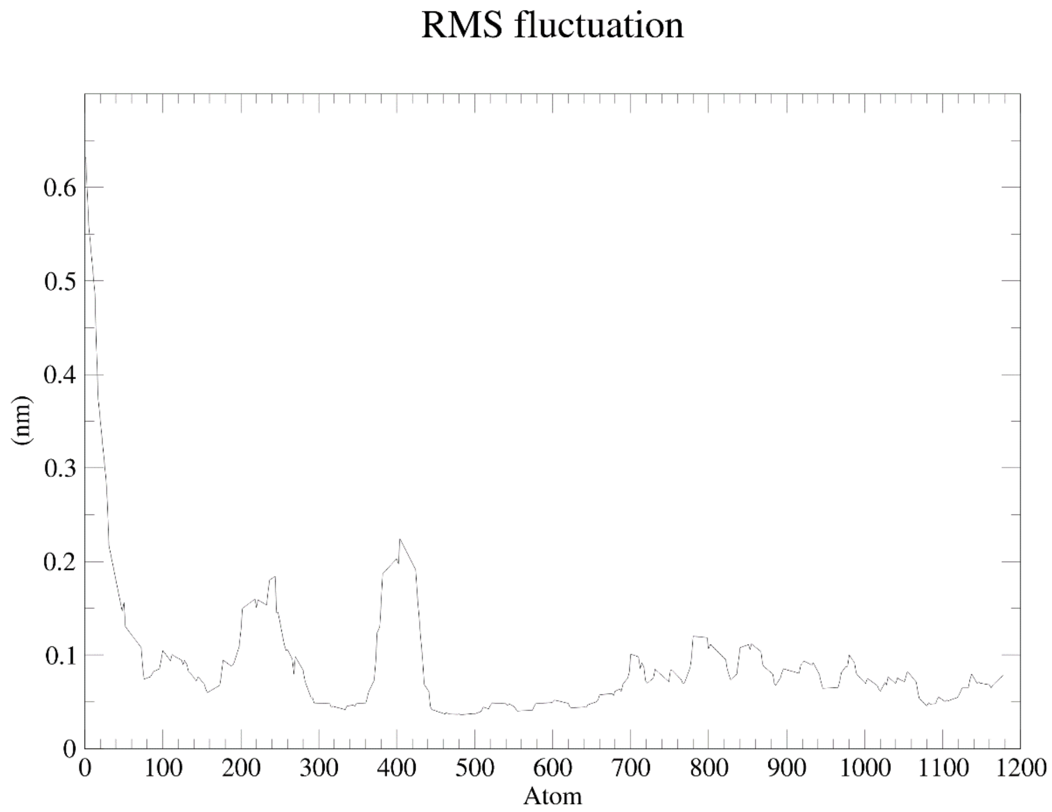

**Supplementary Figure 44.** Root-mean-square fluctuation of BgSmin$^{P}$ D88E backbone atoms for trajectory 2 (15-160 ns). Temperature: 300 K

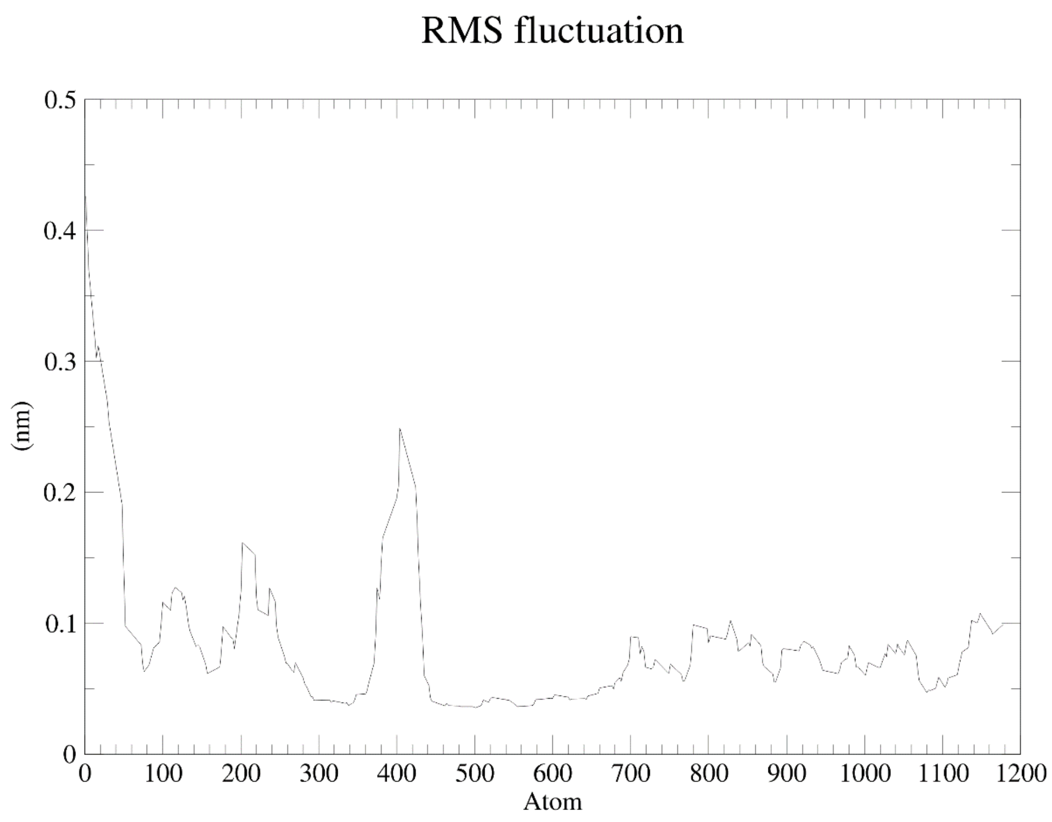

**Supplementary Figure 45.** Root-mean-square fluctuation of BgSmin$^{P}$ D88E backbone atoms for trajectory 3 (10-160 ns). Temperature: 300 K

**7.4.4** Ramachandran plots

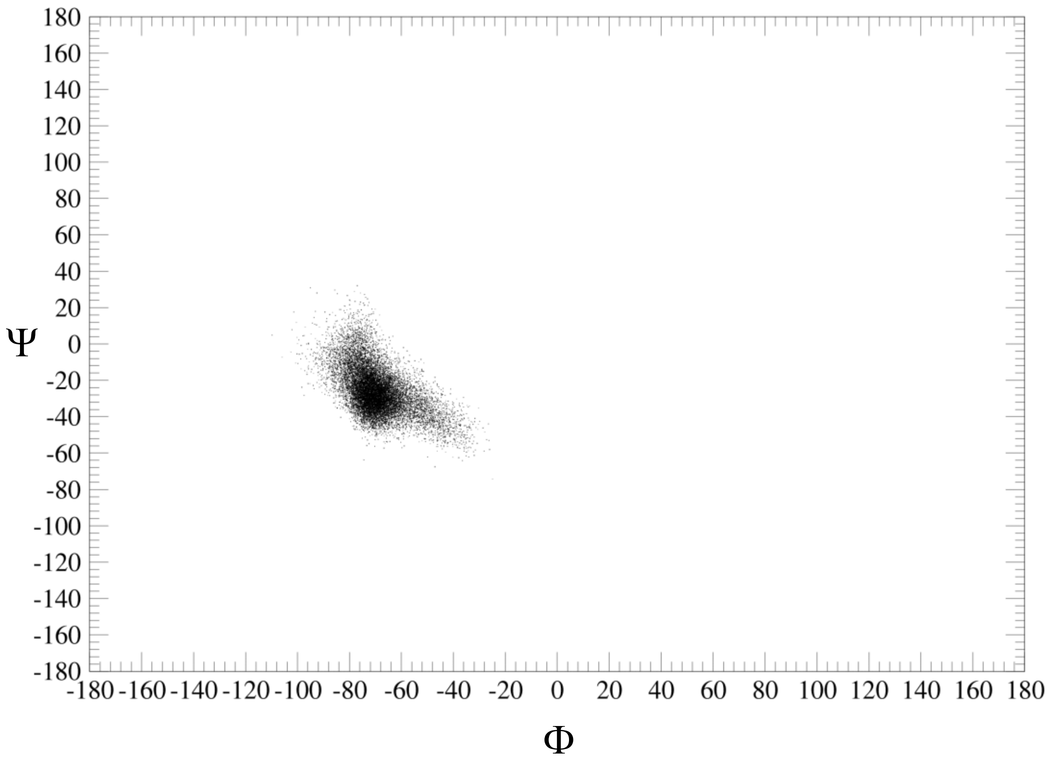

**Supplementary Figure 46.** Ramachandran plot for Pro78 over trajectory 1, 300 K.

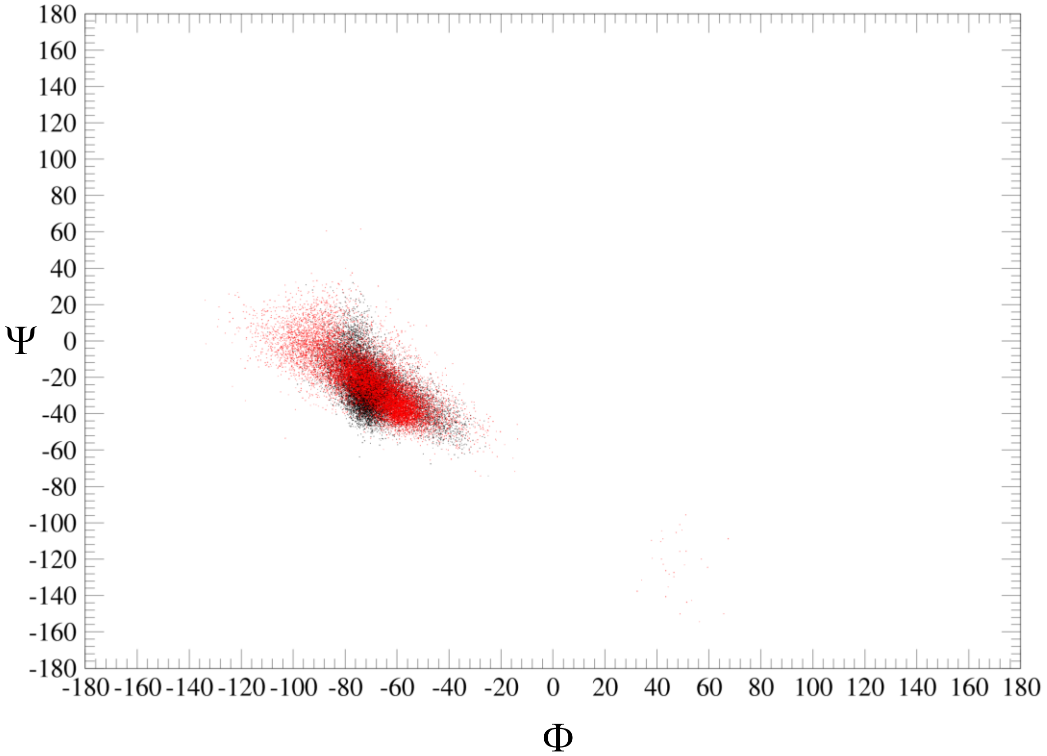

**Supplementary Figure 47.** Overlap of Ramachandran plot for Ala78 (red dots, BgSmin$^{A}$ D88E) and Pro78 (black dots, BgSmin$^{P}$ D88E) over trajectories 1, 300 K.

#### 8. Isolation of schistosomins from *Biomphalaria glabrata* extracts

##### 8.1 Snail maintenance and dissection

*Biomphalaria glabrata* (Puerto Rico strain) were maintained under controlled laboratory conditions in aerated aquaria containing deionized water at 26 °C. Snails were fed ad libitum with fresh spinach and a mixture of spirulina, brewer's yeast, skimmed milk, and wheat germ, under a 12 h light/12 h dark cycle. Animals were starved for 24 h prior to experiments.

##### 8.2 Isolation of native schistosomins

Thirty *Biomphalaria glabrata* snails (17–20 mm shell diameter) were anesthetized in chilled 50 mM $MgCl_2$, and the feet containing the neuronal ganglia were dissected. Organs were immediately frozen in liquid nitrogen and ground into a fine powder with a pre-chilled mortar and pestle. The tissue powder was suspended in 12 mL of ultrapure water and sonicated (30 cycles of 30 s on/off) at 4°C using a Bioruptor Plus (Diagenode) with integrated cooling.

The homogenate was centrifuged at 14,000 rpm for 20 min at 4 °C. Supernatants were collected, split into two tubes (6 mL), and subjected to ammonium sulfate precipitation as described elsewhere.[15]

Proteins were first precipitated at 60% ammonium sulfate saturation for 30 min at room temperature, followed by centrifugation at 15,000 g for 30 min at 4 °C. The resulting supernatant was further precipitated at 80% ammonium sulfate saturation, and pellets were collected by centrifugation.

Final protein pellets were resuspended in 500 µL of 10 mM HEPES buffer (pH 7.5) and dialyzed (3.5 kDa MWCO, Slide-A-Lyzer™ cassettes, Thermo Fisher Scientific) against 500 mL of 10 mM HEPES buffer at 4 °C, with three buffer changes over 6 h. Samples were stored at −20 °C until further use. Protein concentration was determined by BCA assay (ThermoFisher Scientific, #A55865)prior to analysis and separation.

##### 8.3 Analysis of Bg extracts by UPLC-MS

UPLC-MS analysis of Bg protein extracts was performed using the following experimental conditions:

- Thermo Scentific Dionex ultimate 3000 (HPLC) linked with LCQ Fleet (MS)
- Column: ACQUITY UPLC Peptide BEH C18, 300 Å, 1.7 µm, 2.1 mm x 150 mm
- Gradient 0 – 70 % B in 15 min. Eluents: A buffer: Water + 0.1 % v/v TFA. B buffer: Acetonitrile + 0.1 % v/v TFA
- 70 °C, 0.4 mL/min

A)

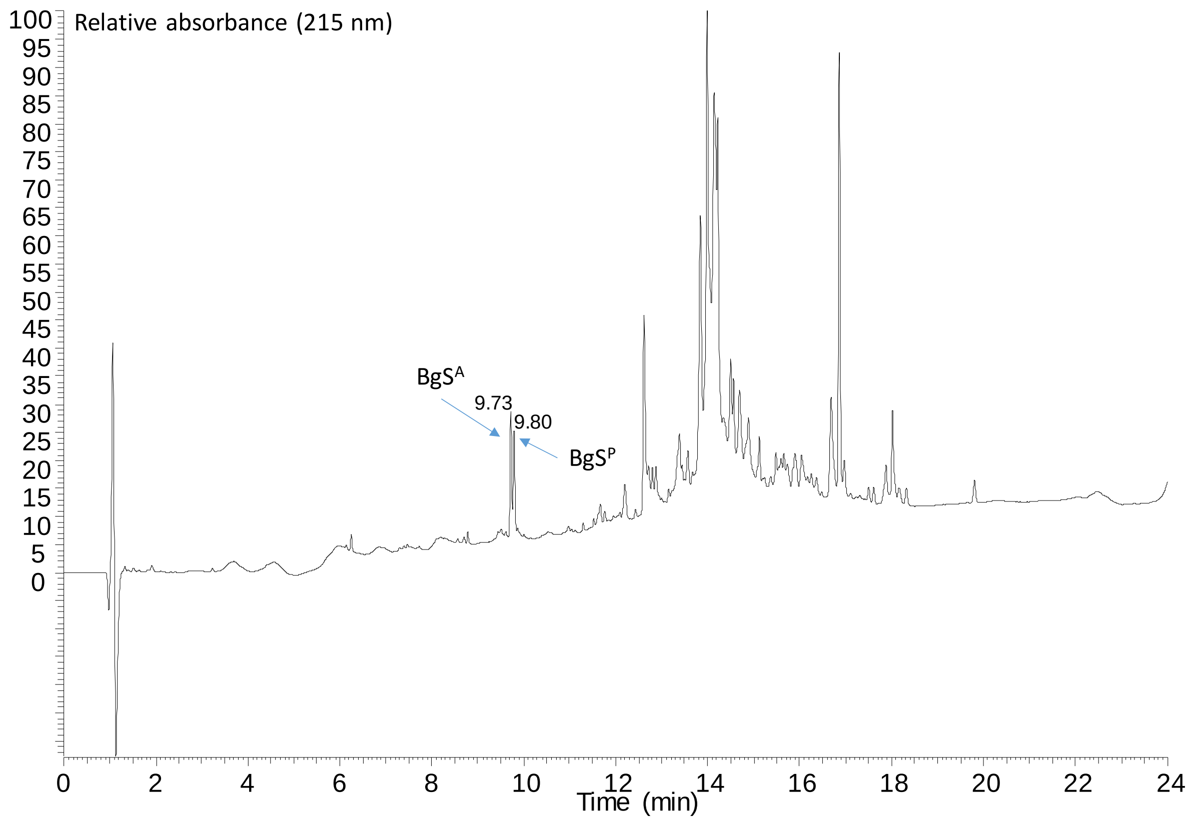

B)

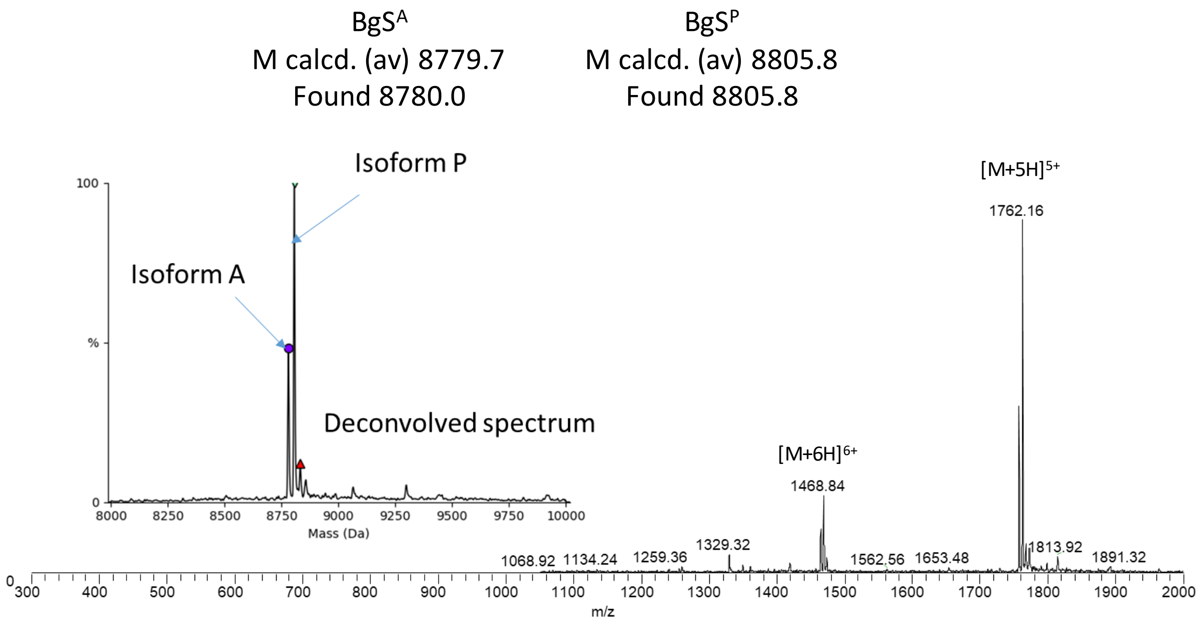

**Supplementary Figure 48.** UPLC analysis of Bg extract. A) LC trace. B) Trace.

##### 8.4 Analysis of purified BgSminA and BgSminP proteins by UPLC-MS

The UPLC analysis of purified BgSmin proteins has been performed using the following experimental conditions :

HPLC sytem: Gilson PLC-2020

Colomn : xBridge Peptide BEH C18 OBD Prep Column, 300 Å, 5 µm, 10 mm × 250 mm

Gradient: 0-10% B (5 min), 15-40% B (60 min). A buffer: Water + 0.1 % v/v TFA. B buffer: Acetonitrile + 0.1 % v/v TFA

5 mL/min, temperature : 50 °C, detection at 215 nm

2 µL of a protein solution at 1 mg/mL was injected for the analysis

A)

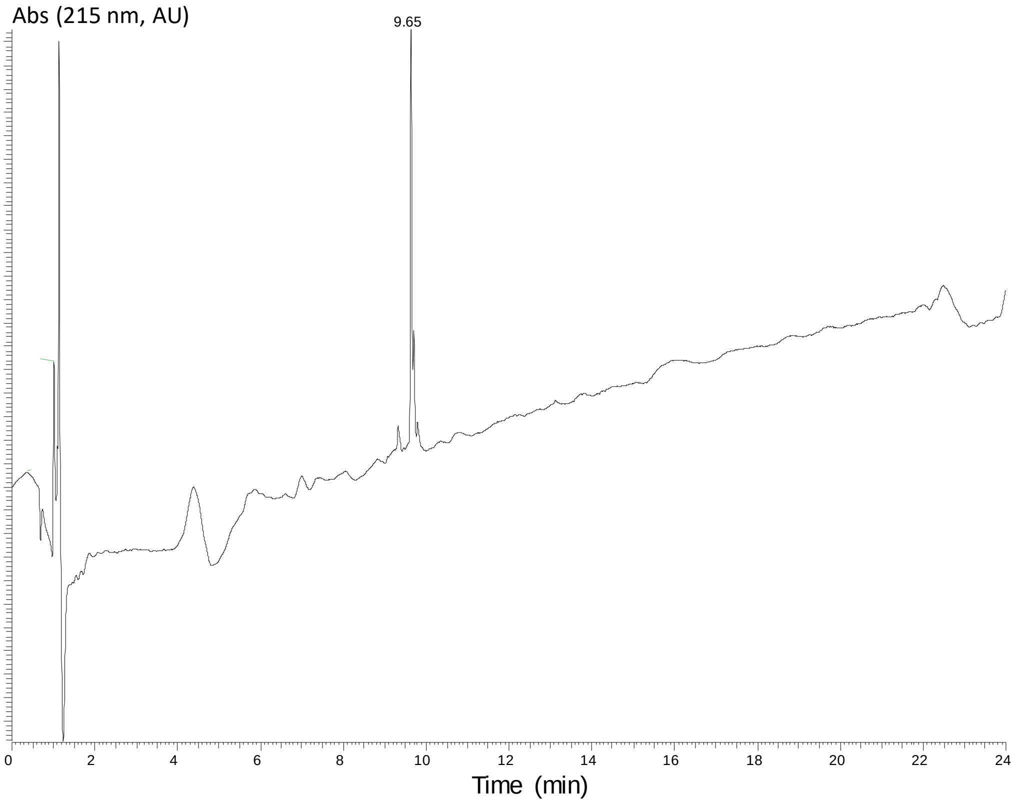

B)

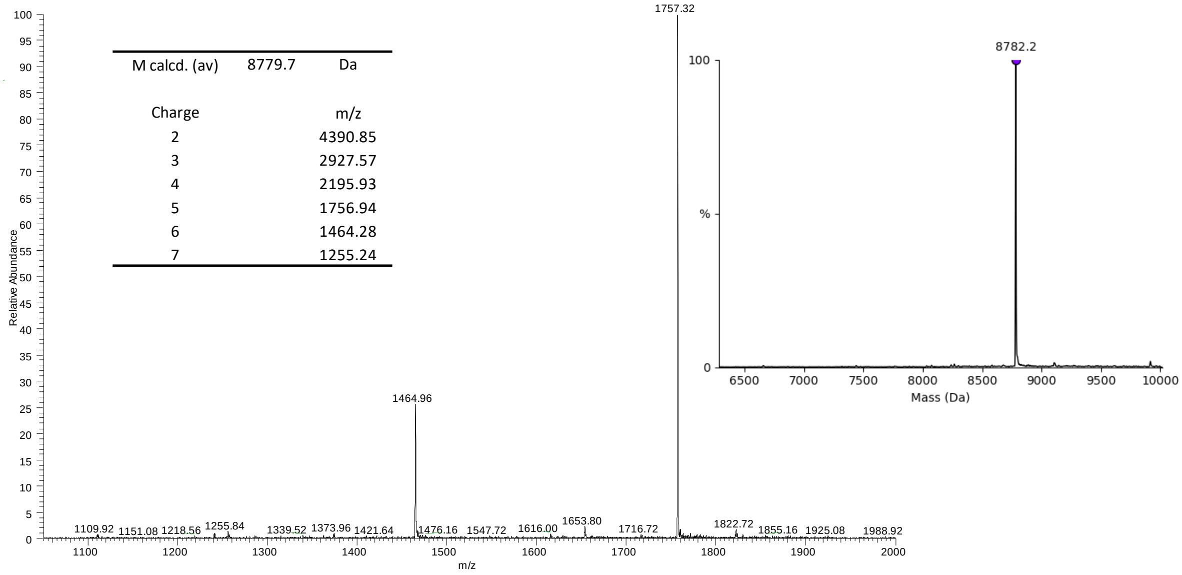

**Supplementary Figure 49.** UPLC analysis of BgSmin$^A$ extract. A) LC trace. B) Trace.

A)

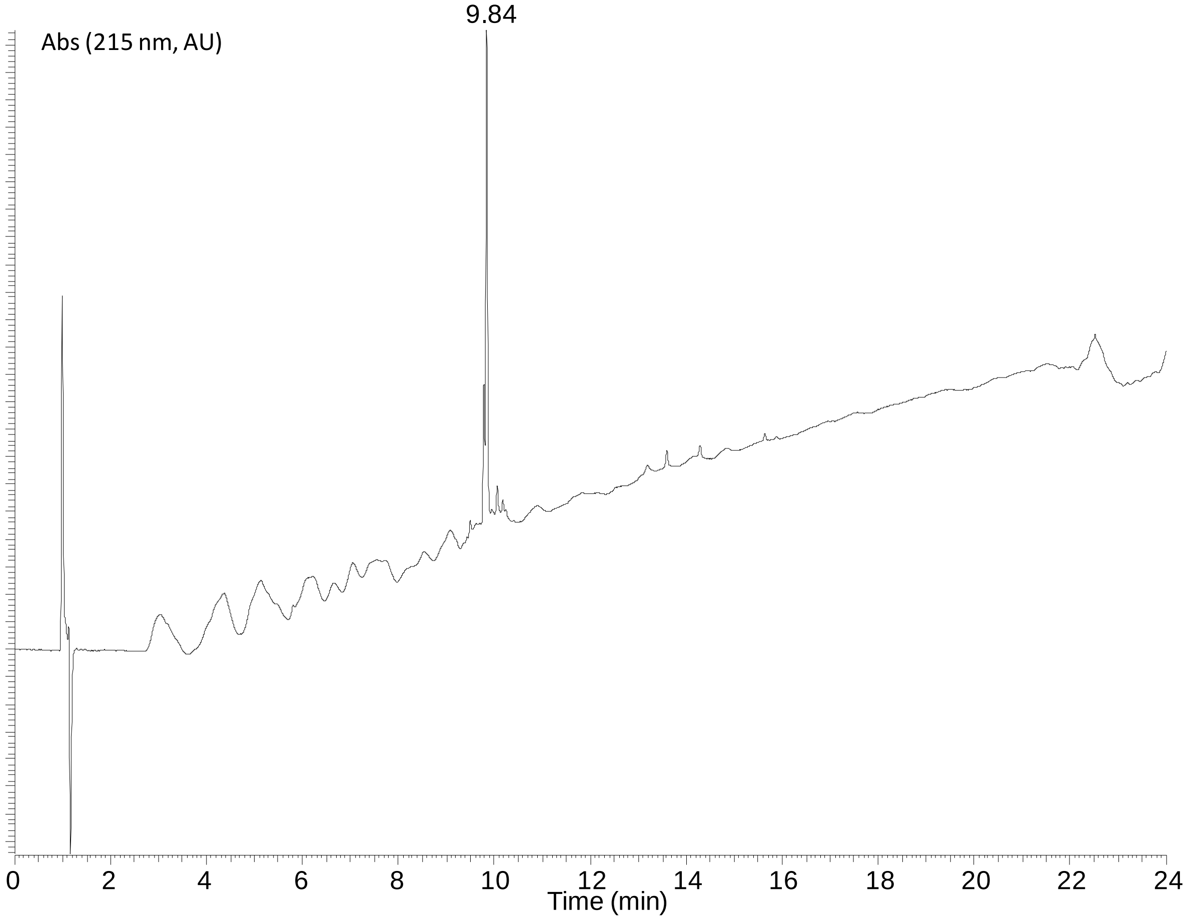

B)

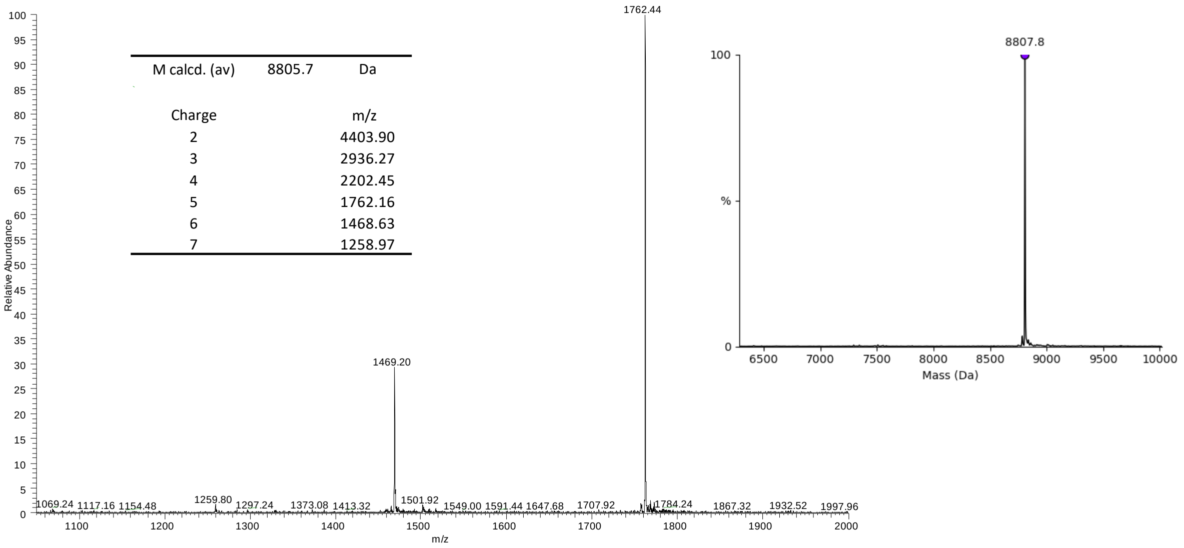

**Supplementary Figure 50.** UPLC analysis of BgSmin$^{P}$ extract. A) LC trace. B) Trace.

#### 8.5 HRMS analysis of BgSminmin isolated from Bg

##### 8.5.1 A isoform

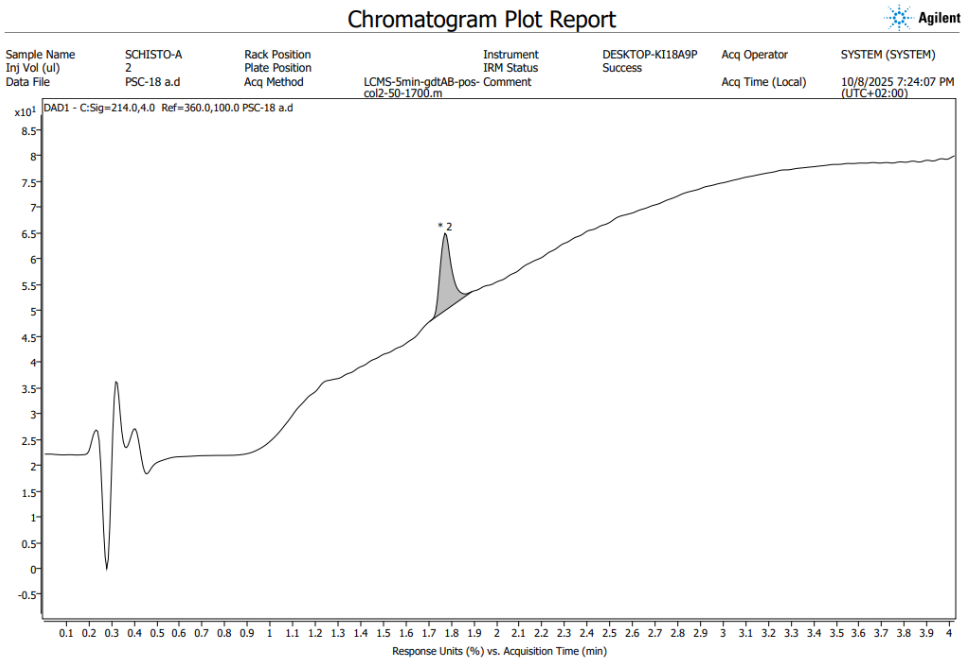

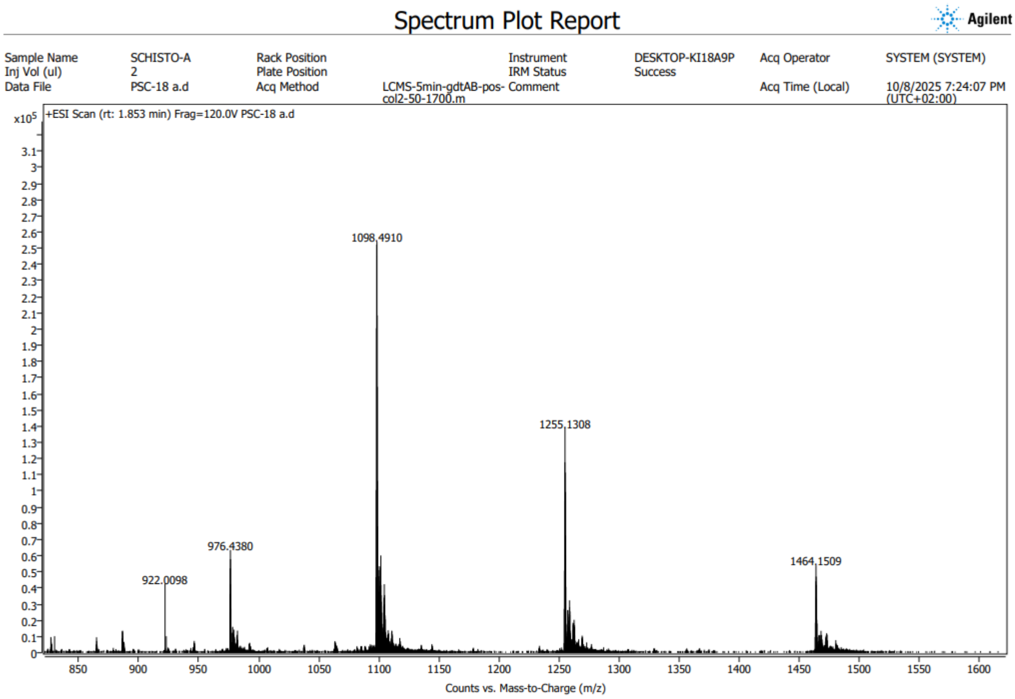

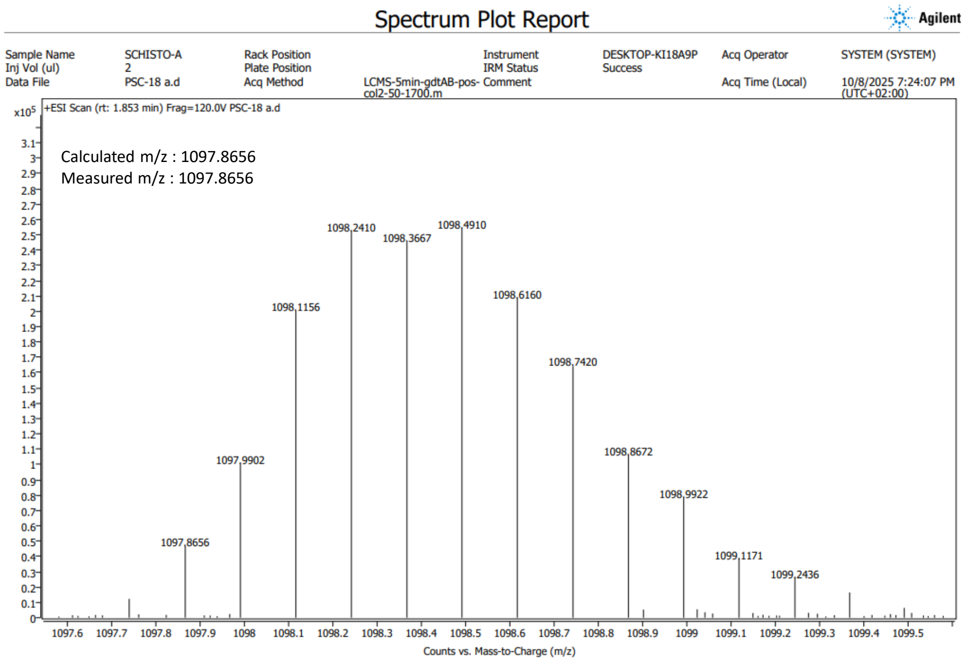

**Supplementary Figure 51**. HRMS analysis of BgSminmin[A]. HRMS data were aquired using An Orbitrap ID-X (Thermo) with ESI source coupled with an UPLC Vanquish (Thermo), with a Kinetex EVO C18 50 x 2.1 mm 1.7 µm column (Phenomenex). UV-Vis chromatograms were acquired on a 200-400 nm range. Mass spectra were acquired in a 100-2000 Da range in positive mode with a 120.000 resolution. ESI, *m/z* for $[C_{369}H_{570}N_{110}O_{122}\ S_{10}]^{8+}$ calculated m/z for A+1 ion: 1097.8656, measured: 1097.8656.

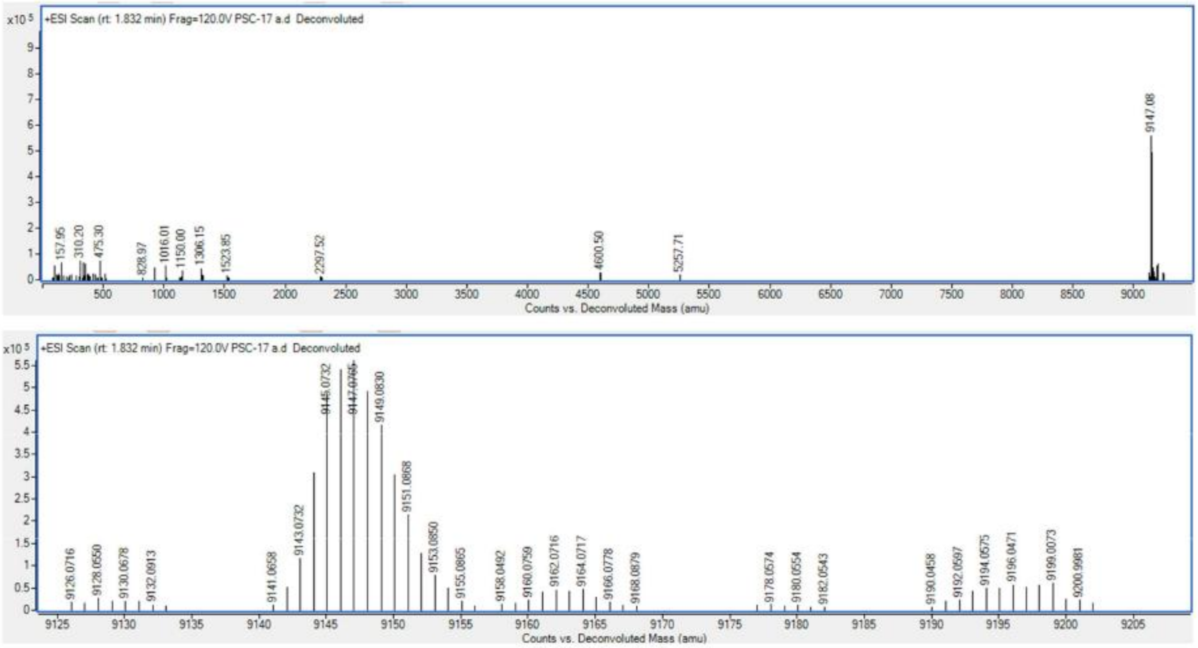

**Supplementary Figure 52.** HRMS deconvoluted spectrum for BgSminmin$^{A}$.

##### 8.5.2 P isoform

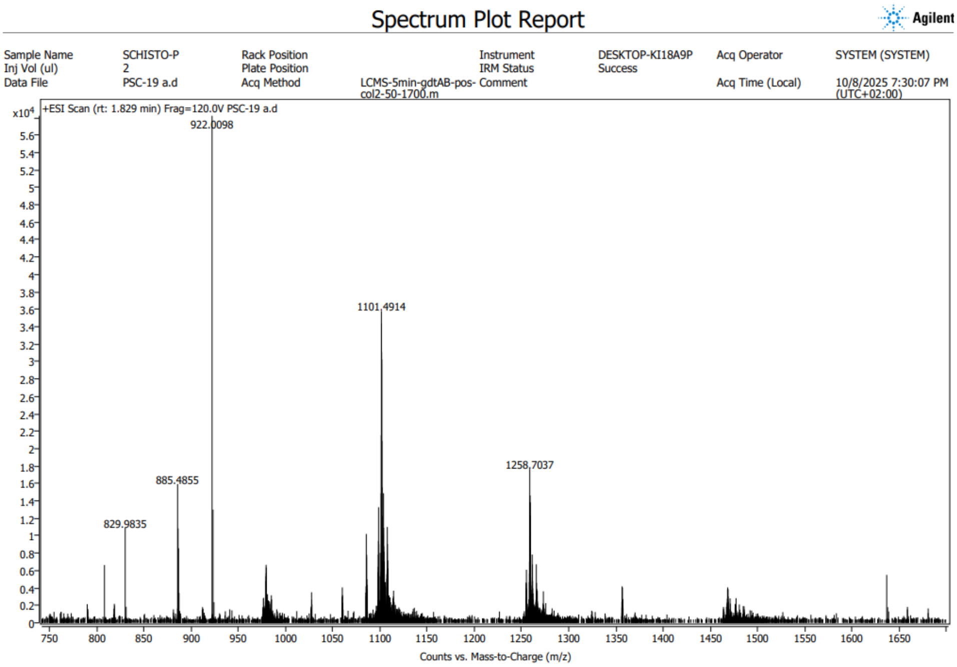

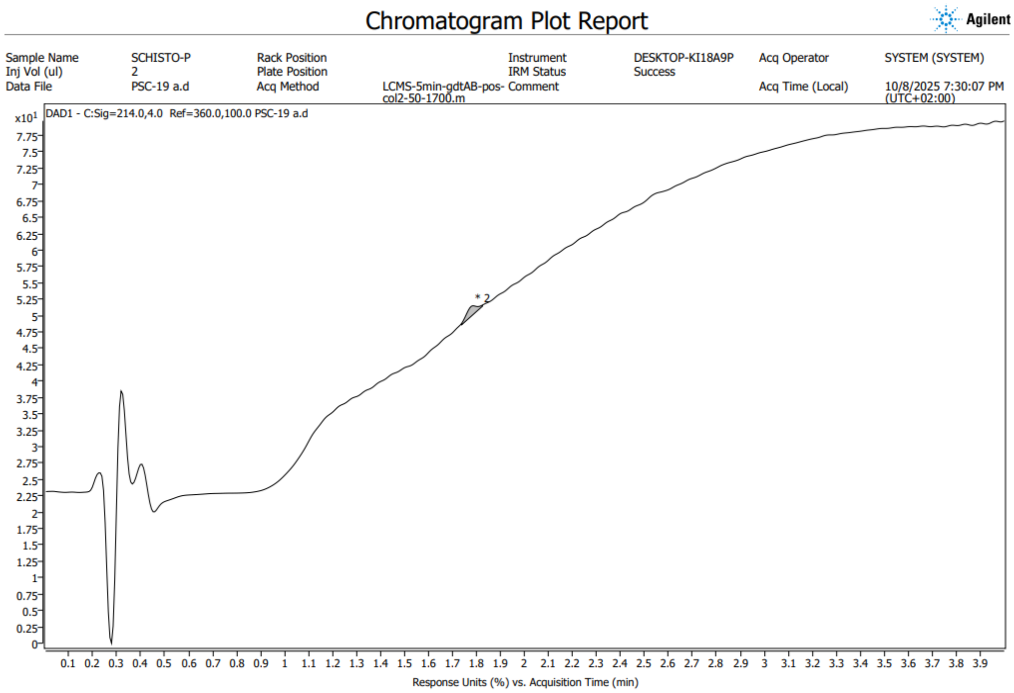

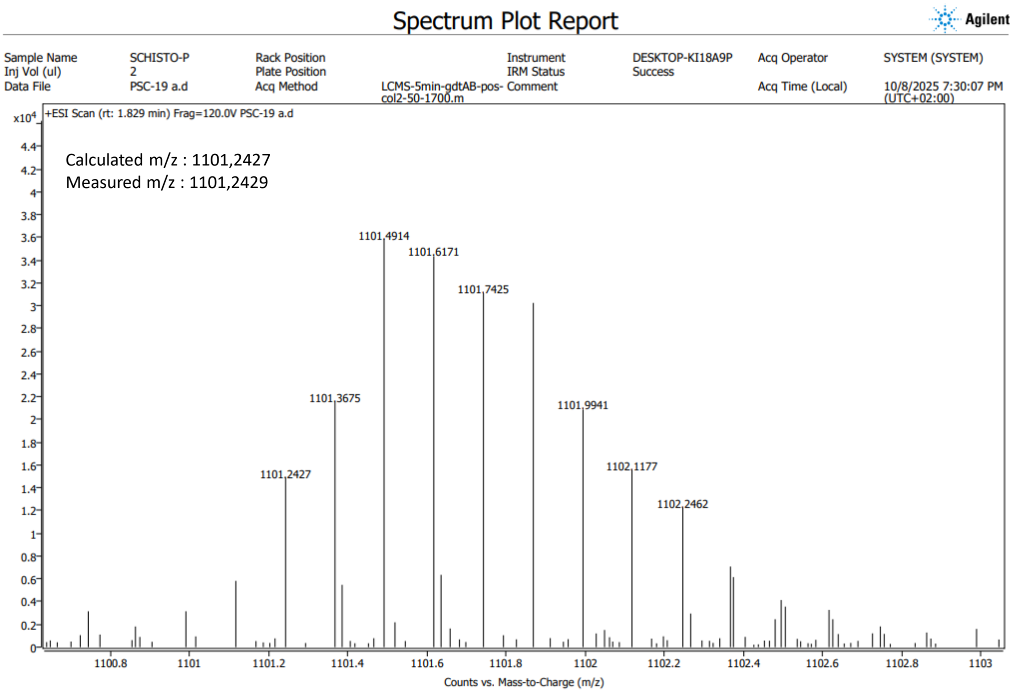

**Supplementary Figure 53.** HRMS analysis of BgSmin$^{P}$. HRMS data were aquired using An Orbitrap ID-X (Thermo) with ESI source coupled with an UPLC Vanquish (Thermo), with a Kinetex EVO C18 50 x 2.1 mm 1.7 µm column (Phenomenex). UV-Vis chromatograms were acquired on a 200-400 nm range. Mass spectra were acquired in a 100-2000 Da range in positive mode with a 120.000 resolution. ESI, *m/z* for $[C_{371}H_{564}N_{110}O_{122}\ S_{10}]^{8+}$ calculated m/z for A+2 ion: 1101.2427, measured: 1101.2429.

### 9 Phylogenic analysis

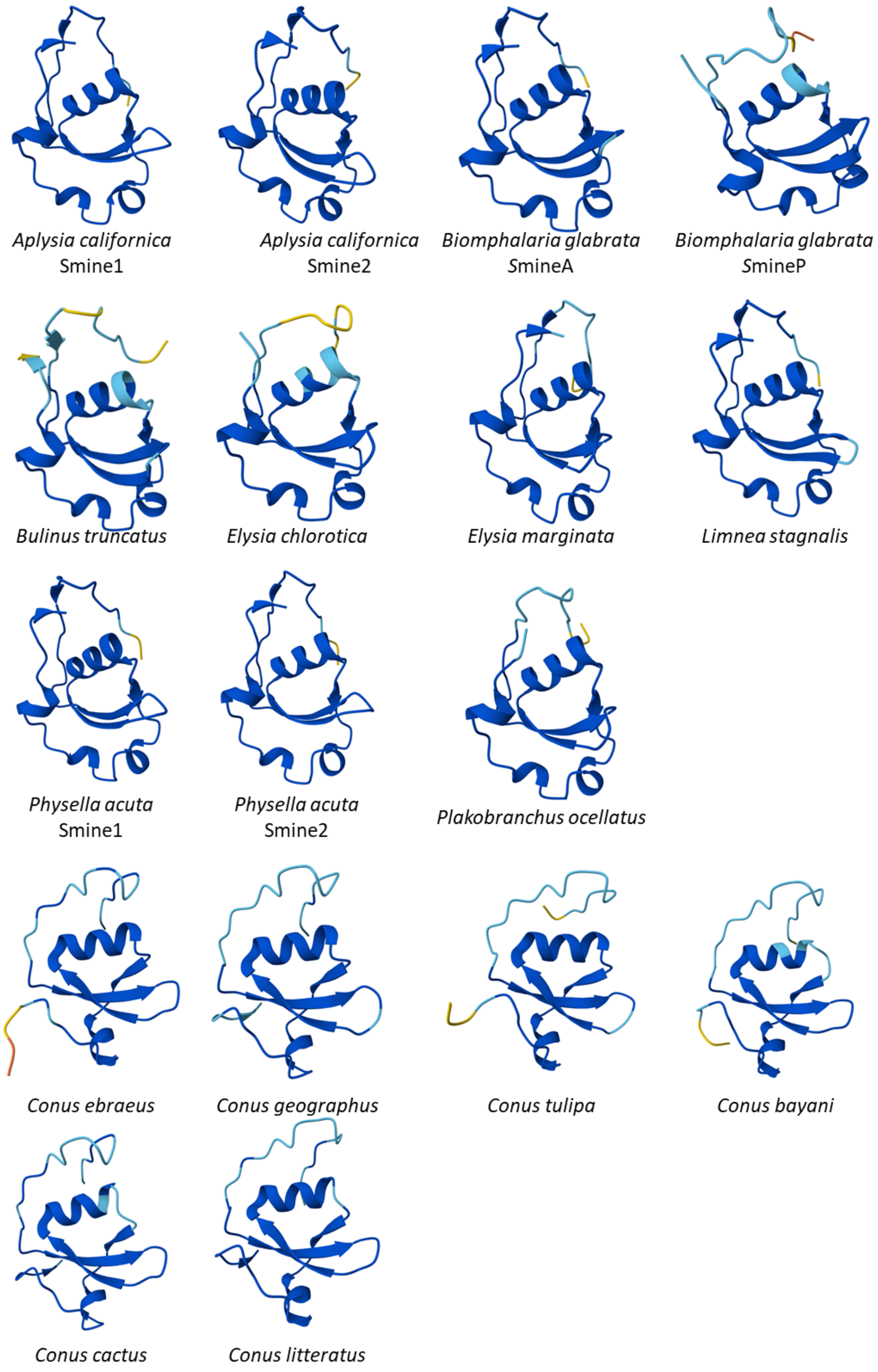

**Supplementary Figure 54**: AlphaFold-predicted structures of putative schistosomin-related peptides from various *Heterobranchia* gastropods and *Conus* peptides

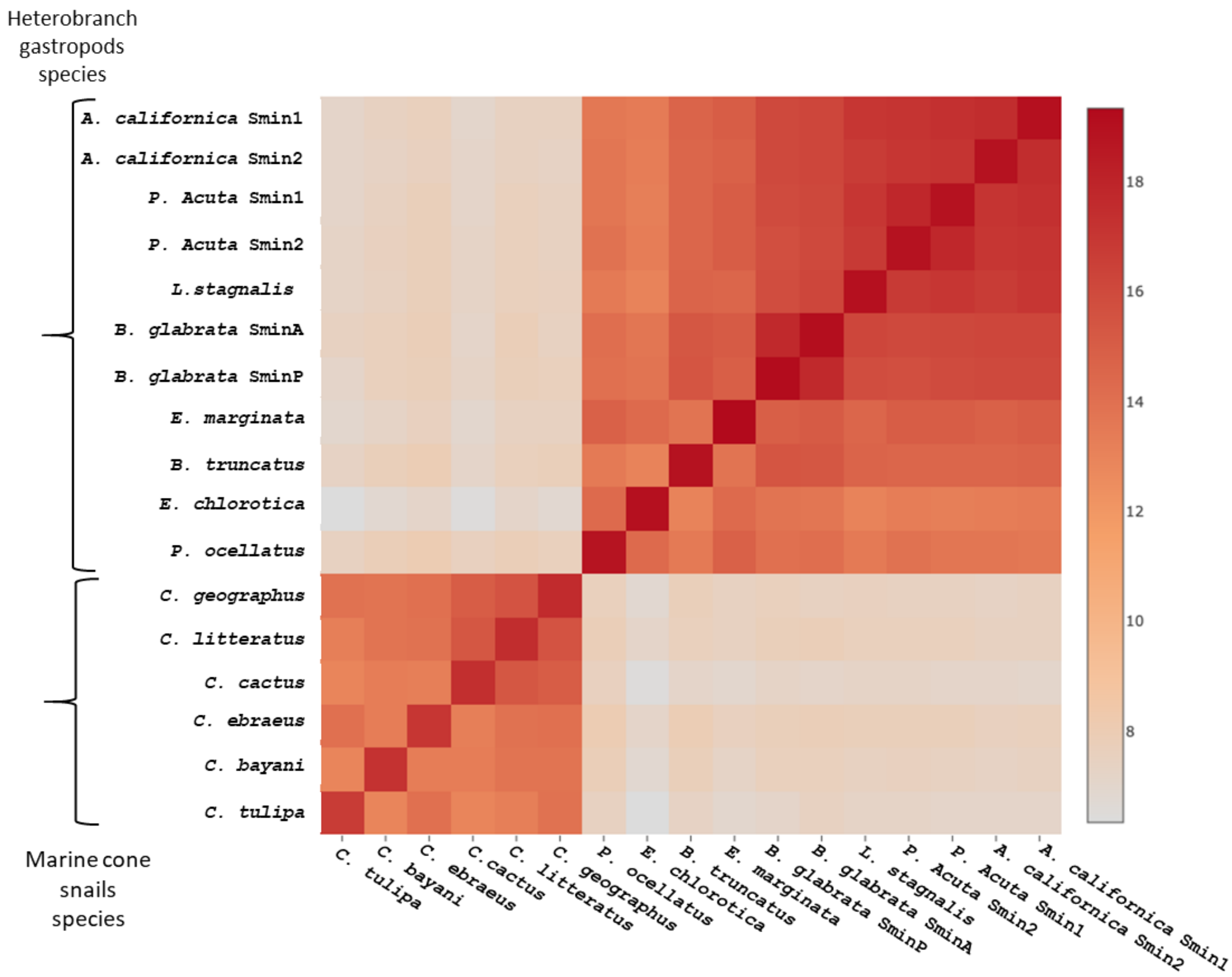

**Supplementary Figure 55**: Heatmap of pairwise structural similarity between putative schistosomin-related peptides. Heatmap was generated based on Z-scores obtained with the DALI server. Higher Z-scores indicate higher structural similarity and conservation.

#### 9. Localization of BgSmin proteins in the snail

##### 9.1 Snail dissection

*Biomphalaria glabrata* (Puerto Rico strain) were maintained under controlled laboratory conditions in aerated aquaria containing deionized water at 26 °C. Snails were fed ad libitum with fresh spinach and a mixture of spirulina, brewer's yeast, skimmed milk, and wheat germ, under a 12 h light/12 h dark cycle. Animals were starved for 24 h prior to experiments.

Sexually mature adult snails (shell diameter 17–20 mm, confirmed by egg-laying capacity) were selected for dissections. Prior to dissection, snails were anesthetized for 10–20 min in chilled 50 mM $MgCl_2$. Soft tissues were carefully removed from the shell using round-tipped scissors, rinsed in snail phosphate-buffered saline (sPBS; 8.41 mM $Na_2HPO_4$, 1.65 mM $NaH_2PO_4 \cdot H_2O$, 45.34 mM NaCl, pH 7.2), and transferred to a Sylgard-coated Petri dish (Dow-Corning, Midland, MI, USA) filled with fresh sPBS. Dissections were performed under a Zeiss stereomicroscope.

The following tissues were collected: tentacles, mantle collar, hepatopancreas, gonads, intestine, kidneys, rectal crest, foot, heart, stomach, radula, and 11 cerebral and peripheral ganglia. Immediately after dissection, tissues were pooled in groups of three and transferred into Lysing Matrix D tubes (MP Biomedicals) containing 1 mL of TRIzol reagent (Thermo Fisher Scientific). RNA and proteins were extracted according to the manufacturer's instructions. RNA concentration and purity were assessed by spectrophotometry using a NanoDrop system (Thermo Fisher Scientific). All dissections were performed in triplicate.

##### 9.2 Hemolymph and hemocyte collection

Hemolymph was collected from 80 snails (shell diameter 15-20 mm) using the head-foot retraction method. After drying the inner surface of the shell, hemolymph was withdrawn by piercing the foot of unanesthetized snails with 200 µL round gel-loading tips (Starlab). Hemolymph from 10 individuals was pooled per sample and transferred into wells of a 12-well culture plate (Falcon), which was incubated at 26 °C for 3 h to allow hemocyte adhesion.

The hemocyte-free supernatant was collected and mixed with TRIzol LS reagent (Thermo Fisher Scientific) for protein extraction. Wells containing adherent hemocytes were washed three times with sPBS, and RNA/protein extraction was performed using the NucleoSpin RNA/Protein kit (Macherey-Nagel). RNA concentration and integrity were verified by spectrophotometry on a NanoDrop system (Thermo Fisher Scientific).

##### 9.3 RT-qPCR quantification

Total RNA extracted from mollusc tissues was reverse-transcribed using the AffinityScript Multi Temperature cDNA Synthesis kit (Agilent, #200436), following the manufacturer's instructions. For each reaction, 500 ng of total RNA were used. Quantitative PCR (qPCR) was performed using the Brilliant III Ultra-Fast SYBR Green QPCR Master Mix (Agilent, #600882) in a final volume of 20 µL in 96-well optical plates (Axygen, PCR-96-FLT-C). Each reaction contained 250 nM of each primer (sequences listed in Supplementary Table) and 1 µL of cDNA. Amplification was carried out on a QuantStudio 3 detection system (Applied Biosystems) with the following cycling parameters: 95 °C for 5 min; 40 cycles of 95 °C for 10 s, 60 °C for 20 s; followed by a melting curve analysis to determine amplicon Tm values.

Relative mRNA levels were normalized to actin as the reference gene and analyzed using the comparative threshold cycle method ($2^{-\Delta\Delta Ct}$). Statistical analyses were performed using a two-way ANOVA with a significance level set at $\alpha = 0.05$.

#### Supplementary Table 5. Primer sequences

| Target | Forward (5'→3') | Reverse (5'→3') |
|---|---|---|
| Smine A | CTGCCAGGTTCTGTGTGTCTGG | GACAGGGATGCCGTTTGTTG |
| Smine P | CTGCCAGGTTCTGTGTGTCTGG | GGTTGTGTCTTGGTTGTGTCTTTAC |
| Actin | CAGGTTTCGCTGGAGACGAT | GCTGTCCTTCTGACCCATACCA |

##### 9.4 Western blot analysis

Production of anti-Schistosomin antibodies

A synthetic Schistosomin linear precursor (HPLC-purified, TFA salt) was used to generate mouse polyclonal antiserum. Three female NMRI mice received subcutaneous injections of 75 µg of peptide emulsified in alum adjuvant (1:1, v/v; total volume 200 µL) on days 0, 21, and 31.
Blood samples were collected from the mandibular vein prior to each injection and on days 45 and 60. Mice were euthanized 35 days after the last injection, and terminal blood was collected by cardiac puncture. Serum was clarified by centrifugation (3,000 rpm, 10 min), aliquoted (20 µL), and stored at −20 °C until use.).
The specificity and titration of the anti-Schistosomin antibodies were assessed by Western blot and enzyme-linked immunosorbent assay (ELISA).

Protein concentrations were determined using the BCA Protein Assay Kit (Pierce). Equal amounts of protein (20 µg per sample) were separated on NuPAGE 4–12% Bis-Tris gels (Novex, Invitrogen) under reducing conditions and transferred for 1.5 h at 65 V onto 0.45 µm PVDF membranes (Immobilon-P®, Millipore®) in Towbin buffer (10% methanol, 1× Tris-glycine, 0.0025% SDS). Synthetic schistosomin $BgSmin^{A}_{Synth}$ was included for comparison of band intensities (5, 10, 30 ng).

Membranes were blocked for 1 h at room temperature in blocking buffer (8 g/L high-purity casein, 1× PBS, 0.2% Tween-20), then incubated overnight at 4 °C with anti-Schistosomin antibody (1:2000 dilution in PBS containing 5% protease-free BSA (Euromedex) and 0.1% sodium azide) under mild agitation.

After three washes (PBS with 0.05% Tween-20, 10 min each), membranes were incubated for 1 h at room temperature with HRP-conjugated anti-mouse secondary antibody (Jackson ImmunoResearch) diluted 1:50,000 in blocking buffer. Detection was performed using West Dura Extended Duration substrate (Thermo Scientific).

Chemiluminescence was recorded using the Amersham™ ImageQuant™ 800 system (Cytiva), and signal intensities were quantified with ImageQuant™ TL software. All experiments were performed in three independent replicates.
